# Towards CRISPR-based editing of the mitochondrial genome in yeast

**DOI:** 10.64898/2025.12.09.693232

**Authors:** Sifei Yin, Daniel F. Jarosz, Alice Y. Ting

## Abstract

Mitochondria, which evolved from symbiotic bacteria, possess their own genomes (mtDNA) and support independent transcription and translation within the organelle. Given the essential role of mtDNA in energy production, metabolism, as well as cellular homeostasis, and the high density of confirmed pathogenic mutations that map to mtDNA, there is a pressing need for versatile methods to study and manipulate this genome. Although CRISPR technology has revolutionized the editing of nuclear genomes, it has not been successfully extended to mtDNA, primarily due to the challenge of delivering single guide RNAs (sgRNAs) across both outer and inner mitochondrial membranes. Here we develop a survival-based reporter in *Saccharomyces cerevisiae* to screen for potential RNA import motifs. We identify a 40-nucleotide aptamer (IM83) that facilitates sgRNA entry into the mitochondrial matrix, enabling CRISPR editing by a mitochondrially-localized adenine base editor. We show that mitochondrial import of IM83 is ATP-dependent and enhanced by the tRNA synthetase Msk1. Further investigations identify barriers to efficient CRISPR editing of mtDNA, including loss of membrane potential associated with mitochondrial targeting of the base editor. These insights lay the groundwork for future improvements in CRISPR-based editing of mtDNA in eukaryotes.

## Introduction

Eukaryotic cells are constructed from two genomes: one nuclear (nDNA) and another mitochondrial (mtDNA) that resides within the mitochondrial matrix. Whereas CRISPR-based approaches have revolutionized the study of nDNA^1^, editing and other manipulations around mtDNA remain in their infancy. Yet the ability to edit, down-or up-regulate transcription, and image mtDNA in a locus-specific manner would open the door to many scientific investigations of mitochondrial gene function, as well as mechanisms of mtDNA replication, segregation, and maintenance. In addition, it would provide a powerful tool to model the ~113 mtDNA-localized confirmed pathogenic disease mutations^2^, which are now studied primarily in patient-derived cells with complex genetic backgrounds^3^.

For several decades, it has been possible to modify yeast^4–6^ and *Chlamydomonas reinhardtii*^7^ mtDNA through a laborious and inefficient process known as biolistic transformation, or “gene gun,” which directly deposits DNA into mitochondria. This approach has led to our current understanding of mitochondrial transcription and translation in eukaryotic mitochondria^8^. However, biolistic transformation cannot be used on mammalian mitochondria because of the high physical damage caused by microprojectiles^9^ and relatively inefficient homologous recombination, which limits the integration of foreign DNA^10,11^. Thus, recent studies have explored different approaches, including sequence-specific restriction enzymes that cut mtDNA within porcine oocytes^12^, mice^13^, plants^14^, metazoans^15^, and human cells^16–18^. The most sophisticated new technologies use programmable DNA binding proteins, TALEs (Transcription Activator-Like Effectors) and Zinc Fingers (ZF), to target non-specific endonucleases (e.g., Fok1^19,20^) or single base editors (e.g., DddA_tox_ ^21^) to specific mammalian mtDNA loci.

Although powerful, TALEs and related approaches lack the versatility and robustness of CRISPR-based methods. For example, only the DddA base editor ^21^ can be used in TALE-based targeting because it possesses unwindase activity; unlike CRISPR proteins, TALEs do not open double-stranded DNA^22^. Yet DddA (and a DddA-TadA fusion^23^) can only catalyze C->T or A->G edits, but not other transitions or transversions. TALE proteins also tend to be large (~160 kD) due to their need for long repetitive sequences, which presents a challenge for gene delivery.

It would be beneficial to bring CRISPR’s ease of use and extensive toolbox permitting insertion or deletion of nucleotides^24,25^, imaging^26^, and gene expression regulation^27,28^, to studies of the mitochondrial genome. Delivery of the CRISPR protein component is in principle straightforward and can be achieved by fusing the protein to a mitochondrial targeting sequence (MTS). However, there is no equivalent strategy for targeting synthetic RNAs to mitochondria.

Most nuclear-encoded RNAs are excluded from the mitochondria. Yet many species import one or more specific endogenous RNAs from the cytosol into the mitochondrial matrix because their mtDNA does not encode a complete set of tRNAs and/or rRNAs to carry out intra-mitochondrial protein translation^29^ – offering clues to how this barrier might be overcome. An extreme example is provided by trypanosomes, whose mtDNA does not encode any tRNAs, requiring the organism to import at least 20 different tRNAs from the cytosol to enable protein translation by mitochondrial ribosomes^30,31^. However, the mechanisms of RNA import are unclear in most species^29^. There is also no clear consensus that mammalian mitochondria import any nuclear-encoded RNAs (see Supplementary Discussion).

For this reason, we started our technology development with the budding yeast *Saccharomyces cerevisiae*, for which the import of two tRNAs (tRNA^Gln^ and tRNA^Lys^) from the cytosol into mitochondria is well-established^29,32^. Furthermore, the mechanism for tRNA^Lys^ import has been studied, and is thought to involve complexation to the mitochondrial tRNA synthetase Msk1, mediated by the enolase paralog, Eno2 (**Figure S1**)^33–35^. tRNA^Lys^ may cross the mitochondrial membrane in complex with Msk1, via the TOM/TIM protein import pathway. Additional benefits of yeast include homoplastic mitochondria (mtDNA sequences are identical within each cell) and the ability to construct mtDNA-based reporters can be constructed using biolistic transformation^6^.

We begin our study by creating a yeast reporter strain for mtDNA base editing that survives on arginine-deficient growth medium only if a stop codon is corrected. We then use this strain to screen a library of 30 potential RNA import motifs and identify a 40 nt RNA aptamer that enables the mitochondrial entry of a nuclear-encoded sgRNA. In combination with a mitochondrial-targeted base editor (ABE), we confirm the editing of the arginine biosynthesis gene *ARG8*^*m*^ within mtDNA. We also characterize the entry of sgRNA into mitochondria by biochemical fractionation and find that entry depends on both ATP and Msk1. Our efforts pave the way for CRISPR-based editing of the mitochondrial genome in eukaryotes.

## Results

### A sensitive reporter of CRISPR activity in mitochondria

Given the low efficiency of biolistic transformation (~1 in 10^11^) and the anticipated inefficiency of RNA import into mitochondria, we developed a sensitive reporter of CRISPR activity capable of detecting very low-frequency events. We wished to couple mtDNA editing to yeast survival on a selective medium, enabling enrichment of rare cells harboring the desired mtDNA edit. To do so, we repurposed a mitochondrial translation reporter^47, 36^ in which the arginine biosynthesis gene *ARG8* is relocalized from nDNA into mtDNA. In yeast lacking nuclear *ARG8*, biolistic transformation^4,37^ introduced *ARG8*^*m*^ into mtDNA, with a point mutation that converts a glutamine codon into a premature stop codon (CAA->TAA, Q79*). The resulting ARG8^m*^ strain (**Figure 1A**) requires exogenous arginine to grow and thus does not proliferate in (-)Arg medium (**Figure S2A**).

**Figure 1.**
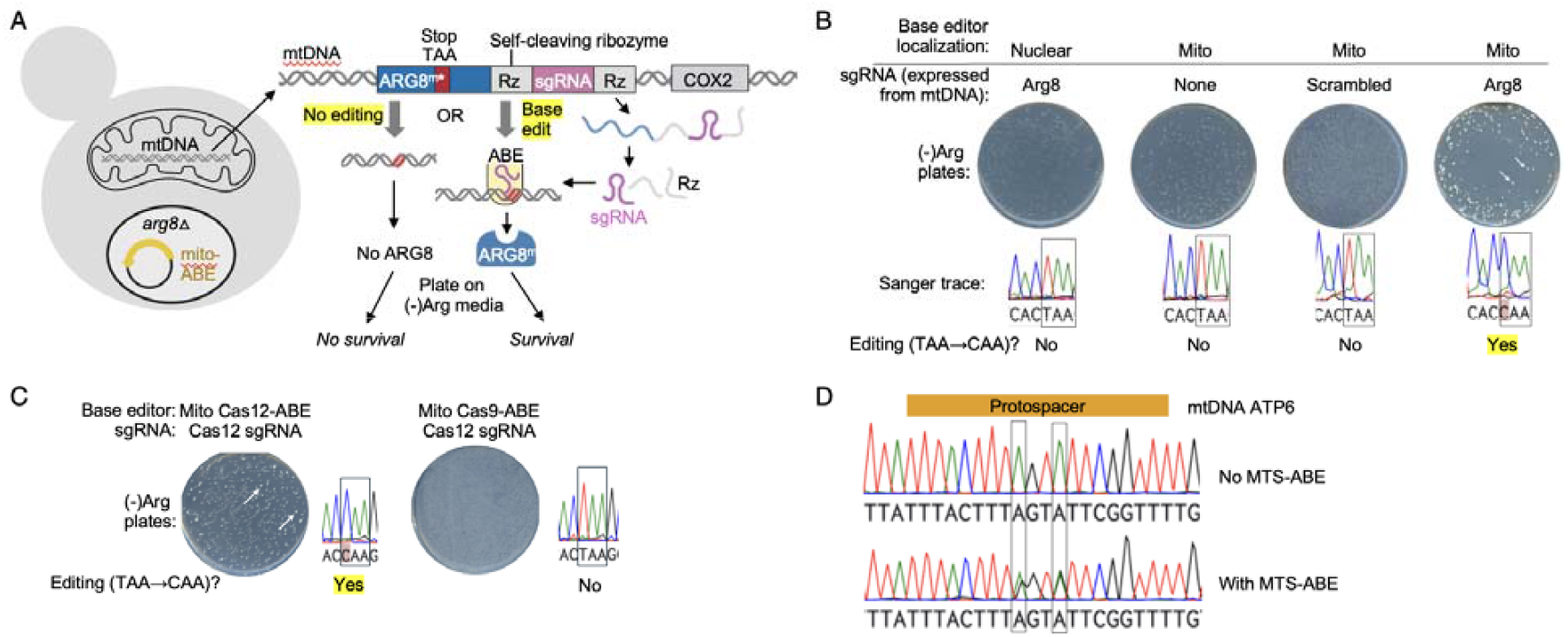
Reporter of mitochondrial CRISPR editing activity in *Saccharomyces cerevisiae*. (**A**) Reporter design. The arginine biosynthesis gene ARG8 was deleted from the nuclear genome (*arg8*_△_) and inserted into the mitochondrial genome (mtDNA) with a stop codon (*ARG8*^*m\**^). The sgRNA expression cassette is flanked by two self-cleaving ribozymes (Rz). Upon base editing by mitochondrial-targeted ABE protein (mito-ABE), full-length functional ARG8^m^ protein is produced, enabling cells to grow on arginine-deficient ((-)Arg) media. (**B**) Validation of reporter assay. Growth on (-)Arg plates is observed only when the base editor (Cas9-ABE) is targeted to mitochondria rather than the nucleus, and *ARG8*^*m\**^*-* targeting sgRNA is expressed from the mitochondrial genome. Bottom: sequencing results from individual colonies. For negative controls, colonies were picked directly from transformation plates before replica plating onto (-)Arg. n=8 colonies per condition from three independent biological replicates. (**C**) Cas12-ABE also edits ARG8^m*^ in mtDNA. Right: negative control with Cas9-ABE which is mismatched to the Cas12 sgRNA used here. sgRNA is expressed from mtDNA. This experiment was repeated 3 times with similar results. (**D**) Editing of an endogenous locus on mtDNA. sgRNA with a protospacer targeting part of the ATP6 gene on mtDNA was expressed from mtDNA as in (C). Top: No editing in the absence of MTS-Cas12 ABE. Bottom: A->G editing at two positions within the protospacer when MTS-Cas12 ABE i expressed.

To test the background reversion rate of this reporter strain - which places a lower bound to our assay sensitivity - we plated 40 billion ARG8^m*^ yeast cells on (-)Arg medium and cultured for 14 days. 97% of yeast cells retained their mtDNA and were able to grow in medium requiring active OXPHOS (**Figure S3B**). No colonies were observed over this time period, suggesting that the frequency of spontaneous mutation (T back to C at codon 79) is lower than 1 in 4×10^10^. Thus it should be possible, at least in principle, to detect rare editing events, as little as 1 in 10 billion. Interestingly, Q79 is highly conserved among ARG8 genes (**Figure S3A**). **Figure S3C** shows that other single nucleotide mutations at codon 79, which produce amino acids other than Q, mostly result in loss of function, with colonies unable to survive on (-)Arg medium.

The difference in growth between ARG8^m*^ and corrected ARG8^m^ provides a means to detect low-frequency editing events that repair the stop codon. ABE is a 187 kD adenine base editor^38^ that uses an RNA-guided Cas protein fused to an adenine deaminase to effect an A->I mutation. Mismatch repair converts the I/T pair to an I/C pair and then a G/C pair^38^, producing glutamine instead of stop during protein translation, thus restoring the cells’ ability to grow on (-)Arg medium.

### CRISPR editing with sgRNA expressed from the mitochondrial genome

Using our reporter, we first tested the feasibility of CRISPR editing in mitochondria when the need for RNA import is bypassed. To do this, we created an expression cassette with sgRNA flanked by two self-cleaving ribozymes^39,40^ and fused this to the 3’ end of our *ARG8*^*m\**^ reporter within mtDNA (**Figure 1A**). This modification did not affect the strength of the selection, as transformed cells were still unable to proliferate in (-)Arg medium (**Figure S2A**). We transformed yeast with ABE fused to an SU9-derived mitochondrial targeting sequence and GFP^41^. GFP fluorescence was observed in mitochondria in these cells (**Figure S2B**). Upon Sanger sequencing, we found that ~70% of cells expressing both MTS-ABE-GFP and mtDNA-encoded sgRNA harbored the desired edit (**Figure S2C**). Among Arg+ colonies, the frequency was 100% (**Figure 1B**). Negative controls in which sgRNA or ABE was omitted, ABE was localized to the nucleus instead of mitochondria, or sgRNA was replaced by a non-targeting control did not increase growth on (-)Arg medium above background (**Figure 1B**).

To assess generality, we tested CRISPR editing at a different position within the *ARG8*^*m*^ gene (TGA -> TAA, W168*). A similar degree of editing was observed (**Figure S2I**). We also tested a different Cas12-based enzyme (the base editor lbCpf1)^38^ with its corresponding sgRNA (**Figure S2J**). Cas12ABE targeted to the mitochondria also produced editing of *ARG8*^*m\**^ when its cognate sgRNA was expressed from mtDNA (**Figure 1C, Figure S2K**). Finally, we demonstrated editing of an endogenous locus, converting A to G within the ATP6 gene in mtDNA (**Figure 1D**). Our results demonstrate that the mitochondrial environment is compatible with efficient CRISPR editing.

### Screening for RNA-based mitochondrial import motifs

We then turned to the central challenge of this work: discovering an RNA motif that could, when fused to sgRNA, mediate its import from the cytosol into the mitochondrial matrix (**Figure 2A**). To do so, we first prepared a library of potential import motifs (IM) (**Figure 2B**) based on: (i) yeast tRNA^Lys^ and tRNA^Gln^ sequences that are naturally imported into yeast mitochondria via Msk1 and Eno2^34^; (ii) fragments of tRNA^Lys^ that were found to have high affinity to Msk1^42,43^; (iii) tRNA^Lys^-derived fusions that have been found to improve RNA import into mitochondria^42^; (iv) human 5S rRNA, H1 RNA, MRP, tRNA^Lys^, and hTERC that have been reported to enter human mitochondria^44,45^ in isolated studies; and (v) RNA sequences that have been reported to enter mitochondria of *Leishmania*^46^ and chloroplasts^47^.

**Figure 2.**
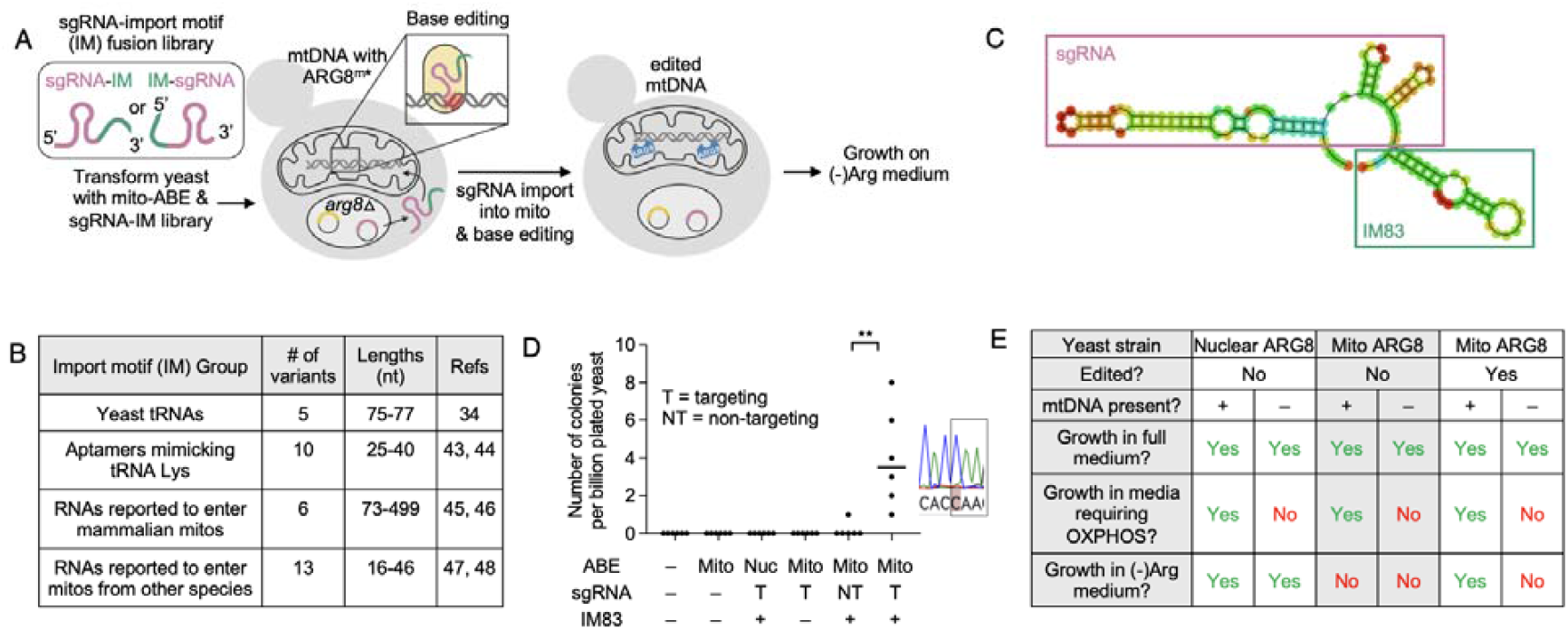
Screening identifies an import motif (IM) that enables sgRNA import into mitochondria and editing of mtDNA. (**A**) Design of the screen. Both mito-ABE and the sgRNA-IM (import motif) librar are expressed from nuclear plasmids. sgRNA-IM import into mitochondria enables CRISPR editing of ARG8^m*^ gene in mtDNA. Edited cells can then grow on arginine-deficient media. Left: Import motifs were fused to both 3’ and 5’ ends of sgRNA. (**B**) Composition of import motif library. (**C**) Predicted structure of the sgRNA-IM83 fusion RNA (from RNAfold^52^). (**D**) Controls for mtDNA editing by the winning clone, sgRNA-IM83. Surviving colonies on (-)Arg plates were counted after performing the assay in (A). T, targeting guide. NT, non-targeting guide. **p<0.05. All growth on (-)Arg media contained the desired mutation (TAA->CAA) as well as the targeting guide. (**E**) Growth on arginine-deficient media is conferred by edit to the mitochondrial genome. For each strain (nuclear ARG8, mito ARG8 before/after editing), a petite strain lacking mtDNA was generated, and growth on different media was assessed. Each condition was repeated 2 times.

The lengths of these RNA motifs range from 16-330 nucleotides (**Figure 2B**). Making no assumptions about the effect of fusion on import motif or sgRNA function, we attached this library of 30 sequences to either the 5’ or 3’ end of the *ARG8*^*m\**^-targeting sgRNA, producing a collection of 60 fusions (**Figure 2A**). We transformed yeast cells with the IM-sgRNA fusion library and a plasmid encoding MTS-ABE. Selection was performed by replica plating transformants onto (-)Arg selection plates. After seven days, we observed a single Arg+ colony expressing the library member sgRNA-IM83 (**Figure 2C**). Sanger sequencing of PCR-amplified *ARG8*^*m*^ gene confirmed the desired edit. We then scaled up the experiment to estimate editing efficiency. Across six replicates in which we plated 1 billion cells expressing sgRNA-IM83 and MTS-ABE, we observed an average Arg+ colony frequency of 4 ± 2.6×10^-9^ (**Figure 2D**). All Arg+ colonies harbored the desired edit according to Sanger sequencing. For reference, this efficiency exceeds that of biolistic transformation by ~400-fold^4,37^.

Controls with IM83 omitted (sgRNA only), non-targeting protospacer replacing the targeting protospacer in sgRNA, or ABE lacking a mitochondrial localization sequence yielded no Arg+ colonies (**Figure 2D**). These results suggest that the editing is driven by the combination of ABE and sgRNA instead of background errors in mtDNA replication or repair.

Additionally, we performed a test to determine if growth on (-)Arg requires mtDNA, and is not somehow conferred by changes in the nuclear genome. To do this, we generated “petite” yeast cells lacking mtDNA by passaging in medium containing ethidium bromide. Petite cells encoding nuclear *ARG8* retained the capacity to grow in (-)Arg medium, whereas petite strains with edited *ARG8*^*m*^ lost this ability (**Figure 2E**). These results show that edited *ARG8*^*m*^ requires mtDNA to grow in arginine-deficient medium.

### Biochemical detection of sgRNA-IM83 import into mitochondria

Manifestation of the growth phenotype in (-)Arg medium requires multiple biochemical steps including sgRNA import, ABE import, base editing, replication repair, and translation of the Arg8 protein. To directly read out and quantify sgRNA import, we used biochemical fractionation to purify mitochondria from yeast cells via rigorous, multi-step centrifugation^48^ (**Figure 3A**). To remove any contaminating RNAs that might adhere to the outside of purified mitochondria, we treated the samples with RNase prior to analysis of mitochondrial RNA content. Mitochondrial purity was confirmed by cytosolic mRNA depletion and mitochondrial RNA enrichment (**Figure S3D**).

**Figure 3.**
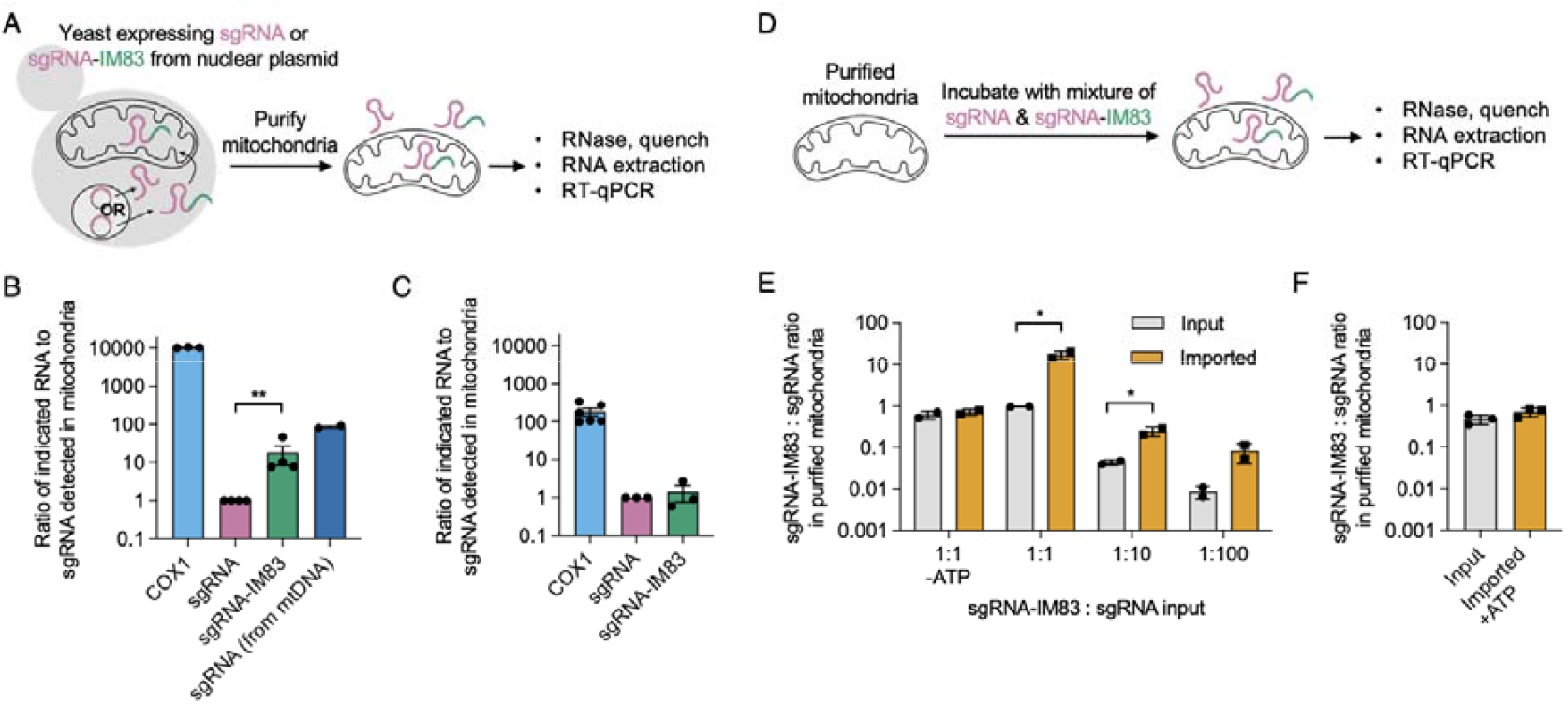
Direct biochemical detection of sgRNA import into yeast mitochondria. (**A**) Assay for detecting sgRNA mitochondrial import. Mitochondria are purified and then analyzed for RNA content by RT-qPCR. (**B**) Results from assay in (A), with sgRNA and sgRNA-IM83 expressed from a 2u plasmid. No MTS-ABE was expressed in these cells. Abundance of each transcript is normalized to that of ACT1, a cytosolic mRNA, then compared to sgRNA. COX1 is a positive control mitochondrial mRNA expressed from mtDNA. **p<0.01. Each condition was repeated 2-4 times. (**C**) Same as (B) but using yeast cells expressing MTS-ABE. (**D**) Assay for in vitro sgRNA import into purified mitochondria. (**E**) Results from assay in (C) with different sgRNA-IM83:sgRNA input ratios. Data from two biological replicates. *p< 0.05. (**F**) Same as (E) but using mitochondria purified from yeast cells expressing MTS-ABE.

In purified mitochondria from yeast cells expressing nuclear-encoded sgRNA-IM83 or untagged sgRNA, we detected 10 times more sgRNA-IM83 within the mitochondrial matrix than sgRNA lacking the import motif (**Figure 3B**). Swapping out the protospacer on sgRNA to a sequence not targeting *ARG8*^*m\**^ preserved the enrichment of IM83-fused guide (**Figure S3B**). The enrichment of sgRNA-IM83 over sgRNA was also observed in mitoplasts, in which the outer mitochondrial membrane (OMM) is selectively removed by digitonin; this indicates that sgRNA-IM83 reached the mitochondrial matrix (**Figure S3A, C**).

As a second test of sgRNA import, we incubated purified mitochondria (from wild-type yeast) with *in vitro* transcribed sgRNA-IM83 or control sgRNA (**Figure 3D**). After 20 minutes, we examined RNA protected from external RNAse digestion (i.e., RNA that had reached the interior of mitochondria). We consistently measured 7-12 fold more internalized sgRNA-IM83 than sgRNA in these experiments (**Figure 3E**). Import depended on ATP (**Figure 3E**), consistent with known mechanisms of natural tRNA import into yeast mitochondria (**Figure S1**)^49^.

### Msk1 enhances the import of sgRNA-IM83

The IM83 RNA import motif is a chimera of two arms of tRK1^Lys^(CUU)^42^, a yeast tRNA imported through complexation with Msk1, a mitochondrial lysyl tRNA synthetase^43^ (**Figure S1**). Knocking out *MSK1* abolished mitochondrial import of sgRNA-IM83 in living yeast (**Figure 4E**). A potential caveat to this *in vivo* experiment is that Msk1 function is also required for mitochondrial translation and thus its knockout has multiple physiological effects. Thus, as an additional test of whether Msk1 promotes import of sgRNA-IM83, we repeated the *in vitro* import assay from **Figure 3D** with purified mitochondria but added recombinant Msk1 protein (**Figure 4A**). Addition of Msk1 with a mitochondrial targeting sequence (preMsk1) increased the ratio of RNAse-resistant sgRNA-IM83:sgRNA by 2.3-fold, whereas truncated Msk1 lacking the mitochondrial targeting sequence (tMsk1) did not (**Figures 4B-C** and **S5**). We also observed direct binding between sgRNA-IM83 and preMsk1 protein using an electrophoretic mobility shift assay (**Figure 4D**). The interaction was dependent on IM83, because binding was not observed when IM83 was deleted from sgRNA (**Figure 4D**). Thus, mitochondrial Msk1 is sufficient to enhance the import of IM83-sgRNA.

**Figure 4.**
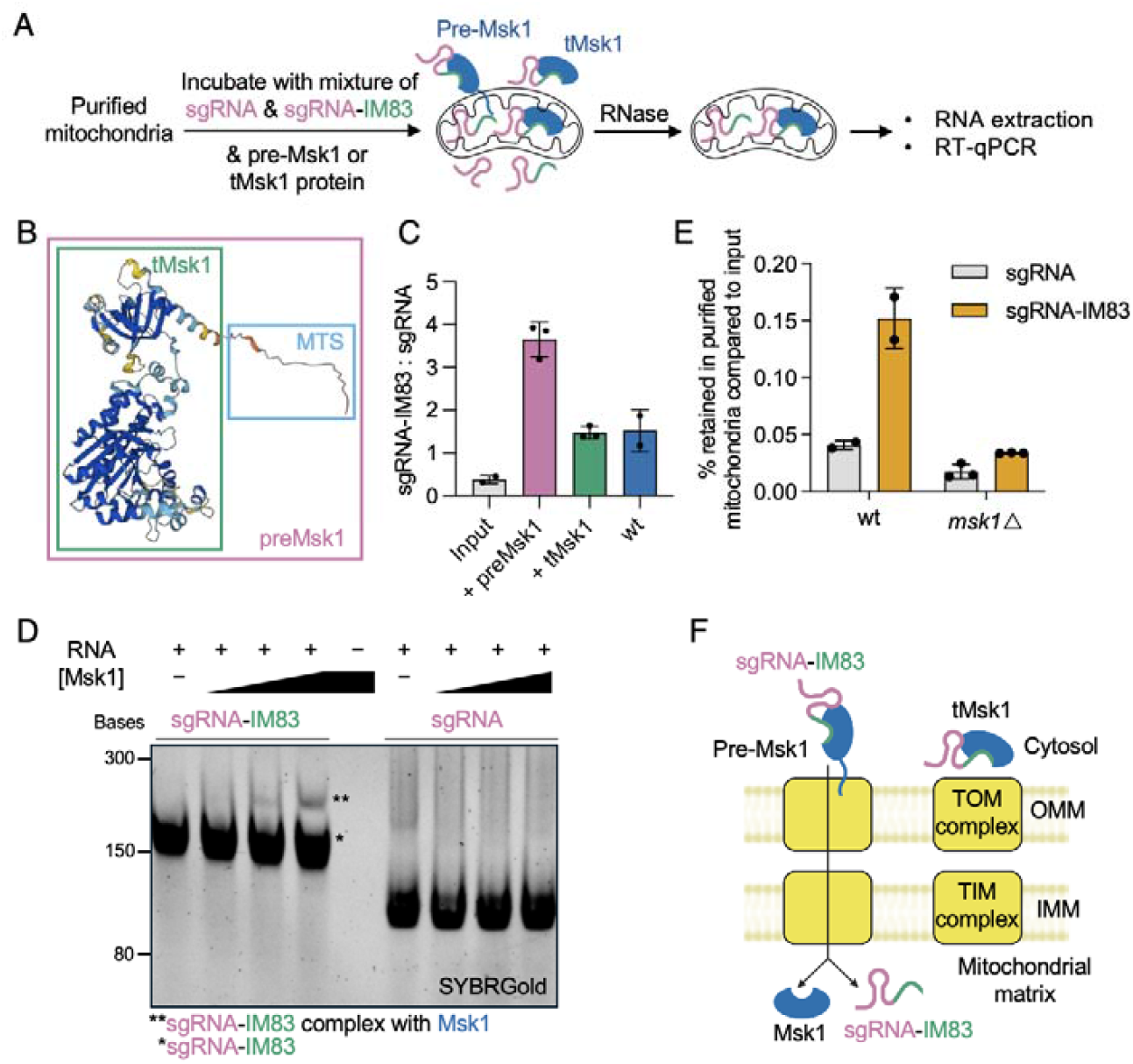
Msk1 enhances sgRNA-IM83 entry into mitochondria. (**A**) Assay schematic. tMsk1 i truncated preMsk1 without a mitochondrial targeting sequence. (**B**) Alphafold model of Msk1 from *Saccharomyces cerevisiae*. Truncated Msk1 (tMsk1) is preMsk1 without the mitochondrial targeting sequence (MTS). (**C**) Result from experiment in (A) showing that addition of pre-Msk1 enhances *in vitro* import of sgRNA-IM83. wt, no preMsk1 or tMsk1 added. Data from 2-3 biological replicates per condition. (**D**) Gel shift assay showing that Msk1 protein binds to sgRNA-IM83 (**), but not to sgRNA. This experiment was repeated three times. Uncropped blot in Figure S4B. (**E**) Knock out of *MSK1* reduce sgRNA-IM83 import. Experiment was performed as in Fig. 3A. Data is reported as the percentage of guide retained in purified mitochondria following RNAse digest, compared to the input (whole cell lysate). Data from 2-3 biological replicates. (**F**) Proposed mechanism of sgRNA-IM83 import via binding to preMsk1 protein. On right, tMsk1 lacking the mitochondrial targeting sequence is unable to import via the TOM/TIM import complex. OMM, outer mitochondrial membrane. IMM, inner mitochondrial membrane.

Interestingly, knocking out *ENO2*, which folds the imported tRNA^Lys^ into an alternative structure before passing it to Msk1^33^, did not affect the editing efficiency of sgRNA-IM83 (**Figure S5E**). This suggests that unlike the full-length tRNA^Lys^, IM83 may not require refolding for mitochondrial import. Taken together, our results support a protein-assisted model of sgRNA-IM83 import (**Figure 4F**), enabling CRISPR-based editing of the mitochondrial genome.

### Attempts to increase mtDNA editing efficiency

Although more efficient than biolistic transformation, IM83-mediated CRISPR mtDNA editing, with an efficiency of just ~4 in 10^9^, has limited practical utility compared to the power of such methods in the nucleus. We considered several factors that could be limiting the efficiency of editing in mitochondria. First, the biochemical characterization of sgRNA uptake suggests that imported sgRNA (from the cytosol, encoded in nDNA) is present at ~10-fold lower levels in mitochondria than sgRNA expressed directly from mtDNA (**Figures 3B, S4B**). While this is a large difference, it is far exceeded by the difference in editing efficiency when using imported sgRNA versus mtDNA-expressed sgRNA (1 in 2.5 × 10^8^ versus 1 in 40, a difference of ~6 million-fold). Thus, additional factors beyond just sgRNA abundance must account for the loss of editing efficiency.

We wondered if the IM83 motif, when fused to sgRNA, could sterically interfere with CRISPR editing. However, both sgRNA-IM83 and untagged sgRNA gave similar ARG8^m*^ editing efficiencies when expressed from mtDNA (**Figure S6C**). We tested altering the location of IM83 by inserting it into the major stem-loop of sgRNA (sgRNA-msIM83). sgRNA-msIM83 too showed similar editing efficiency (when expressed from a nuclear plasmid) (**Figure S6A**) and a similar degree of mitochondrial import as sgRNA-IM83 (**Figure S6B**).

We hypothesized that reducing the size of the imported sgRNA might facilitate mitochondrial uptake and improve editing. To test this, we split the 147 nt sgRNA-IM83 into a 82 nt tracrRNA component (expressed from mtDNA) and a 82 nt crRNA-IM83 component (imported by IM83) (**Figure S7A**). However, no survival on (-)Arg medium was observed. In a control, we noticed that when testing this split system in the nucleus, the editing efficiency was only 0.5% that of the unsplit sgRNA. Optimization to increase annealing gave only modest improvements (**Figure S7B**).

### MTS-ABE expression causes mitochondrial import toxicity, reducing sgRNA uptake

In the course of our experiments, we observed that MTS-ABE expression slowed growth of yeast. Upon further investigation, we noticed a decrease in MitoTracker signal when expressing MTS-ABE from a nuclear plasmid, indicative of a reduction in the mitochondrial membrane potential (**Figure S8A**). This effect was observed across all haploid and diploid strains that we tested (**Figure S8B**). By contrast, a control expressing ABE in the nucleus (NLS-ABE) did not alter mitochondrial membrane potential (**Figure S8B**). Because sgRNA import into mitochondria depends on a proton gradient across the mitochondrial membrane (abolished by valinomycin treatment, **Figure S8C**), the negative effect of MTS-ABE on mitochondrial membrane potential could be a major factor limiting the efficiency of mtDNA CRISPR editing.

To directly test the effect of MTS-ABE expression on sgRNA import, we analyzed mitochondria purified from yeast cells expressing MTS-ABE and sgRNA-IM83. The ~10-fold enrichment of sgRNA-IM83 we previously observed using yeast lacking MTS-ABE (**Figure 3B**) was absent from cells expressing MTS-ABE (**Figures 3C**). Furthermore, we observed no *in vitro* import of sgRNA-IM83 in mitochondria isolated from MTS-ABE-expressing cells (**Figure 3F**).

The “import toxicity” caused by MTS-ABE could be the result of ABE’s large size (187 kD), possible misfolding/aggregation, or off-target editing activity. To counteract this toxicity, we attempted several strategies. First, we reduced MTS-ABE expression level by using a weaker promoter (**Figure S8D**). Although membrane potential was restored, the low level of MTS-ABE meant that the editing efficiency, even with mtDNA-expressed guide, was very low, though still detectable (**Figure S8F**). Second, we tested a promoter of intermediate strength, ADH1, which indeed lessened the impact of MTS-ABE on mitochondrial membrane potential (**Figure S10A, B**). Encouragingly, in these cells, imported guide was still detected by mitochondrial fractionation (**Figure S10D**), and editing efficiency from mtDNA-expressed sgRNA was unchanged (**Figure S10C**). However, editing with nuclear-expressed sgRNA was reduced compared to editing by MTS-ABE expressed from the stronger GPD promoter (**Figure S10E**). We hypothesize that ABE level in the mitochondria may become a limiting factor to editing when the amount of sgRNA is low – as is the case when it must be imported but not when it is expressed from mtDNA.

Next, we reduced the size of ABE by truncating the REC2 or HNH domain (size from 187 kD to 170 kD/169 kD). ΔHNH rescued membrane potential in some cells (**Figure S8E**), but no editing was observed at all (**Figure S8F**). ΔREC2 did not rescue the membrane potential. We then tested a new mutation^50^, V28F, reported to decrease off-target RNA editing activity. However, no editing was observed at all on mtDNA (**Figure S8G**). Finally, to bypass the need for ABE import altogether, we directly expressed ABE from mtDNA, either as a fusion to ARG8^m^ or in a separate cassette (**Figure S9A, B**, see Supplementary Text). Although we observed edits in a positive control expressing ABE^m^-ARG8^m*^-sgRNA, the efficiency was only ~10% of what we observed when expressing MTS-ABE from a plasmid (**Figure S9C**), and no growth was detected on (-)Arg medium when we attempted editing with ABE^m^-ARG8^m*^. Diagnostic experiments suggest that low protein expression is the most likely cause of failure (see Supplementary Text).

### Harassing genetic diversity

Finally, we considered the contribution of genetic backgrounds. We observed that different yeast strains exhibited different CRISPR editing efficiencies. Our ARG8^m*^ reporter strain was generated by sporulation of a diploid (**Figure S2D**). The resulting haploids have identical mtDNA but different combinations of parental nDNA backgrounds. When we performed the editing assay and detected growth on (-)Arg media, we observed editing efficiencies ranging from 2.5% to 97.5% across ten different strains (**Figure S2C**), consistent with multiple genetic contributors to editing efficiency. Quantification of sgRNA expression level from mtDNA in these strains did not show any correlation with editing efficiencies (**Figures S2E-F)**, nor did the percentage of cells expressing ABE (**Figures S2G-H**). Variations in other factors, such as mtDNA repair pathways, may explain the differences in editing efficiencies. Further investigation of the underlying mechanisms likely holds promise for additional improvements in editing efficiency.

## Discussion

In this study, we developed a sensitive reporter of mtDNA editing in yeast, demonstrated that CRISPR is active in yeast mitochondria, and identified an RNA import motif (IM) that enables the entry of sgRNA into mitochondria. We confirmed that sgRNA-IM could mediate a low but detectable amount of mtDNA base editing. Furthermore, we found that mtDNA editing efficiency is affected by the nuclear genetic background of yeast. Our work provides a foundation for mtDNA CRISPR editing, identifies central issues for future optimization, and generates assays for studying RNA import and functions of the mitochondrial genome.

We examined the likely mechanism of IM83-mediated sgRNA import. We showed that import requires ATP and is enhanced by preMsk1, suggesting a resemblance to the natural tRNA import mechanism. In the natural pathway (**Figure S1**), cytosolic tRNA^Lys(CUU)^ first binds to Eno2, which is thought to act as a chaperone that folds the tRNA into an alternative F-stem structure^33^. The tRNA-Eno2 complex transits to the surface of the mitochondrion, where the tRNA is handed to preMsk1, which is translated near the OMM^33^. The preMsk1-tRNA complex is then imported via the canonical TOM/TIM protein import pathway^35^. The IM83 motif was originally designed^60^ by fusing the D arm and amino acid stem of a modified tRNA^Lys^ that has high import into yeast mitochondria^43^. We also observed that editing is not affected in the absence of Eno2, suggesting that IM83 might not need to be refolded for import as tRNA^Lys^ does. Our working model is that sgRNA-IM83 enters mitochondria through the protein import pathway via binding to preMsk1 but does not require Eno2 for this process.

Despite our best efforts, we could only achieve CRISPR editing of the yeast mitochondrial genome with low efficiency when utilizing nuclear-encoded sgRNA. Mitochondrial fractionation and qPCR showed that imported sgRNA levels are at least 10-fold lower than sgRNA expressed from mtDNA (which gave efficient mtDNA editing). We also explored entirely different strategies for delivering sgRNA into mitochondria, such as fusing the sgRNA to PP7 and co-expressing an MTS-tagged PP7 coat protein (**Figure S11**); however, none yielded any detectable editing.

We discovered that a major factor limiting mtDNA editing efficiency is the effect of mitochondrially-targeted base editor on mitochondrial membrane potential. By diminishing membrane potential, MTS-ABE expression further reduces sgRNA import (**Figures 3C, 3F**). Interestingly, a previous study also observed that mitochondrial import of a similarly sized protein, ATP2-lacZ (155 kD compared to 187 kD for MTS-ABE), caused mitochondrial dysfunction ^51^. We tried many strategies to overcome this toxicity including reducing ABE size and expression level and expressing ABE from mtDNA to bypass the need for mitochondrial import. However, these experiments revealed challenges inherent to balancing MTS-ABE’s effect on mitochondrial membrane potential with required expression levels and activity needed to achieve editing. Future innovation to mitigate ABE import toxicity could produce dramatic improvements to mtDNA CRISPR editing efficiency. Dissecting natural genetic variation that impacts editing efficiency may reveal additional avenues for optimization.

The long-term goal is efficient CRISPR editing of mtDNA in all eukaryotes, including humans. The density of disease-causing mutations in human mtDNA exceeds that of nDNA by >100 fold^2^. Whether aptamer-fused sgRNAs can overcome the barrier to entry via the same pathways in humans remains to be established, although the core factors we have identified are broadly conserved. Our work establishes that it is possible to deploy CRISPR in the mitochondria and point toward multiple strategies to enable mtDNA manipulation with a level of throughput and control that can bring insight into fundamental biological questions about mitochondrial inheritance, metabolism, and energy production, and potentially unlock translational approaches to the treatment of mitochondrial diseases.

## Acknowledgments

We are extremely grateful to Thomas Fox for sharing key strains and plasmids, and providing valuable advice on mitochondrial genetics. We also thank Juan Alfonzo for advice on yeast mitochondria purification and tRNA import; David Liu for advice on ABE and sharing of plasmids; Virgina Walbot for sharing the biolistic transformation device; and members of the D.F.J and A.Y.T. laboratory for technical support and advice. This work was supported by the Chan Zuckerberg Biohub – San Francisco, and Stanford University.

## Author contributions

S.Y., D.F.J. and A.Y.T. designed research; S.Y. performed research; S.Y., D.F.J. and A.Y.T. analyzed data; S.Y., D.F.J. and A.Y.T. wrote the paper.

## Materials and Methods

### Cloning

All plasmids listed in the table were constructed with standard molecular biology methods. Double restriction digestion of the vector and Gibson assembly with amplified inserts were used to construct most plasmids, except sgRNA-expression plasmids which were cloned by ligation cloning and Golden Gate assembly. Plasmids were produced from chemically competent XL1-Blue bacteria and miniprep except for those used in biolistic transformation, which were produced from Midiprep (Zymopure II Plasmid Midiprep kit) for higher purity and yield.

### Yeast cell culture

Various strains of *S. cerevisiae* were cultured using standard yeast culture protocols. All transformations were performed with chemically competent yeast (Frozen-EZ Yeast Transformation II Kit, Zymo Research) following the manufacturer’s recommendations. Unless otherwise stated, 1.5 μg of total plasmids were transformed into 40 μl competent cells. Unless otherwise stated, transformed yeast were inoculated and grown in synthetic medium with the appropriate amino acid dropouts (SD-X, 6.7 g/L yeast nitrogen base without amino acids, specified g/L CSM-X (Sunrise Science Products), 20 g/L dextrose).

### Generation of mitochondrial ARG8^m*^ and other mtDNA mutants

A stop codon was introduced into yeast mitochondrial *ARG8*^*m*36^ to generate *ARG8*^*m\**^ as the reporter of mitochondrial base editing activity. This *ARG8*^*m\**^ was cloned into the vector pPT24^36^ between two EcoRI restriction sites using ligation cloning. The vector used for biolistic transformation, pPT24*-ARG8^m*^, contains *ARG8*^*m\**^ (together with a sgRNA expression cassette in some strains) flanked by the 5’ and 3’UTR of COX2 and the complete COX2 gene further downstream (see sequence map in supplementary information). pPT24*-ARG8^m*^ was integrated into the mitochondrial genome via biolistic transformation following this detailed protocol^4^. Briefly, DFS160 (a gift from Dr Thomas Fox) was cultured in YPR (20 g/L peptone, 10 g/L yeast extract, 20 g/L raffinose) for 3 days to saturation. 50 ml culture was concentrated to a concentration of 5 x 10^9^ yeast / ml. 0.1 ml of this concentrated culture was spread onto SD-Leu-sorbitol plates (0.77 g/L CSM-Leu, 6.7 g/L yeast nitrogen base, 50 g/L glucose, 1 M sorbitol, 100 mg/L adenine, 30 g/L bacto agar). For 4 shots, 10 ug of Leu carrier plasmid and the 60 μg of the appropriate pPT24 plasmid were mixed and precipitated onto 200 μl 70% ethanol washed tungsten beads (60 mg/ ml in 50% glycerol), then with 8 μl of 1 M spermidine and 20 μl of 2.5 M ice-cold CaCl_2_. This mixture was incubated on ice with occasional shaking for 10 minutes. The mixture was washed with cold ethanol at least twice until tungsten beads could be easily resuspended. Before loading onto the biolistic transformation device, 60 μl was used to resuspend the beads and 10 μl was used per shot. Operation of the biolistic transformation device (Biorad, PDS-1000 He™ system) was according to the manufacturer’s protocol except that the stopping screen is removed. 1100 psi disks and the highest position of the receiving petri plate were used in all bombardments to generate strains in this paper but these conditions might require optimization based on the individual machine. Where applicable, mitochondria with *ARG8*^*m\**^ integrated into the mitochondrial genome were further introduced to BY4742 strain for downstream experiments using standard cytoduction techniques.

### ARG8^m*^ editing assay

Yeasts with ARG8^m*^ reporter in mitochondria were first transformed with the specified MTS-ABE (SU9MTS-ABE(V106W) was used unless otherwise specified) on a centromeric plasmid. Transformed yeasts were selected on SD-Leu plates with 6.7 g/L yeast nitrogen base without amino acids, 0.77 g/L CSM-Leu (Sunrise Science Products), 20 g/L dextrose and 20 g/L bacto agar. To estimate the editing efficiency, colonies were directly picked from SD-Leu plates. Sections of ARG8^m^ were amplified (Forward primer: 5’-CACAAGAGGTAAAAATGCTAAATTATATGATGATGTAAATGGTAAAG-3’, reverse primer: 5’-AGCATATACAGCTTCGATAGCTTTTTCGAAAGC-3’) from a yeast crude lysate (boiled at 95 °C in 20 mM NaOH) and Sanger sequenced.

To detect growth in the absence of arginine, the transformation plates were replica plated onto 15 cm (-)Arg plates (6.7 g/L yeast nitrogen base without amino acids, 0.74 g/L CSM-Arg (Sunrise Science Products), 20 g/L raffinose, and 20 g/L bacto agar) and incubated at 30 °C. To confirm DNA change on yeast growing on (-)Arg medium, colonies were picked from (-)Arg plate. Sections of ARG8^m^ were amplified (Forward primer: 5’-CACAAGAGGTAAAAATGCTAAATTATATGATGATGTAAATGGTAAAG-3’, reverse primer: 5’-AGCATATACAGCTTCGATAGCTTTTTCGAAAGC-3’) from a yeast crude lysate (boiled at 95 °C in 20 mM NaOH) and Sanger sequenced.

### Screening of IM (import motif)-tagged sgRNA library

A library of sgRNA tagged with import motifs (IM) was constructed using standard ligation cloning into a 2µ plasmid under SNR52 promoter (map in Supplementary information). Yeasts with ARG8^m*^ were transformed with the sgRNA-IM expression plasmid and a centromeric Mito-ABE plasmid (GPD-SU9MTS-ABE(V106W) unless otherwise specified). Transformed yeasts were selected on synthetic medium lacking leucine and uracil (SD-Leu-Ura, 6.7 g/L yeast nitrogen base without amino acids, 0.67 g/L CSM-Leu-Ura (Sunrise Science Products), 20 g/L dextrose and 20 g/L bacto agar). In the first part of the screening, the transformation plate was directly replica plated onto (-Arg) medium on 15 cm dishes (approximately 0.5 billion cells) and cultured at 30 °C for 7 days. To confirm hits after screening, colonies on SD-Leu-Ura plates were scraped to grow in SD-Leu-Ura liquid medium (same makeup as solid medium except no agar) and grown for 2 days at 30°C before counting and spreading 1 billion yeast on a large square petri dish with solid (-)Arg medium.

### Generation of petite yeast and phenotyping

Respective yeast strains were passaged in YPD (20 g/L peptone, 10 g/L yeast extract, 2% dextrose) with 5 mg/ml ethidium bromide twice after 1:1000 dilution and overnight growth at 30 °C. About 100 yeasts were spreaded to YPD plates and replica plated onto YPEG (20 g/L peptone, 10 g/L yeast extract, 2% v/v glycerol, and 2% v/v ethanol, 20 g/L bacto agar) and then YPD plates after 2 days. Petite yeasts are able to grow on the second YPD plate but not on YPEG. Respective yeast strains were grown overnight to saturation in complete medium (YPD) to saturation, washed, and resuspended in PBS. 10 µl of resuspended yeasts were streaked onto YPD, YPEG, and (-)Arg plate, respectively.

### Mitochondrial fractionation

Mitochondria purification was carried out following the published method^48^ with some modifications. Briefly, yeast were grown in media with raffinose (YPR, SR-Leu or SR-Leu-Ura depending on requirements for plasmid maintenance) overnight. Cells were harvested and incubated with 0.1M Tris-SO_4_, 10 mM DTT at 30 °C for 10 min. Spheroplasting was completed in 2 ml/ gram yeast wet weight buffer A (1.2 M sorbitol, 20 mM potassium phosphate pH 7.4) with 1 mM DTT and 2.5 mg/g zymolase 100T (Sunrise Science Products). Spheroplasts were washed twice with ice-cold buffer A and all subsequent steps were performed on ice. Spheroplasts were homogenized in 0.6 M sorbitol, 20 mM Tris-HCl pH 7.2 and freshly added 0.5 mM PMSF with Avestin Emulsiflex C5 (for large samples) or a tight-fitting glass Dounce homogenizer (for small samples). The crude lysate was spined at 1,000 g for 5 min, then supernatant at 2,000 g for 5 min. The combined supernatant was spined the same way and the final supernatant was spined at 12,000g for 12 min. The crude mitochondria were gently resuspended in 0.6 M sorbitol, 50 mM Tris-HCl pH 7.2. In the final step, crude mitochondria were laid on top of a gradient of 2 mL 40% Percoll in 0.6 M sorbitol, 50 mM Tris-HCl pH 7.2 overlaid by 8 mL of 20% Percoll in 0.6 M sorbitol, 50 mM Tris-HCl pH 7.2. The gradient was spun for 30 min at 30,000 rpm in an SW41 rotor at 4 °C and the bottom brownish layer was extracted, further washed in 0.6 M sorbitol, 50 mM Tris-HCl pH 7.2, and pelleted as the purified mitochondria.

### In vitro mitochondrial import assay

In vitro transcribed sgRNA and sgRNA-IM83 (MEGAScript T7 Transcription kit, Invitrogen), incubated at 37 °C overnight for higher yield of small RNAs. The IVT reaction was purified by column (RNA Clean and Concentrator 25, Zymo Biosciences). Purified RNA was folded in an RNA folding buffer (5mM Tris pH 7.5, 15mM NaCl, 0.01mM EDTA) at 85°C for 2 min and cooled to room temperature. 25 μg Freshly purified mitochondria as above were incubated with folded RNA in the specified concentration in an import buffer (0.6 M sorbitol, 20 mM Tris-HCl pH 8.0, 2 mM DTT, 20 mM MgCl2, 2 mM EDTA and 5 mM ATP). ATP was omitted in the -ATP conditions. After 20 min, 10 U/ml of micrococcal nuclease and 100 U/ml of RNase A were added to remove the unimported RNA, and incubated with shaking at 30 °C for 20 min. RNAse activity was stopped by putting the mixture on ice. Mitochondria were washed once in import buffer at 4°C without ATP and immediately subjected to lysis and RNA extraction.

### RNA extraction, reverse transcription, and qPCR

RNA were extracted from mitochondria (miRNeasy Tissue/Cells Advanced Mini/Micro Kit, Qiagen) after RNAse treatment and washing as above. Further DNAse digestion was performed (Turbo DNAase, Invitrogen). RNA was then reverse transcribed (Superscript VILO, Invitrogen), and transcripts were quantified with qPCR (SYBR Green 2x MasterMix, ThermoFisher).

### Recombinant preMsk1 and tMsk1 production

BL21 DE3 E. coli were transformed with a plasmid containing sequences encoding yeast preMsk1 or tMsk1 with 6xHis tag at the N-terminal (Plasmid named Msk1 and tMsk1 in the plasmid table). A pre-culture was grown overnight from a single colony in 5ml LB supplemented with carbenicillin at 37°C. On the second day, the pre-culture was diluted to 1 L of LB supplemented with antibiotics and grown at 37°C until OD600 reached 0.4 to 0.6. The culture was cooled to room temperature, and protein expression was induced with 0.2 mM IPTG overnight at 18°C with shaking at 220 rpm. After 18 hours, bacteria were pelleted (8000 g, 10 min), lysed in 25 ml lysis buffer (10 mM Na_2_HPO_4_, 0.5 M NaCl, 20 mM imidazole, pH 8.0) and 1 mg/ml lysozyme. DNA was sheared further by sonication. The lysed culture was pelleted at 16,400g for 30 min at 4°C. The supernatant was purified using a Ni-NTA column and eluted with a gradient of imidazole (20 mM, 100 mM, 200 mM, 300 mM, 400 mM, 2 column volume each) in lysis buffer with 0.2% CHAPSO. The pooled fractions were further purified through size exclusion chromatography (24 ml Superdex 200 increase column, GE Healthcare) equilibrated in protein storage buffer (20 mM Tris-HCl, pH 7.5, 150 mM NaCl, 10 mM MgCl2, 10% glycerol, 5 mM DTT) with a flow rate of 0.2 ml/ min. The peak fractions were pooled and concentrated by centrifugal ultrafiltration to ~0.8 mg/ml. Purified proteins were analyzed with SDS-PAGE gel electrophoresis (Figure S4A).

### Electrophoretic mobility shift assay (EMSA)

In vitro transcribed sgRNA and sgRNA-IM83 were folded as above. Varying concentrations (0.1, 0.2, 0.4 µM) of Msk1 protein, as indicated (Figure 4D), were incubated with folded sgRNA and sgRNA-IM83 (0.5 µM unless otherwise indicated) for 10 min at room temperature in 20 mM Tris-HCl, pH 7.5, 150 mM NaCl, 10 mM MgCl2, 10% glycerol. Following incubation, the reaction mixtures were analyzed on an 8% Tris-borate-EDTA gels (Sigma) with 0.5x TBE buffer. The gel was incubated with 1:10,000 SYBR Gold nucleic acid gel stain (Invitrogen) at room temperature for 10 min and visualized with a gel imager with appropriate settings for SYBR Gold (ChemiDoc, BioRad).

