## Supplementary Information for "Towards CRISPR-based editing of the mitochondrial genome in yeast"

**This PDF file includes:**

Supplementary Text

Figures S1 to S11

Tables S1 to S2

Supplementary Methods

SI References

**Supplementary Text**

**Expression of ABE from mtDNA to circumvent MTS-ABE toxicity.** We tested several different strategies for expressing ABE from mtDNA, summarized in **Figure S9**. First, we prepared a cassette consisting of the ABE gene fused at the 5’ end of our ARG8*-HH-sgRNA-HDV reporter (**Figure S9A**). This construct produced low but detectable editing of the target ARG8 (**Figure S9C**) but unfortunately, the edited ARG8 levels were so low that they could not rescue growth on (-)Arg medium. To understand why this fusion cassette was inadequate, we separately tested two control constructs: MTS-ABE-linker-ARG8 expressed from a nuclear plasmid, and ABE-linker-ARG8 (not ARG8*) integrated into mtDNA (**Figure S9A**). The former showed strong rescue on (-)Arg medium with very high editing (**Figure S9D, E**), indicating that the fusion protein itself is functional. However, the latter failed to rescue growth on (-)Arg and qPCR showed that its transcript levels were 5-fold lower than that of ARG8-only control (**Figure** **S9F**). Thus, it seems that fusing the ABE gene to our reporter in the context of mtDNA impairs its transcription, and possibly its translation as well.

The second design we tested incorporated ABE into a separate copy of the COX2 UTRs (**Figure S9B**). This resulted in a very high rate of recombination since there are three COX2 UTRs present within mtDNA. Cultures from single colonies lacked either ABE or ARG8. A third design used COX3 UTRs to flank the ABE cassette. However, the minimal COX3 promoters and terminators used resulted in no transcription of ABE.

In parallel, we varied the arginine concentration in the growth medium, aiming to create selection conditions more permissive for low-level ARG8 function. While 1/5 of the normal arginine concentration restored ARG8 expression level in the mtDNA-ABE-ARG8 control strain (**Figure S9G**), that did not translate to a difference in growth in media lacking arginine (**Figure S9H**). Hence, while transcript abundance could be manipulated through changes in culture conditions and construct design, the ultimate barrier seemed to be at the level of protein stability or translation efficiency within mitochondria.

**Further discussion.** In contrast to yeast, mammalian cells may lack a natural pathway for import of endogenous nuclear-encoded RNAs. The human mitochondrial genome encodes a complete set of tRNAs and rRNAs to support intra-mitochondrial protein translation, so RNA import is not obligatory as it is in other species (e.g., trypanosomes^1^). Various studies have reported that certain human RNAs, including nuclear-encoded 5S rRNA^2–5^, H1 RNA^6,7^, and 7-2/MRP RNA^8,9^ are imported into mitochondria by human cells. There is little homology among these RNAs and systematic mapping, such as by our previous APEX-seq method^10^, which is performed in living cells with fully intact mitochondria over 1 minute and is less subject to impurities and false positives than mitochondrial fractionation, did not detect any of these or other nuclear-encoded RNAs within human mitochondria. To the best of our knowledge, human mitochondria may not import endogenous RNAs.

Our efforts revealed the difficulty of carefully titrating ABE expression, to both minimize import toxicity and to allow detectable base editing within mitochondria. Additional factors, such as transcriptional tuning and post-transcriptional regulation are likely to be critical in addressing the challenge of MTS-ABE toxicity.

There are many future avenues to explore in the development of CRISPR editing technology for mtDNA. First, we can look to other species and their mechanisms of endogenous RNA import into mitochondria. In trypanosomes, the ATOM protein import complex has been identified as an essential player in delivering tRNAs across the mitochondrial membrane^11^. In potatoes, VDAC can mediate RNA translocation into mitochondria^12^. Recently, researchers found evidence that the human protein ANT2 may act as a bi-directional translocon of RNA into human mitochondria^13^. In yeast, a third tRNA - cytosolic tRNA^Gln^ - has genetic and biochemical evidence of mitochondrial localization but no known mechanism of RNA import^14^. The import of this tRNA appears to not require cytosolic factors or mitochondrial membrane potential^14^. We did include a direct fusion of tRNA^Gln^ to sgRNA in our original screen but did not observe rescue of ARG8^m^* cells.

Second, though a fully genetically-encoded method for CRISPR editing would be ideal, it is worth exploring chemo-genetic approaches to introducing different components of the CRISPR machinery. Direct covalent linking of a peptide mitochondrial targeting sequence to RNA failed to deliver the RNA to mitochondria^15^. Still, liposome-based nanocarriers have shown some success, including mito-Porter (a mitochondria fusogenic liposome)^16^, β-MEND (a liposome with a specific lipid composition to enable mitochondrial entry)^17^ and poly(amidoamine) dendrimer conjugated to TPP^18^, a small molecule capable of entering mitochondria.

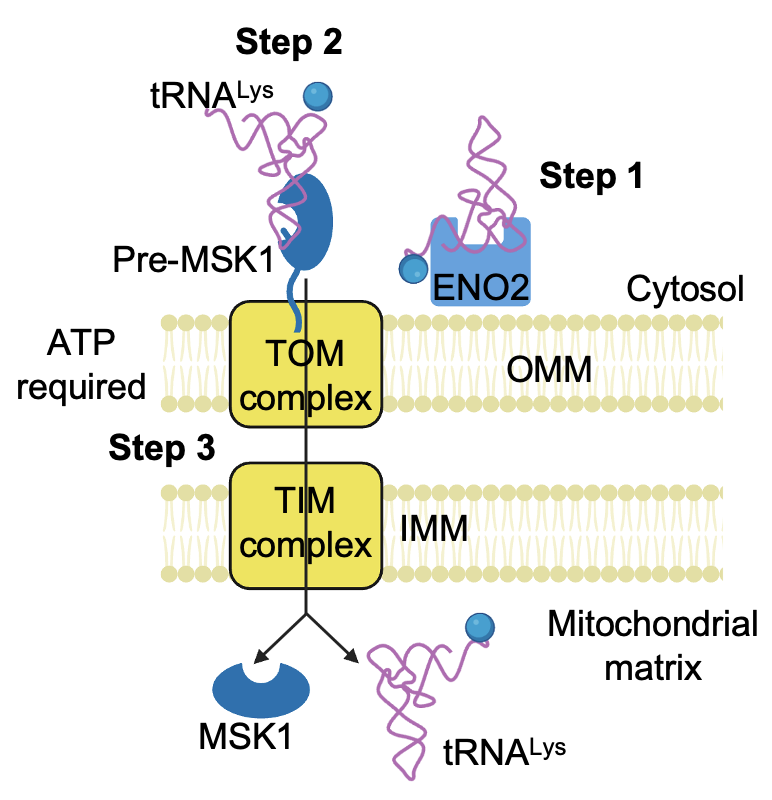

**Supplementary Figure 1. Current model of endogenous tRNA^Lys^ import into yeast mitochondria**^19–23^**.** First, aminoacylated tRNA^Lys^ binds to an enolase, ENO2, which delivers it to the outer mitochondrial membrane (OMM). Second, tRNA^Lys^ binds to the precursor of mitochondrial tRNA synthase, pre-Msk1. Third, the pre-Msk1:tRNA^Lys^ complex is imported into the mitochondrial matrix through the canonical TOM/TIM protein import pathway. ATP is required in the process.

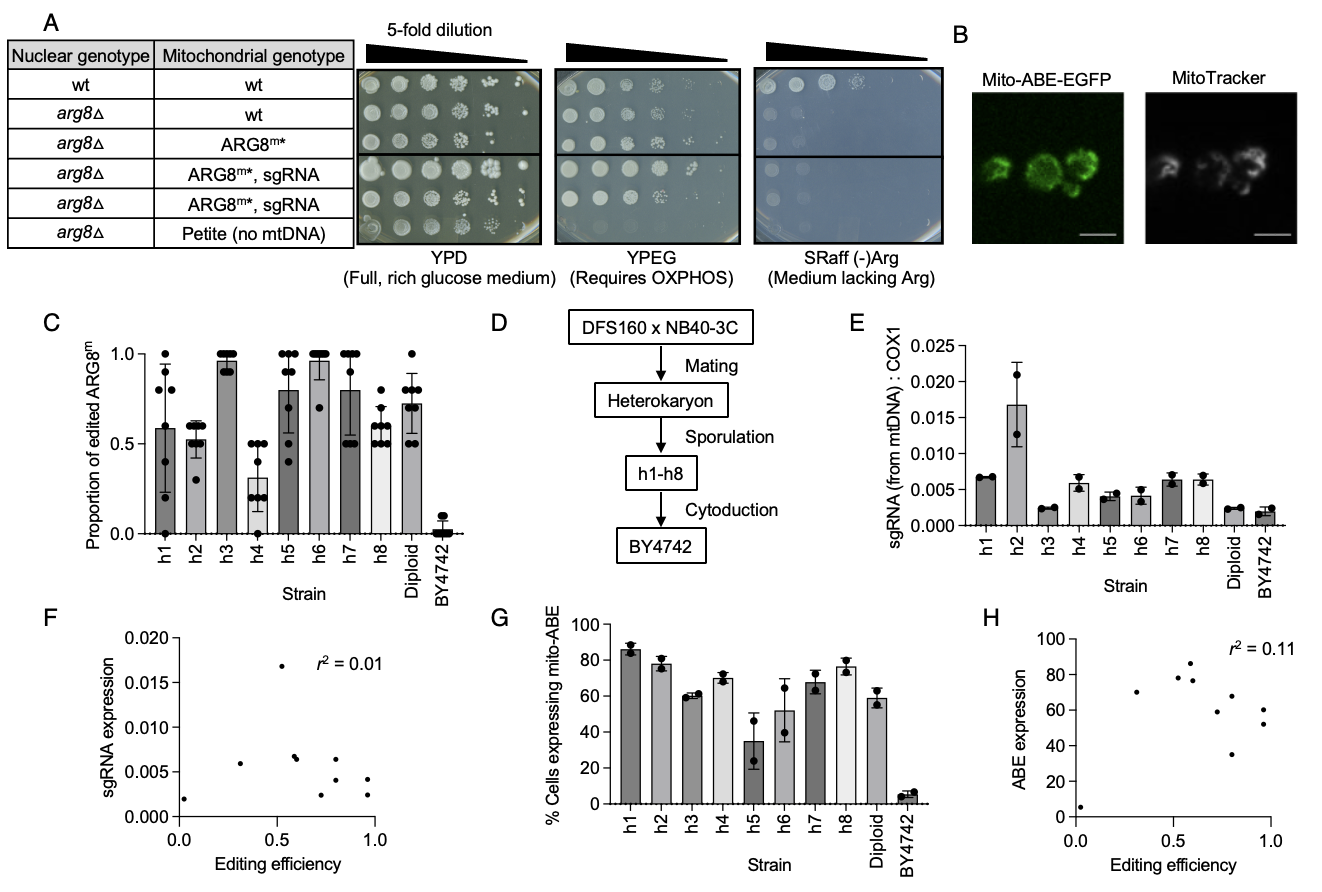

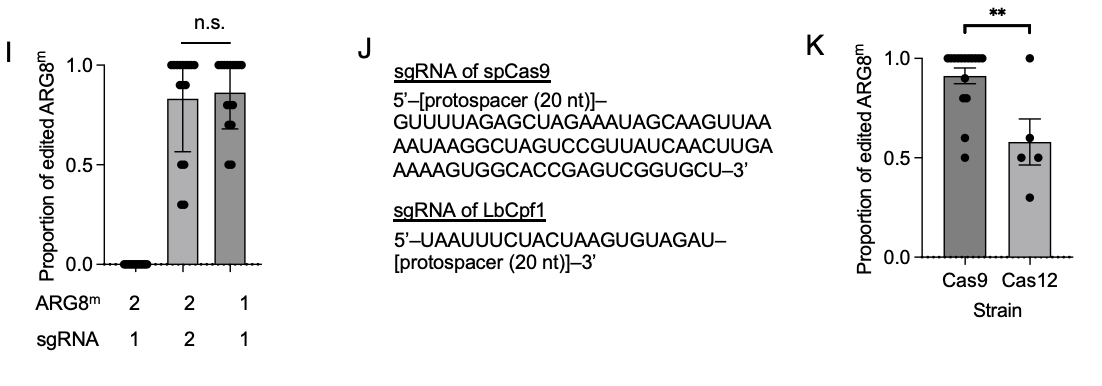

**Supplementary Figure 2. Characterization of BY4742 yeast strain used for mtDNA CRISPR editing assay.** (**A**) Verification of gene gun strains. Saturated yeast cultures were diluted and plated on the indicated media. Full mitochondrial genotype of ARG8^m^* is [COX2 promoter][5’UTR of COX2][ARG8^m^*][3’UTR of COX2]. (**B**) Yeast cells expressing mito-ABE-EGFP and labeled with MitoTracker. Scale bar: 5 µm. (**C**) Proportion of edited ARG8^m^ in colonies transformed with mito-ABE. Eight colonies per strain were directly picked after transforming with mito-ABE. Then the section containing the mutation was amplified and Sanger-sequenced. The proportion of editing is calculated as the height of C peak / height of T peak at the edited nucleotide. (**D**) Flowchart for generation of BY4742 yeast strain. DFS160 is the recipient strain. NB40-3C is the strain containing a mutation in COX2 which allows selection of mitochondrial transformants^1^. Mating of DFS160 and NB40-3C yielded heterokaryons, which were sporulated. h1 to h8 strains were picked from spores and the final BY4742 strain was generated through cytoduction. (**E**) Quantification of sgRNA expression (normalized against mtDNA-encoded COX1) in ten strains. All express sgRNA from mtDNA. (**F**) Scatter plot showing sgRNA expression (as measured in (E)) as a function of editing efficiency (as measured in (C)). (**G**) Quantification of percentage of cells expressing mito-ABE-EGFP, measured by flow cytometry. (**H**) Scatter plot ABE expression (as measured as percentage of cells expressing mito-ABE-EGFP in (G)) as a function of editing efficiency (as measured in (C)). (**I**) Comparison of editing efficiencies with two ARG8^m^* reporters. Reporter 1 harbors the Q79* mutation and reporter 2 harbors the W168* mutation. sgRNA1 targets reporter 1 and sgRNA2 targets reporter 2. Editing efficiency was calculated as in (C). ns, not significant. (**J**) sgRNA sequences used with spCas9 and LbCpf1. (**K**) Comparison of editing efficiencies with Cas9 and Cas12-based ABE, calculated as in (C). ***p*<0.01.

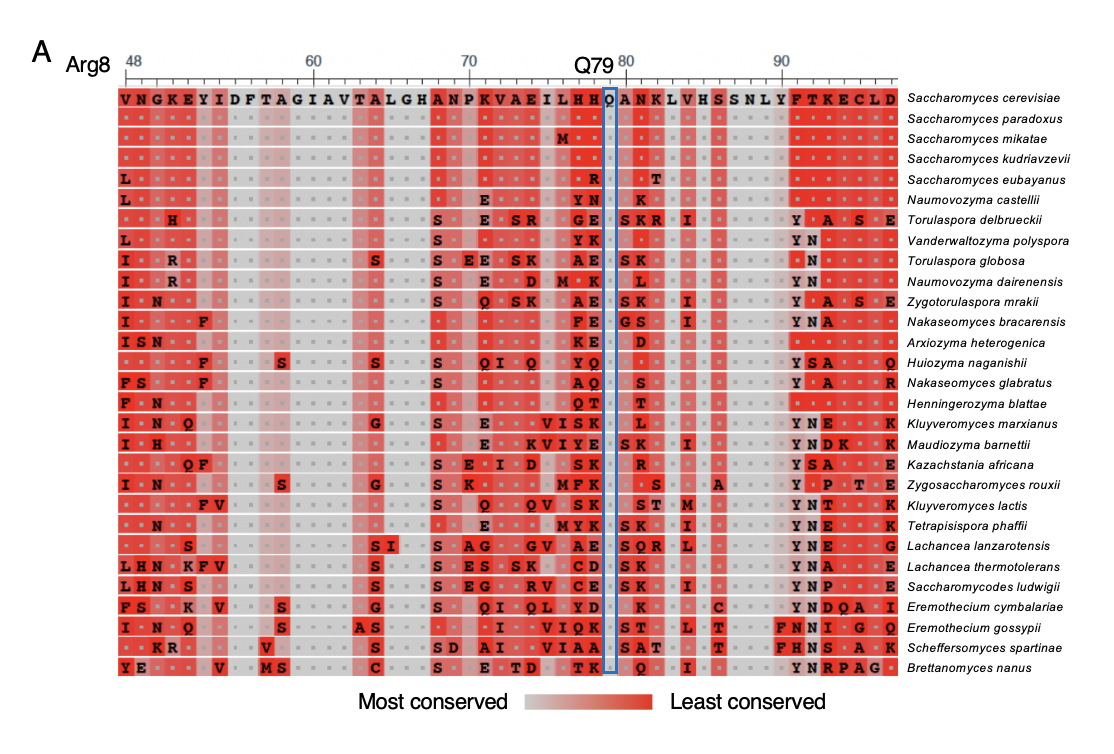

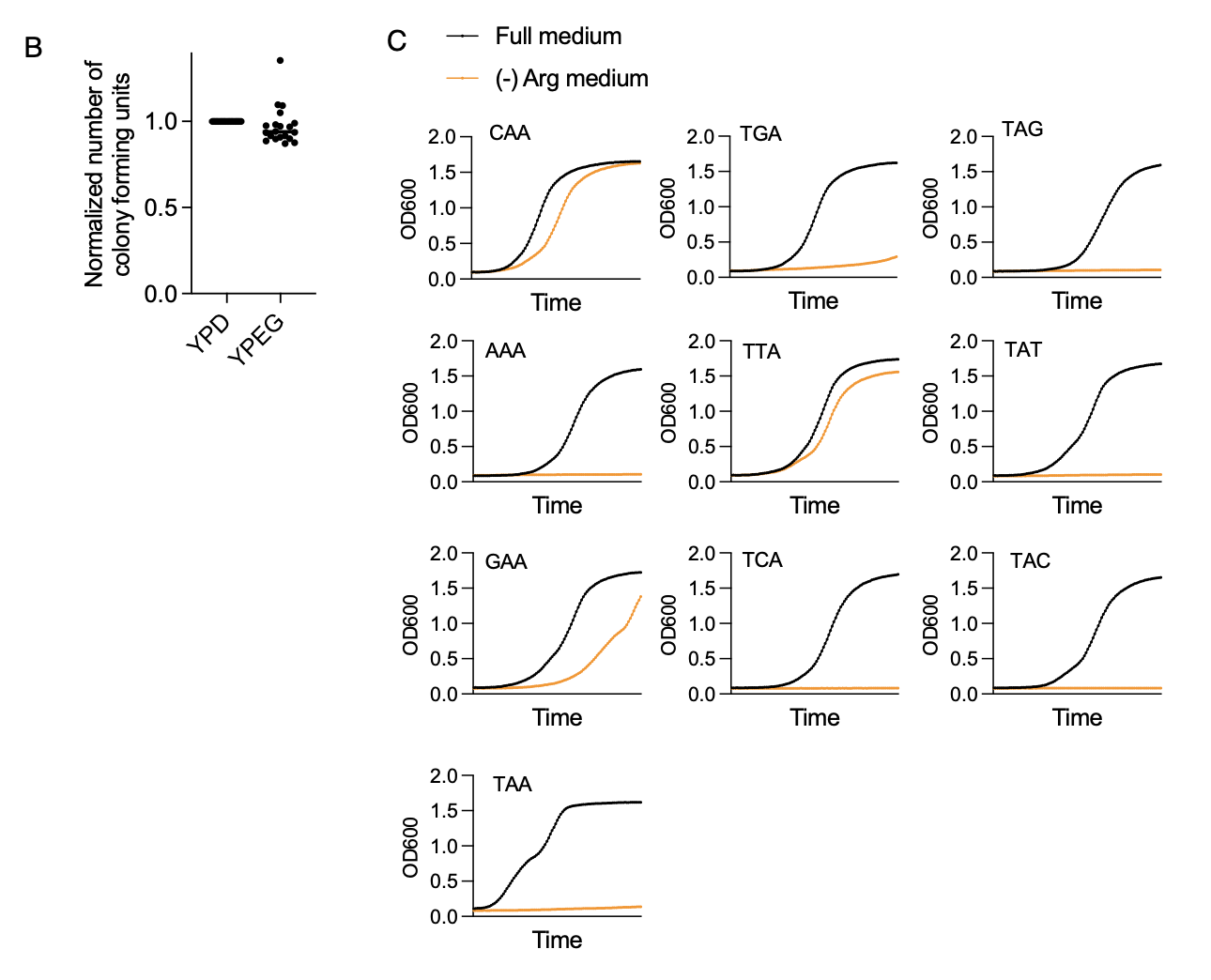

**Supplementary Figure 3: Further characterization of ARG8m.** (**A**) Homologous Arg8 proteins across species were identified using BLASTp analysis. Aligned sequences showed high conservation at Q79 (highlighted), indicating strong evolutionary pressure to preserve this residue. (**B**) Number of colony forming units (cfu) on YPD (full, rich glucose medium) and YPEG (ethanol medium requiring active respiration) for yeasts expressing ARG8m plated to test spontaneous reversion. Cfu on YPEG were normalized to the corresponding cfu number on YPD. (**C**) Growth curves of ARG8m* and all possible single mutants in full medium and (-) Arg medium. Optical density at 600 nm was measured every 15 minutes over a total of 48 hours.

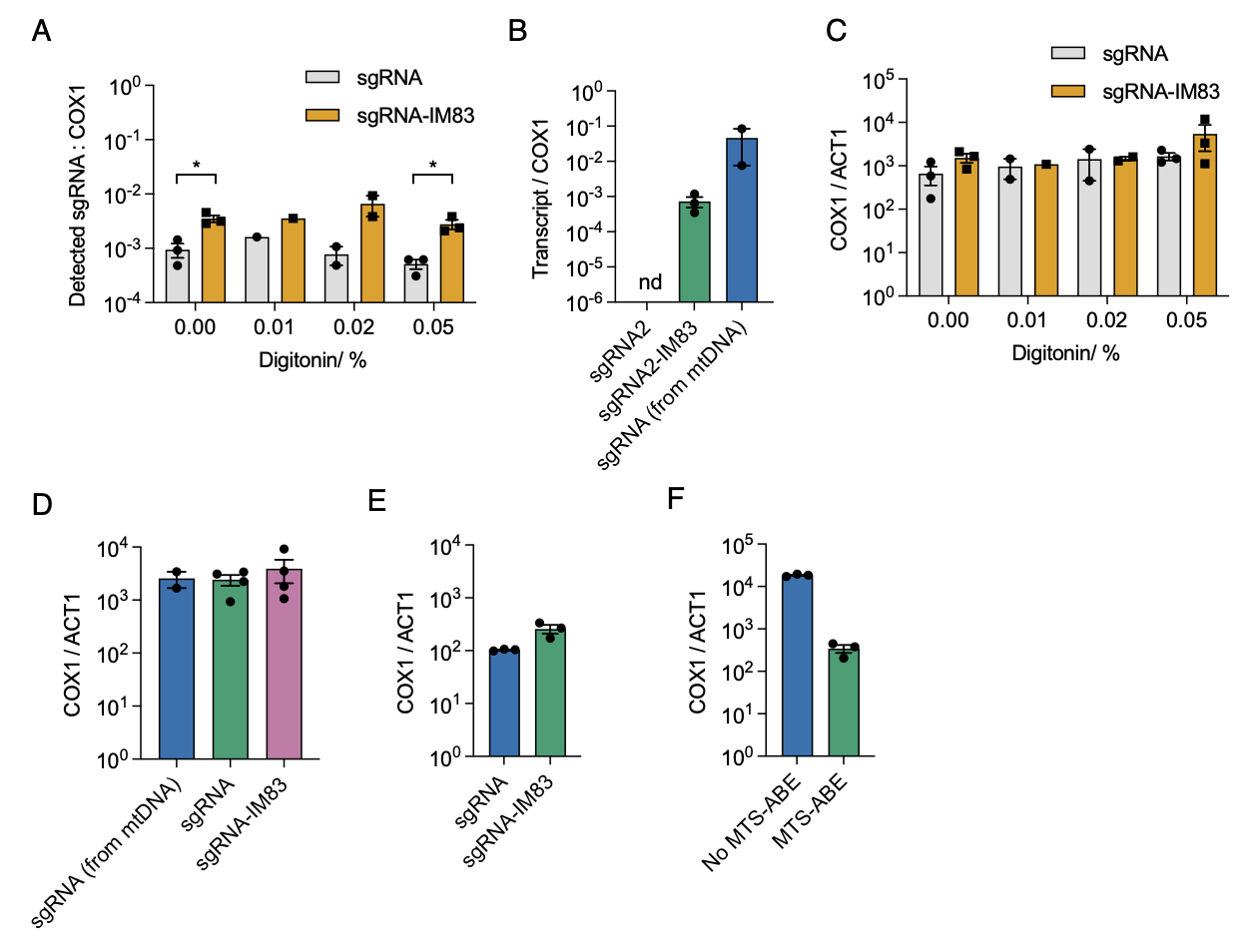

**Supplementary Figure 4. Characterization of mitoplasts used for sgRNA import assay**. (**A**) Same as **Figure 3B** except mitochondria were purified further via digitonin digestion of the OMM, to produce mitoplasts. sgRNA-IM83 was consistently detected at higher levels than sgRNA lacking the import motif. Each condition was repeated 1-3 times. *: *p* < 0.05 (left: *p* = 0.011, right: *p* = 0.015). (**B**) RT-qPCR analysis of purified mitochondria with sgRNA2 or sgRNA2-IM83 expressed from a plasmid, or sgRNA expressed from mtDNA. sgRNA2 is the same as sgRNA1 except that its protospacer does not target ARG8^m^. sgRNA2/ ACT1 was not detected in 35 PCR cycles. Each condition was repeated 2-3 times. (**C**) RT-qPCR analysis of purified mitoplasts from (A). COX1 is encoded in mtDNA, while ACT1 is encoded in the nuclear genome. The ratio of COX1 to ACT1 abundance reflects mitochondrial purity. (**D**) RT-qPCR analysis of purified mitochondria used in indicated samples from **Figure 3B**. (**E**) RT-qPCR analysis of purified mitochondria used in indicated samples from **Figure 3C**. (**F**) RT-qPCR analysis of purified mitochondria used in indicated samples from **Figure 3E** (no MTS-ABE) and **Figure 3F** (with MTS-ABE).

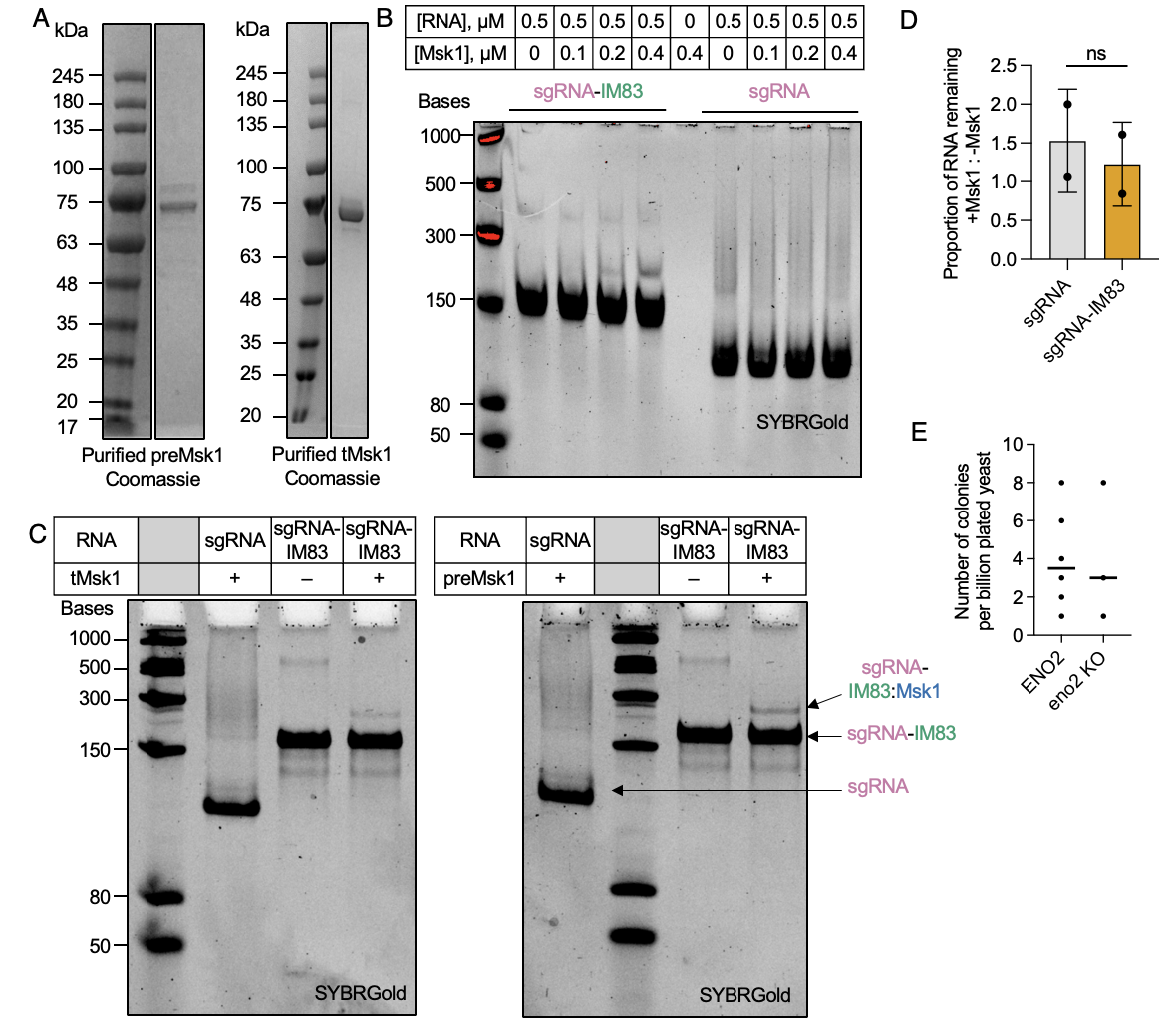

**Supplementary Figure 5. Pre-Msk1 is responsible for sgRNA-IM83 entry into mitochondria. Related to Figure 4.** (**A**) SDS-PAGE analysis of purified recombinant preMsk1 (66.1 kDa, left) and tMsk1 (64 kDa, right). (**B**) Uncropped gel with ladder for **Figure 4D**. (**C**) Gel shift assay showing that both full-length preMsk1 and tMsk1 proteins bind to sgRNA-IM83, but not to sgRNA. RNA and protein concentrations used are 0.3 μM. (**D**) Msk1 did not protect sgRNA-IM83 from RNAse digestion more than sgRNA. The bar graph compares the amounts of sgRNA or sgRNA-IM83 left after limited RNAse digest in the presence and absence of Msk1. ns, not significant. (**E**) Editing efficiency of sgRNA-IM83 expressed from nuclear plasmids in yeast with or without *ENO2*. Experiment is the same as **Figure 2C**.

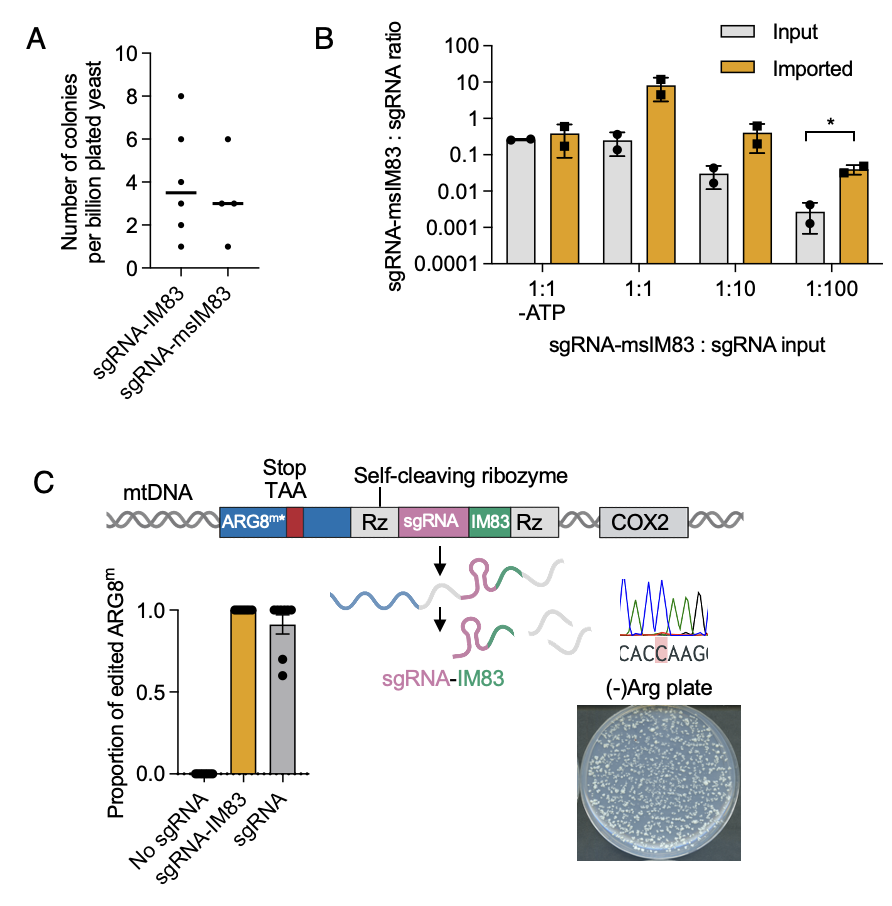

**Supplementary Figure 6. Further characterization of sgRNA-IM83 and variants.** (**A**) Editing efficiency of sgRNA-IM83 and sgRNA-msIM83 expressed from nuclear plasmids. Experiment is the same as **Figure 2C**. (**B**) Same as **Figure 3E** but with sgRNA-msIM83. Data are from two biological replicates. **p*< 0.05 from a two-tailed student’s t-test. (**C**) Control experiment showing that IM83 did not decrease editing efficiency when expressed as sgRNA-IM83 from mtDNA. Top: Same as **Figure 1A** except sgRNA was replaced by sgRNA-IM83. Three colonies were picked from (-)Arg plates and all contained the desired edit. The experiment was repeated 3 times with similar results. Bottom left: comparison of editing efficiencies with no sgRNA/ sgRNA/ sgRNA-IM83 expressed from mtDNA. Quantitation as in **Figure S2C**.

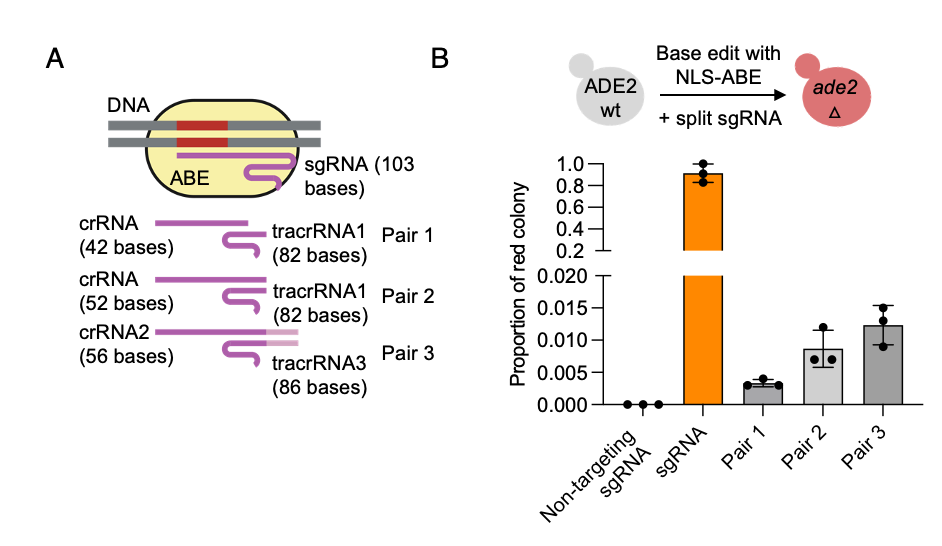

**Supplementary Figure 7.** **Splitting sgRNA to reduce the length required for import.** (**A**) Split sgRNA design. sgRNA was split into crRNA and tracrRNA portions with various annealing lengths (e.g., 19 nt for Pair 1 and 29 nt for Pair 3). TracrRNA was expressed from mtDNA while the variable crRNA was imported via fusion to IM83. We experimentally tested the import of Pair 1 and 2 crRNAs, but no editing was seen. (**B**) Testing split sgRNA designs for nuclear editing. Split guides targeting the adenine biosynthesis gene ADE2 were expressed in yeast with nuclear-targeted ABE. Base editing alters ADE2’s start codon (ATG to GTG), stopping ADE2 translation and producing red colonies. All split pairs of crRNA/ tracrRNA reduced the editing efficiency of nuclear ADE2 by nuclear ABE. NLS, nuclear localization signal.

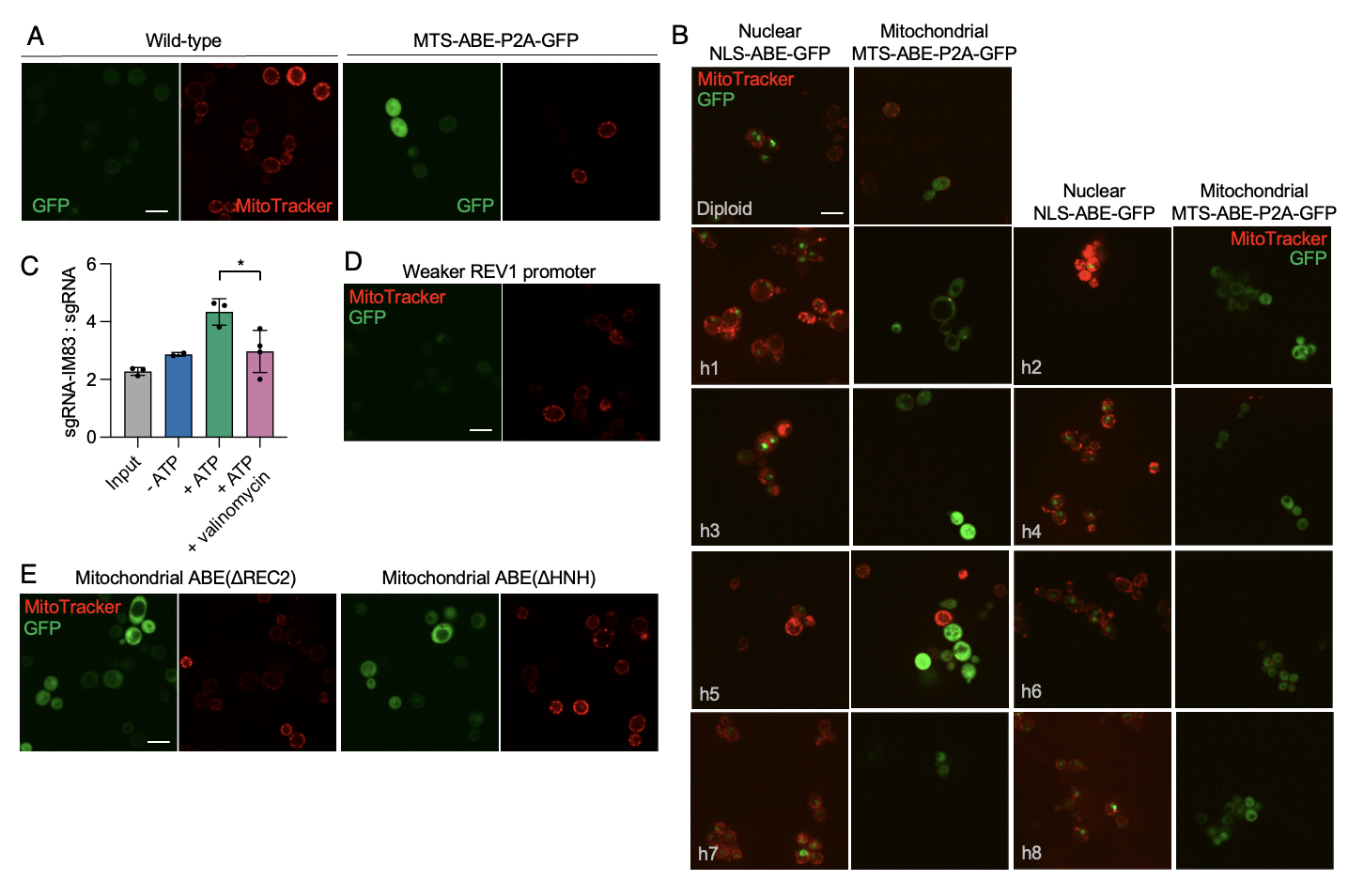

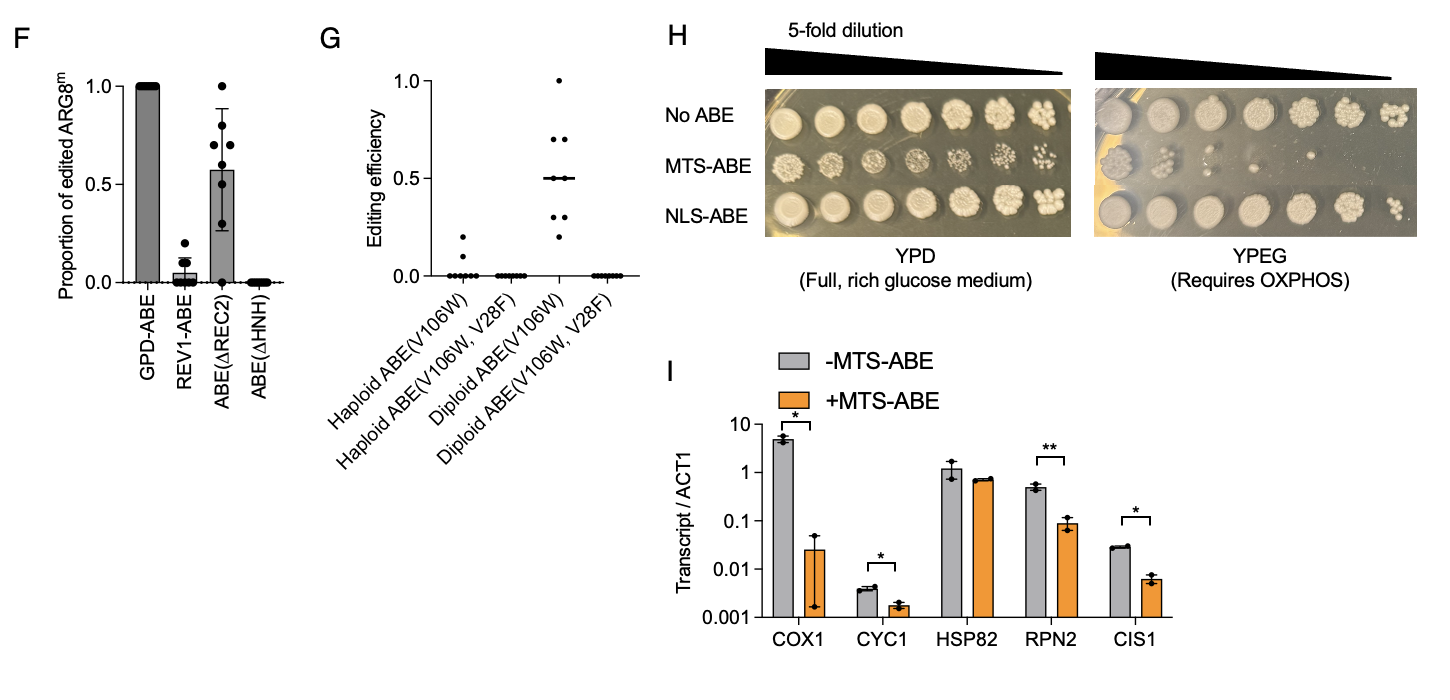

**Supplementary Figure 8.** **Targeting ABE to mitochondria reduces mitochondrial membrane potential.** (**A**) Confocal fluorescence imaging of wild-type yeast cells and yeast cells expressing mito-ABE-P2A-GFP. Yeast were stained with MitoTracker. Scale bar: 5 µm. (**B**) Diploid and haploid (strains h1-h8) yeast cells expressing nuclear-targeted ABE (NLS-ABE-GFP) or mitochondrial-targeted ABE (MTS-ABE-P2A-GFP) under a strong GPD promoter and labeled with MitoTracker. Scale bar, 5 µm. (**C**) Results from the in vitro sgRNA import assay in **Figure 3D**, with valinomycin to disrupt mitochondrial membrane potential. Data from two to three biological replicates per condition. (**D**) Yeast cells expressing MTS-ABE-P2A-GFP under a weak REV1 promoter and labeled with MitoTracker. Scale bar: 5 µm. (**E**) Yeast cells expressing truncated MTS-ABE-P2A-GFP constructs under a strong GPD promoter and labeled with MitoTracker. Scale bar: 5 µm. (**F**) Proportion of edited mitochondrial ARG8 in colonies transformed with ABE constructs from (D) and (E). sgRNA was expressed from mtDNA. Colonies were directly picked after transformation, then the edited region was amplified and sequenced. The proportion of editing is calculated as the ratio of C peak to T peak height in the Sanger sequencing chromatogram. Data from 8 colonies per condition. (**G**) Editing efficiencies of MTS-ABE(V106W) and MTS-ABE(V106W, V28F) expressed from a nuclear plasmid on haploid (BY4742 nuclear background) or diploid yeasts expressing sgRNA from mtDNA. Quantification as in **Figure S2C**. (**H**) Phenotyping of yeast expressing no ABE, MTS-ABE, and NLS-ABE on YPD (full, rich glucose medium) and YPEG (ethanol medium requiring active respiration). Saturated yeast cultures were diluted and plated on the indicated medium. (**I**) RT-qPCR analysis of OXPHOS components (COX1, CYC1) and reported upregulated genes of mitochondrial protein import stress (HSP82, RPN2, CIS1), in yeast with or without MTS-ABE expression. * p< 0.05, ** p< 0.01

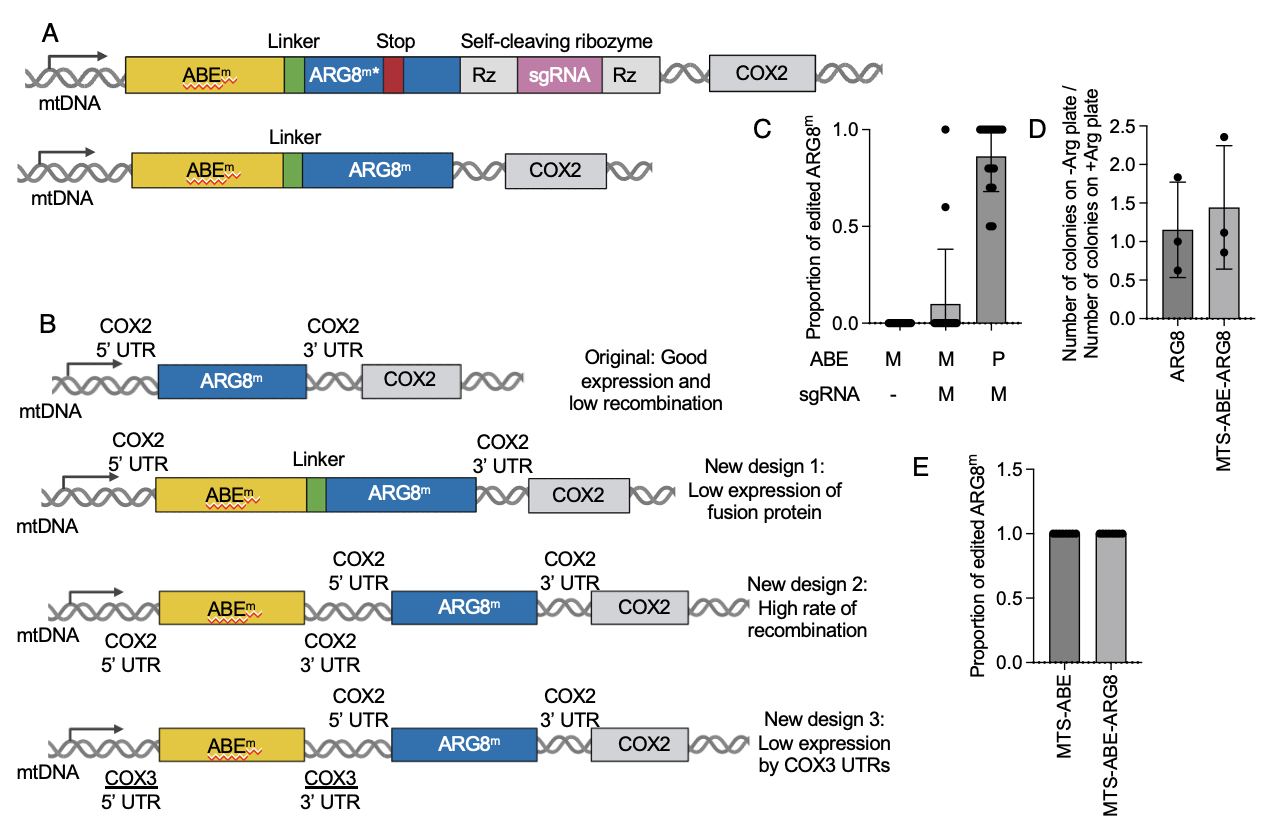

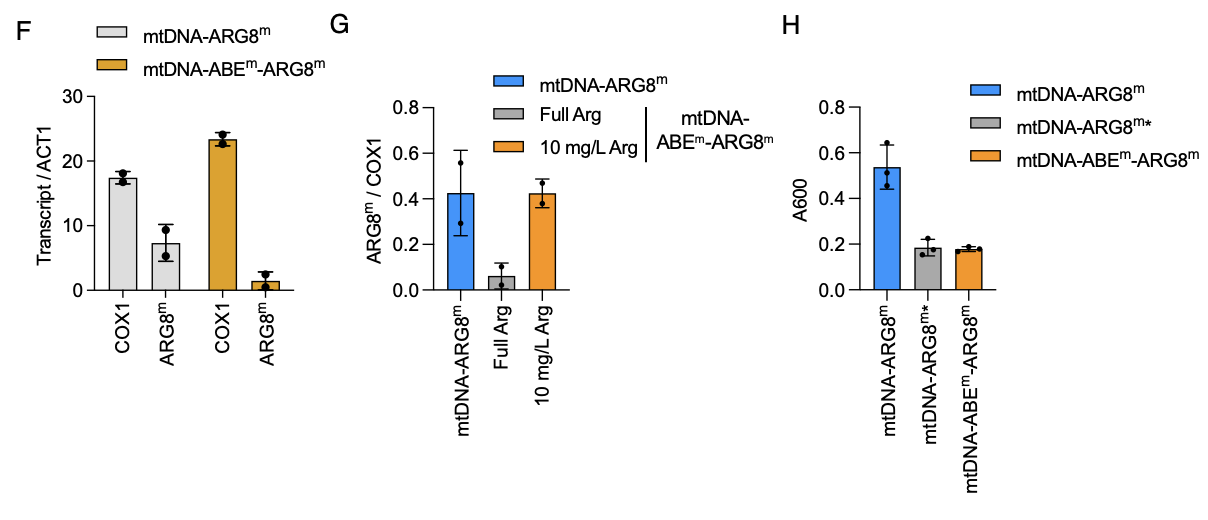

**Supplementary Figure 9. Expression of ABE from mtDNA.** (**A**) Top: Design of ABE^m^-linker-ARG8^m^* fusion protein. The strain is identical to **Figure 1A** except that ABE is fused to the N terminus of ARG8^m^* with a 6-amino acid flexible linker. Bottom: positive control construct with wild-type ARG8^m^ lacking a stop codon. (**B**) The original design and new designs to improve mtDNA expression of ABE^m^ and ARG8^m^. New design 1 expresses a fusion protein under COX2 UTRs. New design 2 places ABE^m^ and ARG8^m^ under two separate COX2 promoters and UTRs. New design 3 places ABE^m^ and ARG8^m^ under COX3 and COX2 promoters and UTRs respectively. (**C**) Proportion of edited ARG8^m^ in colonies transformed with construct in (A). M: ABE or sgRNA expressed from mtDNA. P: mitochondria-targeting ABE expressed from plasmid. Column 1: bottom construct in S9A; column 2: top construct in S9A; column 3: original reporter with mito-ABE expressed from a nuclear plasmid as in **Figure 1A**. Colonies were directly picked after transformation, then sequenced. The proportion of editing is calculated as the ratio of C peak to T peak height in the Sanger sequencing chromatogram, as in **Figure S2C**. (**D**) Data from 16 colonies per condition. The ratio of colonies on -Arg versus +Arg plates, for yeast strains with wild-type ARG8 knocked out and transformed with nuclear plasmids expressing wildtype ARG8 or a MTS-ABE-ARG8 fusion. Data from 3 biological replicates. (**E**) Proportion of edited ARG8^m^ in colonies transformed with MTS-ABE or MTS-ABE-ARG8. Analysis performed as in (C). Data from 8 colonies each. (**F**) RT-qPCR analysis of yeast expressing only ARG8^m^ from mtDNA and yeast expressing ABE^m^-ARG8^m^ fusion from mtDNA. COX1 is a mtDNA-encoded mRNA. Data from 2 biological replicates. (**G**) Culturing mtDNA-expressed ABE^m^-ARG8^m^ fusion with 10 mg/L arginine improved ARG8^m^ expression, as measured by qPCR. Transcript level of ARG8^m^ was normalized against a mitochondrial mRNA, COX1. Data from 2 biological replicates. (**H**) Absorbance (600 nm) of yeast culture after inoculating from a YPEG plate and growing overnight expressing only ARG8^m^, non-functional ARG8^m^* and ABE^m^-ARG8^m^ fusion from mtDNA in media with 10 mg/L arginine. Data from 3 biological replicates.

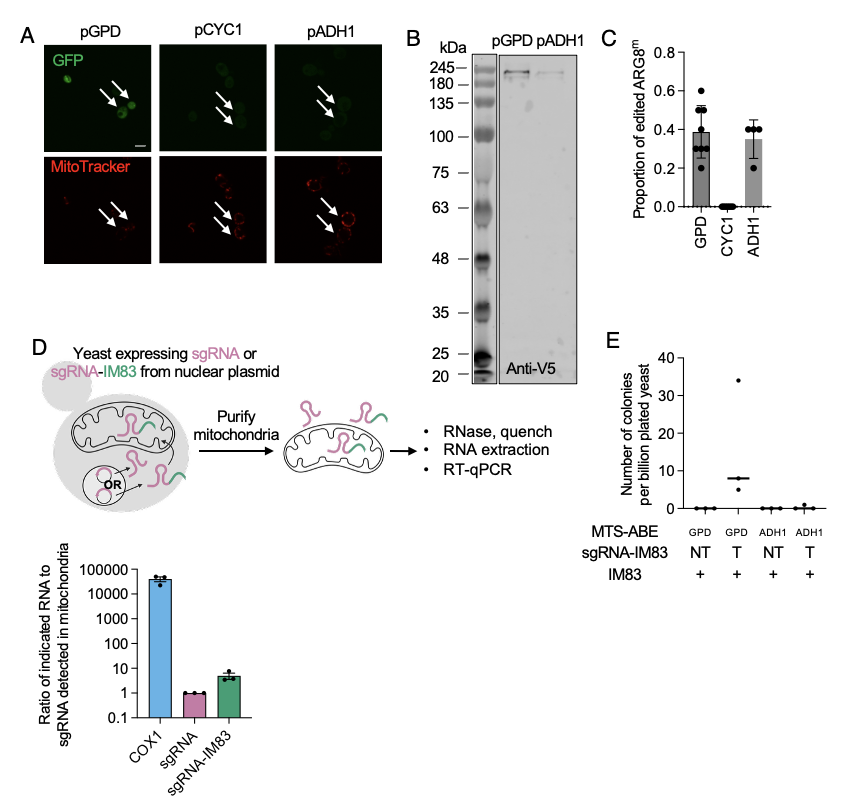

**Supplementary Figure 10 Reduction of MTS-ABE expression to reduce mitochondrial toxicity.** (**A**) Confocal fluorescence imaging of yeast cells expressing mito-ABE-P2A-GFP under a GPD, ADH1 or CYC1 promoter. Yeast were stained with MitoTracker. Scale bar: 5 µm. (**B**) Western blot analysis of lysates from yeast cells expressing mito-ABE-V5-P2A-GFP under a GPD or ADH1 promoter.  (**C**) Proportion of edited ARG8^m^ in colonies transformed with mito-ABE expressed under GPD, ADH or CYC promoter. 4-8 colonies per strain were directly picked after transforming with mito-ABE. Then the section containing the mutation was amplified and Sanger-sequenced. The proportion of editing is calculated as the height of C peak / height of the T peak at the edited nucleotide. (**D**) Schematic for detecting sgRNA mitochondrial import. Mitochondria are purified and then analyzed for RNA content by RT-qPCR and results from this assay. Yeast cells expressing mito-ABE under ADH1 with sgRNA and sgRNA-IM83 expressed from a 2u plasmid. Abundance of each transcript is normalized to that of ACT1, a cytosolic mRNA, then compared to sgRNA. COX1 is a positive control mitochondrial mRNA. Each condition was repeated 3 times. (**E**) Editing of ARG8m* reporter by mito-ABE expressed from GPD or ADH1 promoter and targeting/non-targeting sgRNA-IM83 expressed from a 2u plasmid. Surviving colonies on (-)Arg plates were counted after performing the assay in **Figure 2A**.

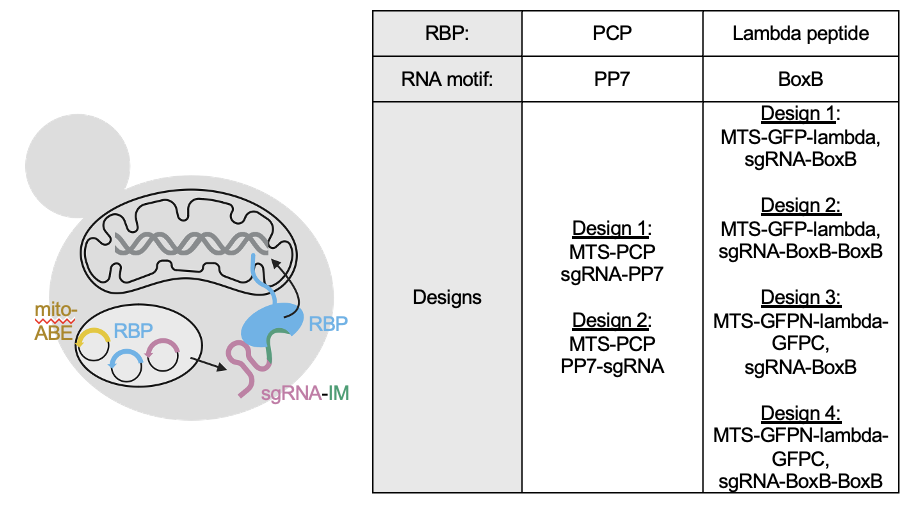

**Supplementary Figure 11.** **Testing RBPs for** **sgRNA import into mitochondria.** Testing RNA binding proteins (RBPs) for sgRNA import into mitochondria. Two RBPs were tested (PCP and lambda peptide) with 2-4 designs each. All failed to give ARG8^m^ editing.

**Supporting Table 1. List of plasmids used in this work.**

| Plasmid name | Description | Reference |
| --- | --- | --- |
| MTS-ABE8e-nCas9-EGFP | pRS415-GPD-SU9MTS-ABE8e(V106W)(nCas9)-V5-EGFP  Expression of SU9-MTS-ABE8e(V106W)(nCas9)-GFP in *S. cerevisiae*.  Used in Figures S2B. | This study |
| MTS-ABE8e-nCas9-P2A-EGFP | pRS415-GPD-SU9MTS-ABE8e(V106W)(nCas9)-V5-P2A-EGFP  Expression of SU9-MTS-ABE8e(V106W) with a GFP expression marker in *S. cerevisiae*.  Used in Figures 1B, 1C, 3C, 3F, S2C, S2G, S2I, S2K. | This study |
| MTS-ABEmax-nCas9-P2A-EGFP | pRS415-GPD-SU9MTS-ABEmax(nCas9)-V5-P2A-EGFP  Expression of SU9-MTS-ABEmax with a GFP expression marker in *S. cerevisiae*.  Used in Figures 2C, | This study |
| Mito-Cas12 | pRS415-GPD-SU9MTS-ABE8e(V106W)(LbCpf1)-V5-P2A-EGFP  Expression of SU9-MTS-ABE8e(V106W)(LbCpf1) with a GFP expression marker in *S. cerevisiae*.  Used in Figures 1C, S2K. | This study |
| sgRNA-IM | pRS426-SNR52-sgRNA-IM  Expression of sgRNA-IM in *S. cerevisiae*.  Used in Figures 2, 3, S4. | This study |
| sgRNA-msIM83 | pRS426-SNR52-sgRNA-msIM83  Expression of sgRNA-msIM83 in *S. cerevisiae*.  Used in Figures S6A, S6B. | This study |
| sgRNA(nt)-IM | pRS426-SNR52-sgRNA(nt)-IM  Expression of non-targeting sgRNA-IM in *S. cerevisiae*.  Used in Figure S2. | This study |
| Msk1 | MSK1-His6  Expression and purification of His6-tagged Msk1 protein from *E. coli*.  Used in Figures 4, S5. | This study |
| tMsk1 | MSK1(Δ1-26)-His6  Expression and purification of His6-tagged Msk1 protein from *E. coli*.  Used in Figures 4, S5. | This study |
| NLS-ABE-GFP | pRS415-GPD-NLS-ABE8e(V106W)-V5-EGFP  Expression of nuclear-localized ABE8e(V106W)(nCas9)-GFP in *S. cerevisiae*.  Used in Figure S8B. | This study |
| REV1-MTS-ABE8e-P2A-EGFP | pRS415-REV1-SU9MTS-ABE8e(V106W)-V5-EGFP  Expression of MTS-ABE8e(V106W)(nCas9) with a GFP expression marker under a low-strength promoter in *S. cerevisiae*.  Used in Figures S8E, S8G. | This study |
| MTS-ABE8e(∆REC2)- EGFP | pRS415-GPD-SU9MTS-ABE8e(V106W, ∆REC2)-V5-EGFP  Expression of MTS-ABE8e(V106W)(nCas9∆REC2) with a GFP expression marker in *S. cerevisiae*.  Used in Figures S8F, S8G. | This study |
| MTS-ABE8e(∆HNH)- EGFP | pRS415-GPD-SU9MTS-ABE8e(V106W, ∆HNH)-V5-EGFP  Expression of MTS-ABE8e(V106W)(nCas9∆HNH) with a GFP expression marker in *S. cerevisiae*.  Used in Figures S8F, S8G. | This study |
| MTS-ABE-ARG8 | pRS415-GPD-SU9MTS-ABE8e-linker-ARG8-V5  Expression of V5-tagged MTS-ABE8e(V106W)(nCas9)-linker-ARG8 in *S. cerevisiae*.  Used in Figures S10C, S10D. | This study |
| crRNA(1-3) | pRS426-SNR52-crRNA  Expression of crRNA in *S. cerevisiae*.  Used in Figure S11. | This study |
| tracrRNA | pRS426-SNR52-tracrRNA  Expression of tracrRNA in *S. cerevisiae*.  Used in Figure S11. | This study |
| sgRNA-PP7 | pRS426-SNR52-sgRNA-PP7  Expression of PP7-tagged sgRNA in *S. cerevisiae*.  Used in Figure S11. | This study |
| PP7-sgRNA | pRS426-SNR52-PP7-sgRNA  Expression of PP7-tagged sgRNA in *S. cerevisiae*.  Used in Figure S11. | This study |
| MTS-PCP | pRS415-GPD-COX4MTS-PCP-V5-EGFP  Expression of MTS-PCP-V5-GFP in *S. cerevisiae*.  Used in Figure S11. | This study |
| MTS-GFP-lambda | pRS415-GPD-COX4MTS-GFP-lambda peptide-V5  Expression of MTS-GFP with lambda peptide at the C-terminus in *S. cerevisiae*.  Used in Figure S11. | This study |
| MTS-GFPN-lambda-GFPC | pRS415-GPD-COX4MTS-GFPN-lambda peptide-GFPC-V5-EGFP  Expression of MTS-GFP with an internal lambda peptide in *S. cerevisiae*.  Used in Figure S11. | This study |
| sgRNA-BoxB | pRS426-SNR52-sgRNA-BoxB  Expression of BoxB-tagged sgRNA in *S. cerevisiae*.  Used in Figure S11. | This study |
| sgRNA-2x BoxB | pRS426-SNR52-sgRNA-BoxB-BoxB  Expression of double BoxB-tagged sgRNA in *S. cerevisiae*.  Used in Figure S11. | This study |

**Supporting Table 2. List of yeast strains used in this work.**

| Strain name | Nuclear genotype | Mitochondrial genotype | Reference |
| --- | --- | --- | --- |
| DFS160 | MAT α Δleu2 ura3-52 ade2-101 arg8::URA3 kar1-1 | ρ^0^ | ^1^ |
| NB40-3C | MAT a lys2 leu2-3,112 ura3-52 his3::HinDIII arg8::hisG | COX2:cox2-62 | ^1^ |
| Diploid ARG8^m^* | MAT a/α lys2/LYS2 Δleu2/leu2-3,112 URA3/ura3-52 ADE2/ade2-101 his3::HinDIII arg8::hisG/ arg8::URA3 KAR1/kar1-1 | COX2::ARG8^m^*-COX2 | This study |
| Diploid ARG8^m^*-sgRNA | MAT a/α lys2/LYS2 Δleu2/leu2-3,112 URA3/ura3-52 ADE2/ade2-101 his3::HinDIII arg8::hisG/ arg8::URA3 KAR1/kar1-1 | COX2::ARG8^m^*-HH-sgRNA(Arg8^m^)-HDV-COX2 | This study |
| Diploid ARG8^m^*-sgRNA(Cas12) | MAT a/α lys2/LYS2 Δleu2/leu2-3,112 URA3/ura3-52 ADE2/ade2-101 his3::HinDIII arg8::hisG/ arg8::URA3 KAR1/kar1-1 | COX2::ARG8^m^*-HH-sgRNA(Arg8^m^, Cas12)-HDV-COX2 | This study |
| Haploid ARG8^m^* | MATα his3Δ1, leu2Δ0, lys2Δ0, ura3Δ0, arg8::KanMX | COX2::ARG8^m^*-COX2 | This study |
| Haploid ARG8^m^*-sgRNA | MATα his3Δ1, leu2Δ0, lys2Δ0, ura3Δ0, arg8::KanMX | COX2::ARG8^m^*-HH-sgRNA(Arg8^m^)-HDV-COX2 | This study |
| Haploid ARG8^m^ | MATα his3Δ1, leu2Δ0, lys2Δ0, ura3Δ0, arg8::KanMX | COX2::ARG8^m^-COX2 | This study |
| Petite haploid ARG8^m^* | MATα his3Δ1, leu2Δ0, lys2Δ0, ura3Δ0, arg8::KanMX | ρ^0^ | This study |
| Petite haploid ARG8^m^ | MATα his3Δ1, leu2Δ0, lys2Δ0, ura3Δ0, arg8::KanMX | ρ^0^ | This study |
| *msk1Δ* | MATα his3Δ1, leu2Δ0, lys2Δ0, ura3Δ0, arg8::KanMX, msk1::HIS3 | COX2::ARG8^m^*-COX2 | This study |
| Diploid ARG8^m^*-sgRNA-IM83 | MAT a/α lys2/LYS2 Δleu2/leu2-3,112 URA3/ura3-52 ADE2/ade2-101 his3::HinDIII arg8::hisG/ arg8::URA3 KAR1/kar1-1 | COX2::ARG8^m^*-HH-sgRNA(Arg8^m^)-IM83-HDV-COX2 | This study |
| Diploid ABE^m^-ARG8^m^*-sgRNA | MAT a/α lys2/LYS2 Δleu2/leu2-3,112 URA3/ura3-52 ADE2/ade2-101 his3::HinDIII arg8::hisG/ arg8::URA3 KAR1/kar1-1 | COX2::ABE^m^-linker-ARG8^m^*-HH-sgRNA(Arg8^m^)-HDV-COX2 | This study |
| Diploid ABE^m^-ARG8^m^ | MAT a/α lys2/LYS2 Δleu2/leu2-3,112 URA3/ura3-52 ADE2/ade2-101 his3::HinDIII arg8::hisG/ arg8::URA3 KAR1/kar1-1 | COX2::ABE^m^-linker-ARG8^m^-COX2 | This study |
| Diploid separate ABE^m^ ARG8^m^ | MAT a/α lys2/LYS2 Δleu2/leu2-3,112 URA3/ura3-52 ADE2/ade2-101 his3::HinDIII arg8::hisG/ arg8::URA3 KAR1/kar1-1 | COX2::COX2 5’UTR - ABE^m^- COX2 3’UTR - COX2 5’UTR – ARG8^m^- COX2 3’UTR -COX2 | This study |
| Diploid separate COX3 ABE^m^ ARG8^m^ | MAT a/α lys2/LYS2 Δleu2/leu2-3,112 URA3/ura3-52 ADE2/ade2-101 his3::HinDIII arg8::hisG/ arg8::URA3 KAR1/kar1-1 | COX2::COX3 5’UTR - ABE^m^- COX3 3’UTR - COX2 5’UTR – ARG8^m^- COX2 3’UTR -COX2 | This study |

**Supplementary Methods**

**Phenotyping of mitochondrial transformants**. Related to Figure S2A. Phenotyping of mitochondrial transformants was performed by preparing overnight saturated yeast cultures with defined nuclear and mitochondrial genotypes. Cultures were serially diluted (five-fold), spotted onto respective media, and incubated at 30 °C for 2 days prior to imaging.

**Generation of diploid and haploid strains**. Related to Figure S2D. The vector used for biolistic transformation, pPT24*-ARG8^m^*, contains *ARG8^m^** (together with a sgRNA expression cassette in some strains) flanked by the 5' and 3'UTR of COX2 and the complete COX2 gene further downstream (see sequence map in supplementary information). pPT24*-ARG8^m^* was integrated into the mitochondrial genome via biolistic transformation following this detailed protocol^4^. In summary, DFS160 (a gift from Dr Thomas Fox) was cultured in YPR (20 g/L peptone, 10 g/L yeast extract, 20 g/L raffinose) for 3 days to saturation. 50 ml culture was concentrated to a concentration of 5 x 10^9^ yeast / ml. 0.1 ml of this concentrated culture was spread onto SD-Leu-sorbitol plates (0.77 g/L CSM-Leu, 6.7 g/L yeast nitrogen base, 50 g/L glucose, 1 M sorbitol, 100 mg/L adenine, 30 g/L bacto agar). For 4 shots, 10 ug of Leu carrier plasmid and the 60 μg of the appropriate pPT24 plasmid were mixed and precipitated onto 200 μl 70% ethanol washed tungsten beads (60 mg/ ml in 50% glycerol), then with 8 μl of 1 M spermidine and 20 μl of 2.5 M ice-cold CaCl_2_. This mixture was incubated on ice with occasional shaking for 10 minutes. The mixture was washed with cold ethanol at least twice until tungsten beads could be easily separated and resuspended. Before loading onto the biolistic transformation device, 60 μl was used to resuspend the beads and 10 μl was used per shot. Operation of the biolistic transformation device (Biorad, PDS-1000 He™ system) was according to the manufacturer’s protocol, except that the stopping screen was removed. 1100 psi disks and the highest position of the receiving petri plate were used in all bombardments to generate strains in this paper. A reading of 27.5 mmHg was reached on the vaccum gauge and the average pressure of breaking is 1000 psi. These conditions might require optimization based on the individual machine. Colonies on the biolistic transformation receiving plate (SD-Leu-sorbitol) were replica plated onto a YPD plate spread with strain NB40-3C with missing sections of COX2. The YPD plate was incubated at 30C overnight to allow mating. The next day, the YPD plate was replica plated onto YPEG plates to select for strains with recombined mtDNA capable of respiration. Typically, 3-10 respiratory competent colonies were isolated per plate in this step. Successful insertion of the *ARG8^m^** was confirmed by repeated streaking and colony PCR (Forward primer: 5’-CACAAGAGGTAAAAATGCTAAATTATATGATGATGTAAATGGTAAAG-3’, reverse primer: 5’-AGCATATACAGCTTCGATAGCTTTTTCGAAAGC-3’) from a yeast crude lysate (boiled at 95 °C in 20 mM NaOH). sgRNA expression was confirmed by RT-qPCR.

Diploids were selected by streaking confirmed, stable strains on SD-Ura-Ade. To sporulate the diploid, 200 µl of an overnight saturated YPD culture of the diploid was washed in 5 ml sterile water and resuspended in 2.5 ml SPM medium (3 g/L potassium acetate and 0.2 g/L raffinose in sterile water). The SPM culture was incubated at 23 °C with good aeration for 2-4 days until tetrads could be readily observed under the microscope. The tetrads were then dissected on YPD plates to give haploid strains.

**MitoTracker imaging.** Related to Figure S2B, S8A, B, E, F. An overnight saturated culture in SD-Leu media (0.77 g/L CSM-Leu, 6.7 g/L yeast nitrogen base, 20 g/L glucose) was diluted and grown for 4 hours to OD600 = 0.5. MitoTracker Red CMX Ros (Invitrogen) was added to a final concentration of 100 nM and incubated for 1 hr at 30 °C before imaging.

Imaging was performed with a Zeiss Axio Observer Z1 microscope on a Yokogawa spinning disk confocal head, Cascade IIL:512 camera, 50 mW 491 nm and 561 nm lasers. EGFP images were captured through the 488 nm channel (491 nm laser excitation, 528/38 nm emission) and MitoTracker images were captured through the 568 nm channel (561 nm laser excitation, 617/73 nm emission) through a 63× oil-immersion objective. Exposure times were 100 to 200 ms per channel. Image acquisition and processing was carried out with the SlideBook 5.0 software (Intelligent Imaging Innovations, 3i).

**Mitoplast preparation.** Related to Figure S4A and C. Following mitochondria purification, the OMM was lysed through adding 0.01-0.05% digitonin solution in mitochondria purification buffer for 10 min at 4 °C with occasional shaking. After 10 min, digitonin was diluted by adding 10x volume of mitochondria purification buffer without digitonin. Mitoplasts were pelleted with 12,000 g for 15 min at 4 °C on a table top centrifuge and subjected to the same RNAse treatment as purified mitochondria.

**Protein and plasmid sequences**

Protein sequence:

ABE^m^-linker-ARG8^m^

MSEVEFSHEYWMRHALTLAKRARDEREVPVGAVLVLNNRVIGEGWNRAIGLHDPTAHAEIMALRQGGLVMQNYRLIDATLYVTFEPCVMCAGAMIHSRIGRVVFGVRNSKRGAAGSLMNVLNYPGMNHRVEITEGILADECAALLCDFYRMPRQVFNAQKKAQSSINSGGSSGGSSGSETPGTSESATPESSGGSSGGSDKKYSIGLAIGTNSVGWAVITDEYKVPSKKFKVLGNTDRHSIKKNLIGALLFDSGETAEATRLKRTARRRYTRRKNRICYLQEIFSNEMAKVDDSFFHRLEESFLVEEDKKHERHPIFGNIVDEVAYHEKYPTIYHLRKKLVDSTDKADLRLIYLALAHMIKFRGHFLIEGDLNPDNSDVDKLFIQLVQTYNQLFEENPIDASGVDAKAILSARLSKSRRLENLIAQLPGEKKNGLFGNLIALSLGLTPNFKSNFDLAEDAKLQLSKDTYDDDLDNLLAQIGDQYADLFLAAKNLSDAILLSDILRVNTEITKAPLSASMIKRYDEHHQDLTLLKALVRQQLPEKYKEIFFDQSKNGYAGYIDGGASQEEFYKFIKPILEKMDGTEELLVKLNREDLLRKQRTFDNGSIPHQIHLGELHAILRRQEDFYPFLKDNREKIEKILTFRIPYYVGPLARGNSRFAWMTRKSEETITPWNFEEVVDKGASAQSFIERMTNFDKNLPNEKVLPKHSLLYEYFTVYNELTKVKYVTEGMRKPAFLSGEQKKAIVDLLFKTNRKVTVKQLKEDYFKKIECFDSVEISGVEDRFNASLGTYHDLLKIIKDKDFLDNEENEDILEDIVLTLTLFEDREMIEERLKTYAHLFDDKVMKQLKRRRYTGWGRLSRKLINGIRDKQSGKTILDFLKSDGFANRNFMQLIHDDSLTFKEDIQKAQVSGQGDSLHEHIANLAGSPAIKKGILQTVKVVDELVKVMGRHKPENIVIEMARENQTTQKGQKNSRERMKRIEEGIKELGSQILKEHPVENTQLQNEKLYLYYLQNGRDMYVDQELDINRLSDYDVDHIVPQSFLKDDSIDNKVLTRSDKNRGKSDNVPSEEVVKKMKNYWRQLLNAKLITQRKFDNLTKAERGGLSELDKAGFIKRQLVETRQITKHVAQILDSRMNTKYDENDKLIREVKVITLKSKLVSDFRKDFQFYKVREINNYHHAHDAYLNAVVGTALIKKYPKLESEFVYGDYKVYDVRKMIAKSEQEIGKATAKYFFYSNIMNFFKTEITLANGEIRKRPLIETNGETGEIVWDKGRDFATVRKVLSMPQVNIVKKTEVQTGGFSKESILPKRNSDKLIARKKDWDPKKYGGFDSPTVAYSVLVVAKVEKGKSKKLKSVKELLGITIMERSSFEKNPIDFLEAKGYKEVKKDLIIKLPKYSLFELENGRKRMLASAGELQKGNELALPSKYVNFLYLASHYEKLKGSPEDNEQKQLFVEQHKHYLDEIIEQISEFSKRVILADANLDKVLSAYNKHRDKPIREQAENIIHLFTLTNLGAPAAFKYFDTTIDRKRYTSTKEVLDATLIHQSITGLYETRIDLSQLGGDSGGSFKRYLSSTSSRRFTSILEEKAFQVTTYSRPEDLCITRGKNAKLYDDVNGKEYIDFTAGIAVTALGHANPKVAEILHHQANKLVHSSNLYFTKECLDLSEKIVEKTKQFGGQHDASRVFLCNSGTEANEAALKFAKKHGIMKNPSKQGIVAFENSFHGRTMGALSVTWNSKYRTPFGDLVPHVSFLNLNDEMTKLQSYIETKKDEIAGLIVEPIQGEGGVFPVEVEKLTGLKKICQDNDVIVIHDEIQCGLGRSGKLWAHAYLPSEAHPDIFTSAKALGNGFPIAATIVNEKVNNALRVGDHGTTYGGNPLACSVSNYVLDTIADEAFLKQVSKKSDILQKRLREIQAKYPNQIKTIRGKGLMLGAEFVEPPTEVIKKARELGLLIITAGKSTVRFVPALTIEDELIEEGMDAFEKAIEAVYAGKPIPNPLLGLDST

ABE^m^: Red, ARG8^m^: Blue

tMsk1

MSLAHAVDTSKMEATRRNGQIVKDLGRYYPSMSESALHDLCQEYKEVTIADFNERFLGNPATLHHEDNPNLLLSINGRIKSIRFSGQKIVFIDLYNGSSGLKNDTQLQLIVNYNKIGGSSEDKANFSEYMNFLKKGDYIKALGYPGFSQSRVKMLSLICNKLPIVLSVSQLPLPSRLNDETKIKSNRVVDYQLNGTQTLLVRARIIKLLRKFLDDRNFVEVETPILSSKSNGAMAKPFITSSKDFDHLELRIAPELWLKRLIISGLQKVYEIGKVFRNEGIDSTHNAEFSTLEFYETYMSMDDIVTRTEDLFKFLITNLQKFFQDTRLPVPKTFSELHLALSENNWKFRKVEFLPTLNKELGIDLMNSGLDINKPSELLKALPKDIAKKYFPSADNTGQLSSLQILNKLSDVFLEQRHCQSTLPTVIYHQPAILSPLAKTDPQNKQVTKRFEVFIKGKEYINAYEEENCPQLQLQKFLQQKQINELTGNKTETLSPVIDYQYVETMKYGMPPVGGFGLGIDRLCMLFCDKKRIEEVLPFGCVDDVNRQKGKPIPNPLLGLDSTHHHHHH*

tMsk1: Red, 6xHis tag: Blue

MTS-GFP-lambda

MLSLRQSIRFFKPATRTLCSSRPLVATVSKGEELFTGVVPILVELDGDVNGHKFSVSGEGEGDATYGKLTLKFICTTGKLPVPWPTLVTTLTYGVQCFSRYPDHMKQHDFFKSAMPEGYVQERTIFFKDDGNYKTRAEVKFEGDTLVNRIELKGIDFKEDGNILGHKLEYNYNSHNVYIMADKQKNGIKVNFKIRHNIEDGSVQLADHYQQNTPIGDGPVLLPDNHYLSTQSALSKDPNEKRDHMVLLEFVTAAGITLGMDELYKSAPPLDGAGAGAGAGAGAGGLATMDAQTRRRERRAEKQAQWKAANLEPPLDGAGAGAGAGAGAGGLATPPSA

COX4 MTS: Red; GFP: green; Lambda peptide: blue.

MTS-GFPN-lambda-GFPC

MLSLRQSIRFFKPATRTLCSSRPLVATVSKGEELFTGVVPILVELDGDVNGHKFSVSGEGEGDATYGKLTLKFICTTGKLPVPWPTLVTTLTYGVQCFSRYPDHMKQHDFFKSAMPEGYVQERTIFFKDDGNYKTRAEVKFEGDTLVNRIELKGIDFKEDGNILGHKLEYNSAPPLDGAGAGAGAGAGAGGLATMDAQTRRRERRAEKQAQWKAANLEPPLDGAGAGAGAGAGAGGLATPPSANSHNVYIMADKQKNGIKVNFKIRHNIEDGSVQLADHYQQNTPIGDGPVLLPDNHYLSTQSALSKDPNEKRDHMVLLEFVTAAGITLGMDELYK

COX4 MTS: Red; GFP: green; Lambda peptide: blue.

DNA sequences:

For biolistic transformation

COX2::ARG8m*-HH-sgRNA(Arg8m)-HDV-COX2

Blue: Upstream sequence of COX2; Green: COX2 5’UTR; Orange:ARG8^m^ (with STOP at Q79); Purple: V5; Light green: COX2 3’UTR; Yellow highlight: HH; Teal highlight: sgRNA(targeting ARG8^m^) Green highlight: HDV; Light blue: COX2

AAGCTTATCGGCGGGGACCCCGAAGGAGTATAAATAAAAATTAATAATATATTATATATATATTATATTAATAATAATAATAATAATAATAAATAATAACTCCTTGCTTCATACCTTTATAAATAAGGTAATCACTAATATATTATAATAATAATAATTATATTTATAATTTATATATTTATATATATAAAAATTATATATATTATATATAATCTAAATATTATATATTTTAATAAATATTAATATATATGATATGAATATTATTAGTTTTCGGGAAGCGGGAATTCATAAATTTTAATTAAAAGTAGTATTAACATATTATAAATAGACAAAAGAGTCTAAAGGTTAAGATTTATTCATatgttcaaaagatatttatcatcaacatcatcaagaagattcacatcaatcttagaagaaaaagcttttCAAgtaacaacatattcaagacctgaagatttatgtatcacaagaggtaaaaatgctaaattatatgatgatgtaaatggtaaagaatatatcgatttcacagctggtattgctgtaacagctttaggtcatgctaatcctaaagtagcCgaaatcttacatcaCTAAgctaataaattagtCcattcatcaaatttatatttcacaaaagaatgtttagatttatcagaaaaaatcgtagaaaaaacaaaaCAAttcggtggtCAAcatgatgcttcaagagtattcttatgtaattcaggtaccgaagctaatgaagctgcattaaaattcgctaaaaaacatggtatcatgaaaaatcctagtaaaCAAggtattgtagctttcgaaaattcattccatggtagaacaatgggtgctttatcagtaacaTGAaattcaaaatatagAacacctttcggtgatttagtacctcatgtatcattcttaaatttaaatgatgaaatgacaaaattaCAAtcatatatcgaaacaaaaaaagatgaaatcgctggtttaatcgtagaacctattCAAggtgaaggtggtgtatttcctgtagaagtagaaaaattaactggtttaaaaaaaatctgtCAAgataatgatgtaatcgtaattcatgatgaaattCAAtgtggtttaggtagatcaggtaaattaTGAgctcatgcttatttaccttcagaagctcatcctgatattttcacatcagctaaagcattaggtaatggtttccctattgctgctacaatcgtaaatgaaaaagtaaataatgctttaagagtaggtgatcatggtacaacatatggtggtaatcctttagcttgttcagtatcaaattatgtattagatacaattgctgatgaagcattcttaaaaCAAgtatcaaaaaaatcagatatcttaCAAaaaagattaagagaaatcCAAgctaaatatcctaatCAAatcaaaacaatcagaggtaaaggtttaatgttaggtgctgaatttgtagaacctccaacagaagtaatcaaaaaagctagagaattaggtttattaatcatcacagctggtaaatcaacagtaagattcgtacctgctttaacaatcgaagatgaattaatcgaagaaggtatggatgctttcgaaaaagctatcgaagctgtatatgctGGCAAGCCCAtccccaaccccttgttaggcttagaCAGCACCtaaCTCGAGTTAATATTTACTTATTATTAATATTTTTAATTATTAAAAATAATAATAATAATAATAATTATAATAATATTCTTAAATATAATAAAGATATAGATTTATATTCTATTCAATCACCTTATGAATTCTAAgctCTGATGAGTCCGTGAGGACGAAACGAGTAAGCTCGTCagcTTAgtgatgtaagatttgttttagagctagaaatagcaagttaaaataaggctagtccgttatcaacttgaaaaagtggcaccgagtcggtggtgcttttGGCCGGCATGGTCCCAGCCTCCTCGCTGGCGCCGGCTGGGCAACATGCTTCGGCATGGCGAATGGGACGAATTCCGTAAGGAGTGAGGGACCCCTCCCTATACTAACGGGAGGGGGACCGAACCCCGAAGGAGTTTTATTTTTAGTATTTTATAAAATATATATTTATATGATTAATAATATTATATATATTATTTATAAAAATAATATATAATTTTAATTATTTTTAATAAAAAAAGGTGGGGTTTGGTAATATAATATTTTTATTTTATTTATAATATATAATAATAAATTATAAATAAATTTTAATTAAAAGTAGTATTAACATATTATAAATAGACAAAAGAGTCTAAAGGTTAAGATTTATTAAAATGTTAGATTTATTAAGATTACAATTAACAACATTCATTATGAATGATGTACCAACACCTTATGCATGTTATTTTCAGGATTCAGCAACACCAAATCAAGAAGGTATTTTAGAATTACATGATAATATTATGTTTTATTTATTAGTTATTTTAGGTTTAGTATCTTGAATGTTATATACAATTGTTATAACATATTCAAAAAATCCTATTGCATATAAATATATTAAACATGGACAAACTATTGAAGTTATTTGAACAATTTTTCCAGCTGTAATTTTATTAATTATTGCTTTCCCTTCATTTATTTTATTATATTTATGTGATGAAGTTATTTCACCAGCTATAACTATTAAAGCTATTGGATATCAATGATATTGAAAATATGAATATTCAGATTTTATTAATGATAGTGGTGAAACTGTTGAATTTGAATCATATGTTATTCCTGATGAATTATTAGAAGAAGGACAATTAAGATTATTAGATACTGATACTTCTATAGTTGTACCTGTAGATACACATATTAGATTCGTTGTAACAGCTGCTGATGTTATTCATGATTTTGCTATCCCAAGTTTAGGTATTAAAGTTGATGCTACTCCTGGTAGATTAAATCAAGTTTCTGCTTTAATTCAAAGAGAAGGTGTCTTCTATGGGGCATGTTCTGAGTTGTGTGGGACAGGTCATGCAAATATGCCAATTAAGATCGAAGCAGTATCATTACCTAAATTTTTGGAATGATTAAATGAACAATAATTAATATTTACTTATTATTAATATTTTTAATTATTAAAAATAATAATAATAATAATAATTATAATAATATTCTTAAATATAATAAAGATATAGATTTATATTCTATTCAATCACCTTATATTAAAAATATAAATATTATTAAAAGAGGTTATCATACTTCTTTAAATAATAAATTAATTATTGGTTCAAAAGGATAATAAAAATAATAATAAGAATAACCCAGAATTGATAATTTTTATAAATGATTAGTGGATTTACAGATGGAGATGGTAGTTTTTATATTAAATTAAATGATAAAAAATATTTAAGATTTTTTTATGGTTTTAGAATACATATTGATGATAAAGCATGTTTAGAAAAGATTAGAAATATATTAAATATACCTTCTAATTTTGAAGAACTACTTAAAACAATTATATTAGTAAATTCACAAAAGAAATGGTTATATTCTAATATTGTAACTATTTTTGATAAGTATCCTTGTTTAACAATTAAATATTATAGTTATTATAAATGAAAAATAGCTATAATTAATAATTTAAATGGTATATCTTATAATAATAAAGATTTATTAAATATTAAAAATACAATTAATAATTATGAAGTTATACCTAATTTAAAAATTCCATATGATAAAATAAATGATTATTGAATTTTAGGTTTTATTGAAGCTGAAGGTTCATTTGATCTATCTCCAAAACGTAATATTTGTGGTTTTAATGTTTCACAACATAAACGTAGTATTAATACATTAAAAGCTATTAAATCTTATGTATTAAATAATTGAAAACCAATTGATAATACACCATTATTAATTAAAAATAAATTATTAAAAGATTGAGATTCATCTATTAAATTAACTAAACCTGATAAAAATGGAGTTATTAAATTAGAATTTAATAGAATAGATTTTTTATATTATGTTATTTTACCTAAATTATATTCATTAAAATGATATAGTCGTAAAGAAATTGATTTCCAATTATGAAAAACACTTATAGAAATCTATATAAAAGGTTTACATAATACACTTAAAGGTTCTAATTTATTAAAATTAATTAATAATAATATTAATAAAAAAAGATATTATTCTAATTATAATATTTCTCCTTTCGGGGTTCCGGCTCCCGTGGCCGGGCCCCGATGATAAGCTGTCAAACATGAGAATTAATTC

COX2::ARG8m*(Q79*)-COX2

Blue: Upstream sequence of COX2; Green: COX2 5’UTR; Orange:ARG8^m^ (with STOP at Q79); Purple: V5; Light green: COX2 3’UTR; Light blue: COX2

AAGCTTATCGGCGGGGACCCCGAAGGAGTATAAATAAAAATTAATAATATATTATATATATATTATATTAATAATAATAATAATAATAATAAATAATAACTCCTTGCTTCATACCTTTATAAATAAGGTAATCACTAATATATTATAATAATAATAATTATATTTATAATTTATATATTTATATATATAAAAATTATATATATTATATATAATCTAAATATTATATATTTTAATAAATATTAATATATATGATATGAATATTATTAGTTTTCGGGAAGCGGGAATTCATAAATTTTAATTAAAAGTAGTATTAACATATTATAAATAGACAAAAGAGTCTAAAGGTTAAGATTTATTCATatgttcaaaagatatttatcatcaacatcatcaagaagattcacatcaatcttagaagaaaaagcttttCAAgtaacaacatattcaagacctgaagatttatgtatcacaagaggtaaaaatgctaaattatatgatgatgtaaatggtaaagaatatatcgatttcacagctggtattgctgtaacagctttaggtcatgctaatcctaaagtagcCgaaatcttacatcaCTAAgctaataaattagtCcattcatcaaatttatatttcacaaaagaatgtttagatttatcagaaaaaatcgtagaaaaaacaaaaCAAttcggtggtCAAcatgatgcttcaagagtattcttatgtaattcaggtaccgaagctaatgaagctgcattaaaattcgctaaaaaacatggtatcatgaaaaatcctagtaaaCAAggtattgtagctttcgaaaattcattccatggtagaacaatgggtgctttatcagtaacaTGAaattcaaaatatagAacacctttcggtgatttagtacctcatgtatcattcttaaatttaaatgatgaaatgacaaaattaCAAtcatatatcgaaacaaaaaaagatgaaatcgctggtttaatcgtagaacctattCAAggtgaaggtggtgtatttcctgtagaagtagaaaaattaactggtttaaaaaaaatctgtCAAgataatgatgtaatcgtaattcatgatgaaattCAAtgtggtttaggtagatcaggtaaattaTGAgctcatgcttatttaccttcagaagctcatcctgatattttcacatcagctaaagcattaggtaatggtttccctattgctgctacaatcgtaaatgaaaaagtaaataatgctttaagagtaggtgatcatggtacaacatatggtggtaatcctttagcttgttcagtatcaaattatgtattagatacaattgctgatgaagcattcttaaaaCAAgtatcaaaaaaatcagatatcttaCAAaaaagattaagagaaatcCAAgctaaatatcctaatCAAatcaaaacaatcagaggtaaaggtttaatgttaggtgctgaatttgtagaacctccaacagaagtaatcaaaaaagctagagaattaggtttattaatcatcacagctggtaaatcaacagtaagattcgtacctgctttaacaatcgaagatgaattaatcgaagaaggtatggatgctttcgaaaaagctatcgaagctgtatatgctGGCAAGCCCAtccccaaccccttgttaggcttagaCAGCACCtaaCTCGAGTTAATATTTACTTATTATTAATATTTTTAATTATTAAAAATAATAATAATAATAATAATTATAATAATATTCTTAAATATAATAAAGATATAGATTTATATTCTATTCAATCACCTTATGAATTCGAATTCCGTAAGGAGTGAGGGACCCCTCCCTATACTAACGGGAGGGGGACCGAACCCCGAAGGAGTTTTATTTTTAGTATTTTATAAAATATATATTTATATGATTAATAATATTATATATATTATTTATAAAAATAATATATAATTTTAATTATTTTTAATAAAAAAAGGTGGGGTTTGGTAATATAATATTTTTATTTTATTTATAATATATAATAATAAATTATAAATAAATTTTAATTAAAAGTAGTATTAACATATTATAAATAGACAAAAGAGTCTAAAGGTTAAGATTTATTAAAATGTTAGATTTATTAAGATTACAATTAACAACATTCATTATGAATGATGTACCAACACCTTATGCATGTTATTTTCAGGATTCAGCAACACCAAATCAAGAAGGTATTTTAGAATTACATGATAATATTATGTTTTATTTATTAGTTATTTTAGGTTTAGTATCTTGAATGTTATATACAATTGTTATAACATATTCAAAAAATCCTATTGCATATAAATATATTAAACATGGACAAACTATTGAAGTTATTTGAACAATTTTTCCAGCTGTAATTTTATTAATTATTGCTTTCCCTTCATTTATTTTATTATATTTATGTGATGAAGTTATTTCACCAGCTATAACTATTAAAGCTATTGGATATCAATGATATTGAAAATATGAATATTCAGATTTTATTAATGATAGTGGTGAAACTGTTGAATTTGAATCATATGTTATTCCTGATGAATTATTAGAAGAAGGACAATTAAGATTATTAGATACTGATACTTCTATAGTTGTACCTGTAGATACACATATTAGATTCGTTGTAACAGCTGCTGATGTTATTCATGATTTTGCTATCCCAAGTTTAGGTATTAAAGTTGATGCTACTCCTGGTAGATTAAATCAAGTTTCTGCTTTAATTCAAAGAGAAGGTGTCTTCTATGGGGCATGTTCTGAGTTGTGTGGGACAGGTCATGCAAATATGCCAATTAAGATCGAAGCAGTATCATTACCTAAATTTTTGGAATGATTAAATGAACAATAATTAATATTTACTTATTATTAATATTTTTAATTATTAAAAATAATAATAATAATAATAATTATAATAATATTCTTAAATATAATAAAGATATAGATTTATATTCTATTCAATCACCTTATATTAAAAATATAAATATTATTAAAAGAGGTTATCATACTTCTTTAAATAATAAATTAATTATTGGTTCAAAAGGATAATAAAAATAATAATAAGAATAACCCAGAATTGATAATTTTTATAAATGATTAGTGGATTTACAGATGGAGATGGTAGTTTTTATATTAAATTAAATGATAAAAAATATTTAAGATTTTTTTATGGTTTTAGAATACATATTGATGATAAAGCATGTTTAGAAAAGATTAGAAATATATTAAATATACCTTCTAATTTTGAAGAACTACTTAAAACAATTATATTAGTAAATTCACAAAAGAAATGGTTATATTCTAATATTGTAACTATTTTTGATAAGTATCCTTGTTTAACAATTAAATATTATAGTTATTATAAATGAAAAATAGCTATAATTAATAATTTAAATGGTATATCTTATAATAATAAAGATTTATTAAATATTAAAAATACAATTAATAATTATGAAGTTATACCTAATTTAAAAATTCCATATGATAAAATAAATGATTATTGAATTTTAGGTTTTATTGAAGCTGAAGGTTCATTTGATCTATCTCCAAAACGTAATATTTGTGGTTTTAATGTTTCACAACATAAACGTAGTATTAATACATTAAAAGCTATTAAATCTTATGTATTAAATAATTGAAAACCAATTGATAATACACCATTATTAATTAAAAATAAATTATTAAAAGATTGAGATTCATCTATTAAATTAACTAAACCTGATAAAAATGGAGTTATTAAATTAGAATTTAATAGAATAGATTTTTTATATTATGTTATTTTACCTAAATTATATTCATTAAAATGATATAGTCGTAAAGAAATTGATTTCCAATTATGAAAAACACTTATAGAAATCTATATAAAAGGTTTACATAATACACTTAAAGGTTCTAATTTATTAAAATTAATTAATAATAATATTAATAAAAAAAGATATTATTCTAATTATAATATTTCTCCTTTCGGGGTTCCGGCTCCCGTGGCCGGGCCCCGATGATAAGCTGTCAAACATGAGAATTAATTC

COX2::ARG8m*(W168*)-HH-sgRNA-HDV-COX2

Blue: Upstream sequence of COX2; Green: COX2 5’UTR; Orange:ARG8^m^ (with STOP at W168*); Purple: V5; Yellow highlight: HH; Teal highlight: sgRNA(targeting ARG8^m*^); Green highlight: HDV; Light green: COX2 3’UTR; Light blue: COX2

AAGCTTATCGGCGGGGACCCCGAAGGAGTATAAATAAAAATTAATAATATATTATATATATATTATATTAATAATAATAATAATAATAATAAATAATAACTCCTTGCTTCATACCTTTATAAATAAGGTAATCACTAATATATTATAATAATAATAATTATATTTATAATTTATATATTTATATATATAAAAATTATATATATTATATATAATCTAAATATTATATATTTTAATAAATATTAATATATATGATATGAATATTATTAGTTTTCGGGAAGCGGGAATTCATAAATTTTAATTAAAAGTAGTATTAACATATTATAAATAGACAAAAGAGTCTAAAGGTTAAGATTTATTCATatgttcaaaagatatttatcatcaacatcatcaagaagattcacatcaatcttagaagaaaaagcttttCAAgtaacaacatattcaagacctgaagatttatgtatcacaagaggtaaaaatgctaaattatatgatgatgtaaatggtaaagaatatatcgatttcacagctggtattgctgtaacagctttaggtcatgctaatcctaaagtagctgaaatcttacatcatCAAgctaataaattagtacattcatcaaatttatatttcacaaaagaatgtttagatttatcagaaaaaatcgtagaaaaaacaaaaCAAttcggtggtCAAcatgatgcttcaagagtattcttatgtaattcaggtaccgaagctaatgaagctgcattaaaattcgctaaaaaacatggtatcatgaaaaatcctagtaaaCAAggtattgtagctttcgaaaattcattccatggtagaacaatgggtgctttatcagtaacaTAAaattcaaaatataggacacctttcggtgatttagtacctcatgtatcattcttaaatttaaatgatgaaatgacaaaattaCAAtcatatatcgaaacaaaaaaagatgaaatcgctggtttaatcgtagaacctattCAAggtgaaggtggtgtatttcctgtagaagtagaaaaattaactggtttaaaaaaaatctgtCAAgataatgatgtaatcgtaattcatgatgaaattCAAtgtggtttaggtagatcaggtaaattaTGAgctcatgcttatttaccttcagaagctcatcctgatattttcacatcagctaaagcattaggtaatggtttccctattgctgctacaatcgtaaatgaaaaagtaaataatgctttaagagtaggtgatcatggtacaacatatggtggtaatcctttagcttgttcagtatcaaattatgtattagatacaattgctgatgaagcattcttaaaaCAAgtatcaaaaaaatcagatatcttaCAAaaaagattaagagaaatcCAAgctaaatatcctaatCAAatcaaaacaatcagaggtaaaggtttaatgttaggtgctgaatttgtagaacctccaacagaagtaatcaaaaaagctagagaattaggtttattaatcatcacagctggtaaatcaacagtaagattcgtacctgctttaacaatcgaagatgaattaatcgaagaaggtatggatgctttcgaaaaagctatcgaagctgtatatgcttaaGGCAAGCCCAtccccaaccccttgttaggcttagaCAGCACCtaaCTCGAGTTAATATTTACTTATTATTAATATTTTTAATTATTAAAAATAATAATAATAATAATAATTATAATAATATTCTTAAATATAATAAAGATATAGATTTATATTCTATTCAATCACCTTATGAATTCTAAcatCTGATGAGTCCGTGAGGACGAAACGAGTAAGCTCGTCtaacaTAAaattcaaaatatgttttagagctagaaatagcaagttaaaataaggctagtccgttatcaacttgaaaaagtggcaccgagtcggtggtgcttttGGCCGGCATGGTCCCAGCCTCCTCGCTGGCGCCGGCTGGGCAACATGCTTCGGCATGGCGAATGGGACGAATTCCGTAAGGAGTGAGGGACCCCTCCCTATACTAACGGGAGGGGGACCGAACCCCGAAGGAGTTTTATTTTTAGTATTTTATAAAATATATATTTATATGATTAATAATATTATATATATTATTTATAAAAATAATATATAATTTTAATTATTTTTAATAAAAAAAGGTGGGGTTTGGTAATATAATATTTTTATTTTATTTATAATATATAATAATAAATTATAAATAAATTTTAATTAAAAGTAGTATTAACATATTATAAATAGACAAAAGAGTCTAAAGGTTAAGATTTATTAAAATGTTAGATTTATTAAGATTACAATTAACAACATTCATTATGAATGATGTACCAACACCTTATGCATGTTATTTTCAGGATTCAGCAACACCAAATCAAGAAGGTATTTTAGAATTACATGATAATATTATGTTTTATTTATTAGTTATTTTAGGTTTAGTATCTTGAATGTTATATACAATTGTTATAACATATTCAAAAAATCCTATTGCATATAAATATATTAAACATGGACAAACTATTGAAGTTATTTGAACAATTTTTCCAGCTGTAATTTTATTAATTATTGCTTTCCCTTCATTTATTTTATTATATTTATGTGATGAAGTTATTTCACCAGCTATAACTATTAAAGCTATTGGATATCAATGATATTGAAAATATGAATATTCAGATTTTATTAATGATAGTGGTGAAACTGTTGAATTTGAATCATATGTTATTCCTGATGAATTATTAGAAGAAGGACAATTAAGATTATTAGATACTGATACTTCTATAGTTGTACCTGTAGATACACATATTAGATTCGTTGTAACAGCTGCTGATGTTATTCATGATTTTGCTATCCCAAGTTTAGGTATTAAAGTTGATGCTACTCCTGGTAGATTAAATCAAGTTTCTGCTTTAATTCAAAGAGAAGGTGTCTTCTATGGGGCATGTTCTGAGTTGTGTGGGACAGGTCATGCAAATATGCCAATTAAGATCGAAGCAGTATCATTACCTAAATTTTTGGAATGATTAAATGAACAATAATTAATATTTACTTATTATTAATATTTTTAATTATTAAAAATAATAATAATAATAATAATTATAATAATATTCTTAAATATAATAAAGATATAGATTTATATTCTATTCAATCACCTTATATTAAAAATATAAATATTATTAAAAGAGGTTATCATACTTCTTTAAATAATAAATTAATTATTGGTTCAAAAGGATAATAAAAATAATAATAAGAATAACCCAGAATTGATAATTTTTATAAATGATTAGTGGATTTACAGATGGAGATGGTAGTTTTTATATTAAATTAAATGATAAAAAATATTTAAGATTTTTTTATGGTTTTAGAATACATATTGATGATAAAGCATGTTTAGAAAAGATTAGAAATATATTAAATATACCTTCTAATTTTGAAGAACTACTTAAAACAATTATATTAGTAAATTCACAAAAGAAATGGTTATATTCTAATATTGTAACTATTTTTGATAAGTATCCTTGTTTAACAATTAAATATTATAGTTATTATAAATGAAAAATAGCTATAATTAATAATTTAAATGGTATATCTTATAATAATAAAGATTTATTAAATATTAAAAATACAATTAATAATTATGAAGTTATACCTAATTTAAAAATTCCATATGATAAAATAAATGATTATTGAATTTTAGGTTTTATTGAAGCTGAAGGTTCATTTGATCTATCTCCAAAACGTAATATTTGTGGTTTTAATGTTTCACAACATAAACGTAGTATTAATACATTAAAAGCTATTAAATCTTATGTATTAAATAATTGAAAACCAATTGATAATACACCATTATTAATTAAAAATAAATTATTAAAAGATTGAGATTCATCTATTAAATTAACTAAACCTGATAAAAATGGAGTTATTAAATTAGAATTTAATAGAATAGATTTTTTATATTATGTTATTTTACCTAAATTATATTCATTAAAATGATATAGTCGTAAAGAAATTGATTTCCAATTATGAAAAACACTTATAGAAATCTATATAAAAGGTTTACATAATACACTTAAAGGTTCTAATTTATTAAAATTAATTAATAATAATATTAATAAAAAAAGATATTATTCTAATTATAATATTTCTCCTTTCGGGGTTCCGGCTCCCGTGGCCGGGCCCCGATGATAAGCTGTCAAACATGAGAATTAATTC

COX2::ARG8m*-HH-sgRNA(Arg8^m^, Cas12)-HDV-COX2

Blue: Upstream sequence of COX2; Green: COX2 5’UTR; Orange:ARG8^m^ (with STOP at Q79); Purple: V5; Light green: COX2 3’UTR; Yellow highlight: HH; Teal highlight: sgRNA(targeting ARG8^m^, LbCpf) Green highlight: HDV; Light blue: COX2

AAGCTTATCGGCGGGGACCCCGAAGGAGTATAAATAAAAATTAATAATATATTATATATATATTATATTAATAATAATAATAATAATAATAAATAATAACTCCTTGCTTCATACCTTTATAAATAAGGTAATCACTAATATATTATAATAATAATAATTATATTTATAATTTATATATTTATATATATAAAAATTATATATATTATATATAATCTAAATATTATATATTTTAATAAATATTAATATATATGATATGAATATTATTAGTTTTCGGGAAGCGGGAATTCATAAATTTTAATTAAAAGTAGTATTAACATATTATAAATAGACAAAAGAGTCTAAAGGTTAAGATTTATTCATatgttcaaaagatatttatcatcaacatcatcaagaagattcacatcaatcttagaagaaaaagcttttCAAgtaacaacatattcaagacctgaagatttatgtatcacaagaggtaaaaatgctaaattatatgatgatgtaaatggtaaagaatatatcgatttcacagctggtattgctgtaacagctttaggtcatgctaatcctaaagtagcCgaaatcttacatcaCTAAgctaataaattagtCcattcatcaaatttatatttcacaaaagaatgtttagatttatcagaaaaaatcgtagaaaaaacaaaaCAAttcggtggtCAAcatgatgcttcaagagtattcttatgtaattcaggtaccgaagctaatgaagctgcattaaaattcgctaaaaaacatggtatcatgaaaaatcctagtaaaCAAggtattgtagctttcgaaaattcattccatggtagaacaatgggtgctttatcagtaacaTGAaattcaaaatatagAacacctttcggtgatttagtacctcatgtatcattcttaaatttaaatgatgaaatgacaaaattaCAAtcatatatcgaaacaaaaaaagatgaaatcgctggtttaatcgtagaacctattCAAggtgaaggtggtgtatttcctgtagaagtagaaaaattaactggtttaaaaaaaatctgtCAAgataatgatgtaatcgtaattcatgatgaaattCAAtgtggtttaggtagatcaggtaaattaTGAgctcatgcttatttaccttcagaagctcatcctgatattttcacatcagctaaagcattaggtaatggtttccctattgctgctacaatcgtaaatgaaaaagtaaataatgctttaagagtaggtgatcatggtacaacatatggtggtaatcctttagcttgttcagtatcaaattatgtattagatacaattgctgatgaagcattcttaaaaCAAgtatcaaaaaaatcagatatcttaCAAaaaagattaagagaaatcCAAgctaaatatcctaatCAAatcaaaacaatcagaggtaaaggtttaatgttaggtgctgaatttgtagaacctccaacagaagtaatcaaaaaagctagagaattaggtttattaatcatcacagctggtaaatcaacagtaagattcgtacctgctttaacaatcgaagatgaattaatcgaagaaggtatggatgctttcgaaaaagctatcgaagctgtatatgctGGCAAGCCCAtccccaaccccttgttaggcttagaCAGCACCtaaCTCGAGTTAATATTTACTTATTATTAATATTTTTAATTATTAAAAATAATAATAATAATAATAATTATAATAATATTCTTAAATATAATAAAGATATAGATTTATATTCTATTCAATCACCTTATGAATTCGAAATTCTGATGAGTCCGTGAGGACGAAACGAGTAAGCTCGTCAATTTCTACTGTTGTAGATttagcTTAgtgatgtaagatttcGGCCGGCATGGTCCCAGCCTCCTCGCTGGCGCCGGCTGGGCAACATGCTTCGGCATGGCGAATGGGACGAATTCCGTAAGGAGTGAGGGACCCCTCCCTATACTAACGGGAGGGGGACCGAACCCCGAAGGAGTTTTATTTTTAGTATTTTATAAAATATATATTTATATGATTAATAATATTATATATATTATTTATAAAAATAATATATAATTTTAATTATTTTTAATAAAAAAAGGTGGGGTTTGGTAATATAATATTTTTATTTTATTTATAATATATAATAATAAATTATAAATAAATTTTAATTAAAAGTAGTATTAACATATTATAAATAGACAAAAGAGTCTAAAGGTTAAGATTTATTAAAATGTTAGATTTATTAAGATTACAATTAACAACATTCATTATGAATGATGTACCAACACCTTATGCATGTTATTTTCAGGATTCAGCAACACCAAATCAAGAAGGTATTTTAGAATTACATGATAATATTATGTTTTATTTATTAGTTATTTTAGGTTTAGTATCTTGAATGTTATATACAATTGTTATAACATATTCAAAAAATCCTATTGCATATAAATATATTAAACATGGACAAACTATTGAAGTTATTTGAACAATTTTTCCAGCTGTAATTTTATTAATTATTGCTTTCCCTTCATTTATTTTATTATATTTATGTGATGAAGTTATTTCACCAGCTATAACTATTAAAGCTATTGGATATCAATGATATTGAAAATATGAATATTCAGATTTTATTAATGATAGTGGTGAAACTGTTGAATTTGAATCATATGTTATTCCTGATGAATTATTAGAAGAAGGACAATTAAGATTATTAGATACTGATACTTCTATAGTTGTACCTGTAGATACACATATTAGATTCGTTGTAACAGCTGCTGATGTTATTCATGATTTTGCTATCCCAAGTTTAGGTATTAAAGTTGATGCTACTCCTGGTAGATTAAATCAAGTTTCTGCTTTAATTCAAAGAGAAGGTGTCTTCTATGGGGCATGTTCTGAGTTGTGTGGGACAGGTCATGCAAATATGCCAATTAAGATCGAAGCAGTATCATTACCTAAATTTTTGGAATGATTAAATGAACAATAATTAATATTTACTTATTATTAATATTTTTAATTATTAAAAATAATAATAATAATAATAATTATAATAATATTCTTAAATATAATAAAGATATAGATTTATATTCTATTCAATCACCTTATATTAAAAATATAAATATTATTAAAAGAGGTTATCATACTTCTTTAAATAATAAATTAATTATTGGTTCAAAAGGATAATAAAAATAATAATAAGAATAACCCAGAATTGATAATTTTTATAAATGATTAGTGGATTTACAGATGGAGATGGTAGTTTTTATATTAAATTAAATGATAAAAAATATTTAAGATTTTTTTATGGTTTTAGAATACATATTGATGATAAAGCATGTTTAGAAAAGATTAGAAATATATTAAATATACCTTCTAATTTTGAAGAACTACTTAAAACAATTATATTAGTAAATTCACAAAAGAAATGGTTATATTCTAATATTGTAACTATTTTTGATAAGTATCCTTGTTTAACAATTAAATATTATAGTTATTATAAATGAAAAATAGCTATAATTAATAATTTAAATGGTATATCTTATAATAATAAAGATTTATTAAATATTAAAAATACAATTAATAATTATGAAGTTATACCTAATTTAAAAATTCCATATGATAAAATAAATGATTATTGAATTTTAGGTTTTATTGAAGCTGAAGGTTCATTTGATCTATCTCCAAAACGTAATATTTGTGGTTTTAATGTTTCACAACATAAACGTAGTATTAATACATTAAAAGCTATTAAATCTTATGTATTAAATAATTGAAAACCAATTGATAATACACCATTATTAATTAAAAATAAATTATTAAAAGATTGAGATTCATCTATTAAATTAACTAAACCTGATAAAAATGGAGTTATTAAATTAGAATTTAATAGAATAGATTTTTTATATTATGTTATTTTACCTAAATTATATTCATTAAAATGATATAGTCGTAAAGAAATTGATTTCCAATTATGAAAAACACTTATAGAAATCTATATAAAAGGTTTACATAATACACTTAAAGGTTCTAATTTATTAAAATTAATTAATAATAATATTAATAAAAAAAGATATTATTCTAATTATAATATTTCTCCTTTCGGGGTTCCGGCTCCCGTGGCCGGGCCCCGATGATAAGCTGTCAAACATGAGAATTAATTC

COX2::ARG8m*-HH-sgRNA(Arg8m)-IM83-HDV-COX2

Blue: Upstream sequence of COX2; Green: COX2 5’UTR; Orange:ARG8^m^ (with STOP at Q79); Purple: V5; Light green: COX2 3’UTR; Yellow highlight: HH; Teal highlight: sgRNA-IM83(targeting ARG8^m^) Green highlight: HDV; Light blue: COX2

AAGCTTATCGGCGGGGACCCCGAAGGAGTATAAATAAAAATTAATAATATATTATATATATATTATATTAATAATAATAATAATAATAATAAATAATAACTCCTTGCTTCATACCTTTATAAATAAGGTAATCACTAATATATTATAATAATAATAATTATATTTATAATTTATATATTTATATATATAAAAATTATATATATTATATATAATCTAAATATTATATATTTTAATAAATATTAATATATATGATATGAATATTATTAGTTTTCGGGAAGCGGGAATTCATAAATTTTAATTAAAAGTAGTATTAACATATTATAAATAGACAAAAGAGTCTAAAGGTTAAGATTTATTCATatgttcaaaagatatttatcatcaacatcatcaagaagattcacatcaatcttagaagaaaaagcttttCAAgtaacaacatattcaagacctgaagatttatgtatcacaagaggtaaaaatgctaaattatatgatgatgtaaatggtaaagaatatatcgatttcacagctggtattgctgtaacagctttaggtcatgctaatcctaaagtagcCgaaatcttacatcaCTAAgctaataaattagtCcattcatcaaatttatatttcacaaaagaatgtttagatttatcagaaaaaatcgtagaaaaaacaaaaCAAttcggtggtCAAcatgatgcttcaagagtattcttatgtaattcaggtaccgaagctaatgaagctgcattaaaattcgctaaaaaacatggtatcatgaaaaatcctagtaaaCAAggtattgtagctttcgaaaattcattccatggtagaacaatgggtgctttatcagtaacaTGAaattcaaaatatagAacacctttcggtgatttagtacctcatgtatcattcttaaatttaaatgatgaaatgacaaaattaCAAtcatatatcgaaacaaaaaaagatgaaatcgctggtttaatcgtagaacctattCAAggtgaaggtggtgtatttcctgtagaagtagaaaaattaactggtttaaaaaaaatctgtCAAgataatgatgtaatcgtaattcatgatgaaattCAAtgtggtttaggtagatcaggtaaattaTGAgctcatgcttatttaccttcagaagctcatcctgatattttcacatcagctaaagcattaggtaatggtttccctattgctgctacaatcgtaaatgaaaaagtaaataatgctttaagagtaggtgatcatggtacaacatatggtggtaatcctttagcttgttcagtatcaaattatgtattagatacaattgctgatgaagcattcttaaaaCAAgtatcaaaaaaatcagatatcttaCAAaaaagattaagagaaatcCAAgctaaatatcctaatCAAatcaaaacaatcagaggtaaaggtttaatgttaggtgctgaatttgtagaacctccaacagaagtaatcaaaaaagctagagaattaggtttattaatcatcacagctggtaaatcaacagtaagattcgtacctgctttaacaatcgaagatgaattaatcgaagaaggtatggatgctttcgaaaaagctatcgaagctgtatatgctGGCAAGCCCAtccccaaccccttgttaggcttagaCAGCACCtaaCTCGAGTTAATATTTACTTATTATTAATATTTTTAATTATTAAAAATAATAATAATAATAATAATTATAATAATATTCTTAAATATAATAAAGATATAGATTTATATTCTATTCAATCACCTTATGAATTCTAAgctCTGATGAGTCCGTGAGGACGAAACGAGTAAGCTCGTCagcTTAgtgatgtaagatttgttttagagctagaaatagcaagttaaaataaggctagtccgttatcaacttgaaaaagtggcaccgagtcggtggtgcttttgatccgccttgttagctcagttggtagagcccctatgaggctccaagcGGCCGGCATGGTCCCAGCCTCCTCGCTGGCGCCGGCTGGGCAACATGCTTCGGCATGGCGAATGGGACGAATTCCGTAAGGAGTGAGGGACCCCTCCCTATACTAACGGGAGGGGGACCGAACCCCGAAGGAGTTTTATTTTTAGTATTTTATAAAATATATATTTATATGATTAATAATATTATATATATTATTTATAAAAATAATATATAATTTTAATTATTTTTAATAAAAAAAGGTGGGGTTTGGTAATATAATATTTTTATTTTATTTATAATATATAATAATAAATTATAAATAAATTTTAATTAAAAGTAGTATTAACATATTATAAATAGACAAAAGAGTCTAAAGGTTAAGATTTATTAAAATGTTAGATTTATTAAGATTACAATTAACAACATTCATTATGAATGATGTACCAACACCTTATGCATGTTATTTTCAGGATTCAGCAACACCAAATCAAGAAGGTATTTTAGAATTACATGATAATATTATGTTTTATTTATTAGTTATTTTAGGTTTAGTATCTTGAATGTTATATACAATTGTTATAACATATTCAAAAAATCCTATTGCATATAAATATATTAAACATGGACAAACTATTGAAGTTATTTGAACAATTTTTCCAGCTGTAATTTTATTAATTATTGCTTTCCCTTCATTTATTTTATTATATTTATGTGATGAAGTTATTTCACCAGCTATAACTATTAAAGCTATTGGATATCAATGATATTGAAAATATGAATATTCAGATTTTATTAATGATAGTGGTGAAACTGTTGAATTTGAATCATATGTTATTCCTGATGAATTATTAGAAGAAGGACAATTAAGATTATTAGATACTGATACTTCTATAGTTGTACCTGTAGATACACATATTAGATTCGTTGTAACAGCTGCTGATGTTATTCATGATTTTGCTATCCCAAGTTTAGGTATTAAAGTTGATGCTACTCCTGGTAGATTAAATCAAGTTTCTGCTTTAATTCAAAGAGAAGGTGTCTTCTATGGGGCATGTTCTGAGTTGTGTGGGACAGGTCATGCAAATATGCCAATTAAGATCGAAGCAGTATCATTACCTAAATTTTTGGAATGATTAAATGAACAATAATTAATATTTACTTATTATTAATATTTTTAATTATTAAAAATAATAATAATAATAATAATTATAATAATATTCTTAAATATAATAAAGATATAGATTTATATTCTATTCAATCACCTTATATTAAAAATATAAATATTATTAAAAGAGGTTATCATACTTCTTTAAATAATAAATTAATTATTGGTTCAAAAGGATAATAAAAATAATAATAAGAATAACCCAGAATTGATAATTTTTATAAATGATTAGTGGATTTACAGATGGAGATGGTAGTTTTTATATTAAATTAAATGATAAAAAATATTTAAGATTTTTTTATGGTTTTAGAATACATATTGATGATAAAGCATGTTTAGAAAAGATTAGAAATATATTAAATATACCTTCTAATTTTGAAGAACTACTTAAAACAATTATATTAGTAAATTCACAAAAGAAATGGTTATATTCTAATATTGTAACTATTTTTGATAAGTATCCTTGTTTAACAATTAAATATTATAGTTATTATAAATGAAAAATAGCTATAATTAATAATTTAAATGGTATATCTTATAATAATAAAGATTTATTAAATATTAAAAATACAATTAATAATTATGAAGTTATACCTAATTTAAAAATTCCATATGATAAAATAAATGATTATTGAATTTTAGGTTTTATTGAAGCTGAAGGTTCATTTGATCTATCTCCAAAACGTAATATTTGTGGTTTTAATGTTTCACAACATAAACGTAGTATTAATACATTAAAAGCTATTAAATCTTATGTATTAAATAATTGAAAACCAATTGATAATACACCATTATTAATTAAAAATAAATTATTAAAAGATTGAGATTCATCTATTAAATTAACTAAACCTGATAAAAATGGAGTTATTAAATTAGAATTTAATAGAATAGATTTTTTATATTATGTTATTTTACCTAAATTATATTCATTAAAATGATATAGTCGTAAAGAAATTGATTTCCAATTATGAAAAACACTTATAGAAATCTATATAAAAGGTTTACATAATACACTTAAAGGTTCTAATTTATTAAAATTAATTAATAATAATATTAATAAAAAAAGATATTATTCTAATTATAATATTTCTCCTTTCGGGGTTCCGGCTCCCGTGGCCGGGCCCCGATGATAAGCTGTCAAACATGAGAATTAATTC

COX2::ABE^m^-linker-ARG8^m^*-HH-sgRNA(Arg8^m^)-HDV-COX2

Blue: Upstream sequence of COX2; Green: COX2 5’UTR; Red: ABE^m^; Orange:ARG8^m^ (with STOP at Q79); Purple: V5; Light green: COX2 3’UTR; Yellow highlight: HH; Teal highlight: sgRNA(targeting ARG8^m^) Green highlight: HDV; Light blue: COX2

AAGCTTATCGGCGGGGACCCCGAAGGAGTATAAATAAAAATTAATAATATATTATATATATATTATATTAATAATAATAATAATAATAATAAATAATAACTCCTTGCTTCATACCTTTATAAATAAGGTAATCACTAATATATTATAATAATAATAATTATATTTATAATTTATATATTTATATATATAAAAATTATATATATTATATATAATCTAAATATTATATATTTTAATAAATATTAATATATATGATATGAATATTATTAGTTTTCGGGAAGCGGGAATTCATAAATTTTAATTAAAAGTAGTATTAACATATTATAAATAGACAAAAGAGTCTAAAGGTTAAGATTTATTCATATGTCTGAGGTGGAGTTTTCCcacgagtactggatgagacatgccTTAaccTTAgcAaagagggcacgggatgagagggaggtgcctgtgggagccgtgTTAgtgTTGaacaatagagtgatcggcgagggctggaacagagccatcggcTTAcacgacccaacagcccatgccgaaattatggccTTGagacagggcggcTTAgtcatgcagaactacagaTTGattgacgccaccTTGtacgtgacattcgagccttgcgtgatgtgcgccggcgccatgatccactctaggatcggccgcgtggtgtttggcgtgaggaactcaaaaagaggcgccgcaggctccctgatgaacgtgTTAaactaccccggcatgaatcaccgcgtcgaaattaccgagggaatcTTGgcagatgaatgtgccgccTTATTGtgcgatttctatcggatgcctagacaggtgttcaatgctcagaagaaggcccagagctccatcaactccggaggatctagcggaggctcctctggctctgagacacctggcacaagcgagagcgcaacacctgaaagcagcGGAggcTCTagcGGTgggtcagacaagaagtacagcatcggcTTGgccatcggcaccaactctgtgggcTGAgccgtgatcaccgacgagtacaaggtgcccagcaagaaattcaaggtgTTAggcaacaccgaccggcacagcatcaagaagaacTTGatcggagccTTGTTGttcgacagcggcgaaacagccgaggccacccggTTGaagagaaccgccagaagaagatacaccagacggaagaaccggatctgctatTTAcaagagatcttcagcaacgagatggccaaggtggacgacagcttcttccacagaTTAgaagagtccttcTTAgtggaagaggataagaagcacgagcggcaccccatcttcggcaacatcgtggacgaggtggcctaccacgagaagtaccccaccatctaccacTTAagaaagaaaTTGgtggacagcaccgacaaggccgacTTAcggTTGatctatTTAgccTTAgcccacatgatcaagttccggggccacttcTTAatcgagggcgacTTGaaccccgacaacagcgacgtggacaagTTGttcatccagTTAgtgcagacctacaaccagTTGttcgaggaaaaccccatcaacgccagcggcgtggacgccaaggccatcTTAtctgccagaTTGagcaagagcagacggTTGgaaaatTTGatcgcccagTTAcccggcgagaagaagaatggcTTGttcggaaacTTAattgccTTGagcTTAggcTTGacccccaacttcaagagcaacttcgacTTAgccgaggatgccaaaTTGcagTTAagcaaggacacctacgacgacgacTTGgacaacTTGTTAgcccagatcggcgaccagtacgccgacTTGtttTTAgccgccaagaacTTAtccgacgccatcTTGTTGagcgacatcTTGagagtgaacaccgagatcaccaaggcccccTTAagcgcctctatgatcaagagatacgacgagcaccaccaggacTTAaccTTGTTGaaagctTTAgtgcggcagcagTTGcctgagaagtacaaagagattttcttcgaccagagcaagaacggctacgccggctacattgacggcggagccagccaggaagagttctacaagttcatcaagcccatcTTAgaaaagatggacggcaccgaggaaTTATTGgtgaagTTGaacagagaggacTTATTAcggaagcagcggaccttcgacaacggcagcatcccccaccagatccacTTAggagagTTAcacgccattTTAcggcggcaggaagatttttacccattcTTGaaggacaaccgggaaaagatcgagaagatcTTAaccttccgcatcccctactacgtgggccctTTAgccaggggaaacagcagattcgccTGAatgaccagaaagagcgaggaaaccatcacccccTGAaacttcgaggaagtggtggacaagggcgcttccgcccagagcttcatcgagcggatgaccaacttcgataagaacTTGcccaacgagaaggtgTTAcccaagcacagcTTATTAtacgagtacttcaccgtgtataacgagTTGaccaaagtgaaatacgtgaccgagggaatgagaaagcccgccttcTTGagcggcgagcagaaaaaggccatcgtggacTTGTTGttcaagaccaaccggaaagtgaccgtgaagcagTTGaaagaggactacttcaagaaaatcgagtgcttcgactccgtggaaatctccggcgtggaagatcggttcaacgcctccTTAggcacataccacgatTTGTTGaaaattatcaaggacaaggacttcTTGgacaatgaggaaaacgaggacattTTAgaagatatcgtgTTGaccTTGacaTTGtttgaggacagagagatgatcgaggaacggTTAaaaacctatgcccacTTGttcgacgacaaagtgatgaagcagTTGaagcggcggagatacaccggcTGAggcaggctgagccggaagTTGatcaacggcatccgggacaagcagtccggcaagacaatcTTAgatttcTTGaagtccgacggcttcgccaacagaaacttcatgcagTTAatccacgacgacagcTTAacctttaaagaggacatccagaaagcccaggtgtccggccagggcgatagcTTGcacgagcacattgccaatTTAgccggcagccccgccattaagaagggcatcTTGcagacagtgaaggtggtggacgagTTAgtgaaagtgatgggccggcacaagcccgagaacatcgtgatcgaaatggccagagagaaccagaccacccagaagggacagaagaacagccgcgagagaatgaagcggatcgaagagggcatcaaagagTTAggcagccagatcTTGaaagaacaccccgtggaaaacacccagTTAcagaacgagaagTTGtacTTGtactacTTAcagaatgggcgggatatgtacgtggaccaggaaTTAgacatcaaccggTTAtccgactacgatgtggaccatatcgtgcctcagagctttTTGaaggacgactccatcgacaacaaggtgTTGaccagaagcgacaagaaccggggcaagagcgacaacgtgccctccgaagaggtcgtgaagaagatgaagaactacTGAcggcagTTGTTAaacgccaagTTGattacccagagaaagttcgacaatTTGaccaaggccgagagaggcggcTTGagcgaaTTAgataaggccggcttcatcaagagacagTTAgtggaaacccggcagatcacaaagcacgtggcacagatcTTAgactcccggatgaacactaagtacgacgagaatgacaagTTGatccgggaagtgaaagtgatcaccTTGaagtccaagTTAgtgtccgatttccggaaggatttccagttttacaaagtgcgcgagatcaacaactaccaccacgcccacgacgcctacTTGaacgccgtcgtgggaaccgccTTAatcaaaaagtaccctaagTTAgaaagcgagttcgtgtacggcgactacaaggtgtacgacgtgcggaagatgatcgccaagagcgagcaggaaatcggcaaggctaccgccaagtacttcttctacagcaacatcatgaactttttcaagaccgagattaccTTAgccaacggcgagatccggaagcggcctTTGatcgagacaaacggcgaaaccggggagatcgtgTGAgataagggccgggattttgccaccgtgcggaaagtgTTGagcatgccccaagtgaatatcgtgaaaaagaccgaggtgcagacaggcggcttcagcaaagagtctatcTTAcccaagaggaacagcgataagTTGatcgccagaaagaaggacTGGgaccctaagaagtacggcGGAttcgacTCTcccaccgtggcctattctgtgTTAgtggtggccaaagtggaaaagggcaagtccaagaaaTTGaagagtgtgaaagagTTGTTAgggatcaccatcatggaaagaagcagcttcgagaagaatcccatcgactttTTGgaagccaagggctacaaagaagtgaaaaaggacTTGatcatcaagTTAcctaagtactccTTAttcgagTTGgaaaacggccggaagagaatgTTAgcctctgccggcgaaTTAcagaagggaaacgaaTTAgccTTAccctccaaatatgtgaacttcTTGtacTTAgccagccactatgagaagTTGaagggctcccccgaggataatgagcagaaacagTTGtttgtggaacagcacaagcactacTTAgacgagatcatcgagcagatcagcgagttctccaagagagtgatcTTAgccgacgctaatTTAgacaaagtgTTAtccgcctacaacaagcaccgggataagcccatcagagagcaggccgagaatatcatccacTTGtttaccTTAaccaatTTAggagcccctgccgccttcaagtactttgacaccaccatcgaccggaagaggtacaccagcaccaaagaggtgTTAgacgccaccTTAatccaccagagcatcaccggcTTAtacgagacacggatcgacTTAtctcagTTAggagGTGACTctggcggctcattcaaaagatatttatcatcaacatcatcaagaagattcacatcaatcttagaagaaaaagcttttCAAgtaacaacatattcaagacctgaagatttatgtatcacaagaggtaaaaatgctaaattatatgatgatgtaaatggtaaagaatatatcgatttcacagctggtattgctgtaacagctttaggtcatgctaatcctaaagtagcCgaaatcttacatcaCTAAgctaataaattagtCcattcatcaaatttatatttcacaaaagaatgtttagatttatcagaaaaaatcgtagaaaaaacaaaaCAAttcggtggtCAAcatgatgcttcaagagtattcttatgtaattcaggtaccgaagctaatgaagctgcattaaaattcgctaaaaaacatggtatcatgaaaaatcctagtaaaCAAggtattgtagctttcgaaaattcattccatggtagaacaatgggtgctttatcagtaacaTGAaattcaaaatatagAacacctttcggtgatttagtacctcatgtatcattcttaaatttaaatgatgaaatgacaaaattaCAAtcatatatcgaaacaaaaaaagatgaaatcgctggtttaatcgtagaacctattCAAggtgaaggtggtgtatttcctgtagaagtagaaaaattaactggtttaaaaaaaatctgtCAAgataatgatgtaatcgtaattcatgatgaaattCAAtgtggtttaggtagatcaggtaaattaTGAgctcatgcttatttaccttcagaagctcatcctgatattttcacatcagctaaagcattaggtaatggtttccctattgctgctacaatcgtaaatgaaaaagtaaataatgctttaagagtaggtgatcatggtacaacatatggtggtaatcctttagcttgttcagtatcaaattatgtattagatacaattgctgatgaagcattcttaaaaCAAgtatcaaaaaaatcagatatcttaCAAaaaagattaagagaaatcCAAgctaaatatcctaatCAAatcaaaacaatcagaggtaaaggtttaatgttaggtgctgaatttgtagaacctccaacagaagtaatcaaaaaagctagagaattaggtttattaatcatcacagctggtaaatcaacagtaagattcgtacctgctttaacaatcgaagatgaattaatcgaagaaggtatggatgctttcgaaaaagctatcgaagctgtatatgctGGCAAGCCCAtccccaaccccttgttaggcttagaCAGCACCtaaCTCGAGTTAATATTTACTTATTATTAATATTTTTAATTATTAAAAATAATAATAATAATAATAATTATAATAATATTCTTAAATATAATAAAGATATAGATTTATATTCTATTCAATCACCTTATGAATTCTAAgctCTGATGAGTCCGTGAGGACGAAACGAGTAAGCTCGTCagcTTAgtgatgtaagatttgttttagagctagaaatagcaagttaaaataaggctagtccgttatcaacttgaaaaagtggcaccgagtcggtggtgcttttGGCCGGCATGGTCCCAGCCTCCTCGCTGGCGCCGGCTGGGCAACATGCTTCGGCATGGCGAATGGGACGAATTCCGTAAGGAGTGAGGGACCCCTCCCTATACTAACGGGAGGGGGACCGAACCCCGAAGGAGTTTTATTTTTAGTATTTTATAAAATATATATTTATATGATTAATAATATTATATATATTATTTATAAAAATAATATATAATTTTAATTATTTTTAATAAAAAAAGGTGGGGTTTGGTAATATAATATTTTTATTTTATTTATAATATATAATAATAAATTATAAATAAATTTTAATTAAAAGTAGTATTAACATATTATAAATAGACAAAAGAGTCTAAAGGTTAAGATTTATTAAAATGTTAGATTTATTAAGATTACAATTAACAACATTCATTATGAATGATGTACCAACACCTTATGCATGTTATTTTCAGGATTCAGCAACACCAAATCAAGAAGGTATTTTAGAATTACATGATAATATTATGTTTTATTTATTAGTTATTTTAGGTTTAGTATCTTGAATGTTATATACAATTGTTATAACATATTCAAAAAATCCTATTGCATATAAATATATTAAACATGGACAAACTATTGAAGTTATTTGAACAATTTTTCCAGCTGTAATTTTATTAATTATTGCTTTCCCTTCATTTATTTTATTATATTTATGTGATGAAGTTATTTCACCAGCTATAACTATTAAAGCTATTGGATATCAATGATATTGAAAATATGAATATTCAGATTTTATTAATGATAGTGGTGAAACTGTTGAATTTGAATCATATGTTATTCCTGATGAATTATTAGAAGAAGGACAATTAAGATTATTAGATACTGATACTTCTATAGTTGTACCTGTAGATACACATATTAGATTCGTTGTAACAGCTGCTGATGTTATTCATGATTTTGCTATCCCAAGTTTAGGTATTAAAGTTGATGCTACTCCTGGTAGATTAAATCAAGTTTCTGCTTTAATTCAAAGAGAAGGTGTCTTCTATGGGGCATGTTCTGAGTTGTGTGGGACAGGTCATGCAAATATGCCAATTAAGATCGAAGCAGTATCATTACCTAAATTTTTGGAATGATTAAATGAACAATAATTAATATTTACTTATTATTAATATTTTTAATTATTAAAAATAATAATAATAATAATAATTATAATAATATTCTTAAATATAATAAAGATATAGATTTATATTCTATTCAATCACCTTATATTAAAAATATAAATATTATTAAAAGAGGTTATCATACTTCTTTAAATAATAAATTAATTATTGGTTCAAAAGGATAATAAAAATAATAATAAGAATAACCCAGAATTGATAATTTTTATAAATGATTAGTGGATTTACAGATGGAGATGGTAGTTTTTATATTAAATTAAATGATAAAAAATATTTAAGATTTTTTTATGGTTTTAGAATACATATTGATGATAAAGCATGTTTAGAAAAGATTAGAAATATATTAAATATACCTTCTAATTTTGAAGAACTACTTAAAACAATTATATTAGTAAATTCACAAAAGAAATGGTTATATTCTAATATTGTAACTATTTTTGATAAGTATCCTTGTTTAACAATTAAATATTATAGTTATTATAAATGAAAAATAGCTATAATTAATAATTTAAATGGTATATCTTATAATAATAAAGATTTATTAAATATTAAAAATACAATTAATAATTATGAAGTTATACCTAATTTAAAAATTCCATATGATAAAATAAATGATTATTGAATTTTAGGTTTTATTGAAGCTGAAGGTTCATTTGATCTATCTCCAAAACGTAATATTTGTGGTTTTAATGTTTCACAACATAAACGTAGTATTAATACATTAAAAGCTATTAAATCTTATGTATTAAATAATTGAAAACCAATTGATAATACACCATTATTAATTAAAAATAAATTATTAAAAGATTGAGATTCATCTATTAAATTAACTAAACCTGATAAAAATGGAGTTATTAAATTAGAATTTAATAGAATAGATTTTTTATATTATGTTATTTTACCTAAATTATATTCATTAAAATGATATAGTCGTAAAGAAATTGATTTCCAATTATGAAAAACACTTATAGAAATCTATATAAAAGGTTTACATAATACACTTAAAGGTTCTAATTTATTAAAATTAATTAATAATAATATTAATAAAAAAAGATATTATTCTAATTATAATATTTCTCCTTTCGGGGTTCCGGCTCCCGTGGCCGGGCCCCGATGATAAGCTGTCAAACATGAGAATTAATTC

COX2::ABE^m^-linker-ARG8^m^-COX2

Blue: Upstream sequence of COX2; Green: COX2 5’UTR; Red: ABE^m^; Orange:ARG8^m^; Purple: V5; Light green: COX2 3’UTR; Light blue: COX2

AAGCTTATCGGCGGGGACCCCGAAGGAGTATAAATAAAAATTAATAATATATTATATATATATTATATTAATAATAATAATAATAATAATAAATAATAACTCCTTGCTTCATACCTTTATAAATAAGGTAATCACTAATATATTATAATAATAATAATTATATTTATAATTTATATATTTATATATATAAAAATTATATATATTATATATAATCTAAATATTATATATTTTAATAAATATTAATATATATGATATGAATATTATTAGTTTTCGGGAAGCGGGAATTCATAAATTTTAATTAAAAGTAGTATTAACATATTATAAATAGACAAAAGAGTCTAAAGGTTAAGATTTATTCATATGTCTGAGGTGGAGTTTTCCcacgagtactggatgagacatgccTTAaccTTAgcAaagagggcacgggatgagagggaggtgcctgtgggagccgtgTTAgtgTTGaacaatagagtgatcggcgagggctggaacagagccatcggcTTAcacgacccaacagcccatgccgaaattatggccTTGagacagggcggcTTAgtcatgcagaactacagaTTGattgacgccaccTTGtacgtgacattcgagccttgcgtgatgtgcgccggcgccatgatccactctaggatcggccgcgtggtgtttggcgtgaggaactcaaaaagaggcgccgcaggctccctgatgaacgtgTTAaactaccccggcatgaatcaccgcgtcgaaattaccgagggaatcTTGgcagatgaatgtgccgccTTATTGtgcgatttctatcggatgcctagacaggtgttcaatgctcagaagaaggcccagagctccatcaactccggaggatctagcggaggctcctctggctctgagacacctggcacaagcgagagcgcaacacctgaaagcagcGGAggcTCTagcGGTgggtcagacaagaagtacagcatcggcTTGgccatcggcaccaactctgtgggcTGAgccgtgatcaccgacgagtacaaggtgcccagcaagaaattcaaggtgTTAggcaacaccgaccggcacagcatcaagaagaacTTGatcggagccTTGTTGttcgacagcggcgaaacagccgaggccacccggTTGaagagaaccgccagaagaagatacaccagacggaagaaccggatctgctatTTAcaagagatcttcagcaacgagatggccaaggtggacgacagcttcttccacagaTTAgaagagtccttcTTAgtggaagaggataagaagcacgagcggcaccccatcttcggcaacatcgtggacgaggtggcctaccacgagaagtaccccaccatctaccacTTAagaaagaaaTTGgtggacagcaccgacaaggccgacTTAcggTTGatctatTTAgccTTAgcccacatgatcaagttccggggccacttcTTAatcgagggcgacTTGaaccccgacaacagcgacgtggacaagTTGttcatccagTTAgtgcagacctacaaccagTTGttcgaggaaaaccccatcaacgccagcggcgtggacgccaaggccatcTTAtctgccagaTTGagcaagagcagacggTTGgaaaatTTGatcgcccagTTAcccggcgagaagaagaatggcTTGttcggaaacTTAattgccTTGagcTTAggcTTGacccccaacttcaagagcaacttcgacTTAgccgaggatgccaaaTTGcagTTAagcaaggacacctacgacgacgacTTGgacaacTTGTTAgcccagatcggcgaccagtacgccgacTTGtttTTAgccgccaagaacTTAtccgacgccatcTTGTTGagcgacatcTTGagagtgaacaccgagatcaccaaggcccccTTAagcgcctctatgatcaagagatacgacgagcaccaccaggacTTAaccTTGTTGaaagctTTAgtgcggcagcagTTGcctgagaagtacaaagagattttcttcgaccagagcaagaacggctacgccggctacattgacggcggagccagccaggaagagttctacaagttcatcaagcccatcTTAgaaaagatggacggcaccgaggaaTTATTGgtgaagTTGaacagagaggacTTATTAcggaagcagcggaccttcgacaacggcagcatcccccaccagatccacTTAggagagTTAcacgccattTTAcggcggcaggaagatttttacccattcTTGaaggacaaccgggaaaagatcgagaagatcTTAaccttccgcatcccctactacgtgggccctTTAgccaggggaaacagcagattcgccTGAatgaccagaaagagcgaggaaaccatcacccccTGAaacttcgaggaagtggtggacaagggcgcttccgcccagagcttcatcgagcggatgaccaacttcgataagaacTTGcccaacgagaaggtgTTAcccaagcacagcTTATTAtacgagtacttcaccgtgtataacgagTTGaccaaagtgaaatacgtgaccgagggaatgagaaagcccgccttcTTGagcggcgagcagaaaaaggccatcgtggacTTGTTGttcaagaccaaccggaaagtgaccgtgaagcagTTGaaagaggactacttcaagaaaatcgagtgcttcgactccgtggaaatctccggcgtggaagatcggttcaacgcctccTTAggcacataccacgatTTGTTGaaaattatcaaggacaaggacttcTTGgacaatgaggaaaacgaggacattTTAgaagatatcgtgTTGaccTTGacaTTGtttgaggacagagagatgatcgaggaacggTTAaaaacctatgcccacTTGttcgacgacaaagtgatgaagcagTTGaagcggcggagatacaccggcTGAggcaggctgagccggaagTTGatcaacggcatccgggacaagcagtccggcaagacaatcTTAgatttcTTGaagtccgacggcttcgccaacagaaacttcatgcagTTAatccacgacgacagcTTAacctttaaagaggacatccagaaagcccaggtgtccggccagggcgatagcTTGcacgagcacattgccaatTTAgccggcagccccgccattaagaagggcatcTTGcagacagtgaaggtggtggacgagTTAgtgaaagtgatgggccggcacaagcccgagaacatcgtgatcgaaatggccagagagaaccagaccacccagaagggacagaagaacagccgcgagagaatgaagcggatcgaagagggcatcaaagagTTAggcagccagatcTTGaaagaacaccccgtggaaaacacccagTTAcagaacgagaagTTGtacTTGtactacTTAcagaatgggcgggatatgtacgtggaccaggaaTTAgacatcaaccggTTAtccgactacgatgtggaccatatcgtgcctcagagctttTTGaaggacgactccatcgacaacaaggtgTTGaccagaagcgacaagaaccggggcaagagcgacaacgtgccctccgaagaggtcgtgaagaagatgaagaactacTGAcggcagTTGTTAaacgccaagTTGattacccagagaaagttcgacaatTTGaccaaggccgagagaggcggcTTGagcgaaTTAgataaggccggcttcatcaagagacagTTAgtggaaacccggcagatcacaaagcacgtggcacagatcTTAgactcccggatgaacactaagtacgacgagaatgacaagTTGatccgggaagtgaaagtgatcaccTTGaagtccaagTTAgtgtccgatttccggaaggatttccagttttacaaagtgcgcgagatcaacaactaccaccacgcccacgacgcctacTTGaacgccgtcgtgggaaccgccTTAatcaaaaagtaccctaagTTAgaaagcgagttcgtgtacggcgactacaaggtgtacgacgtgcggaagatgatcgccaagagcgagcaggaaatcggcaaggctaccgccaagtacttcttctacagcaacatcatgaactttttcaagaccgagattaccTTAgccaacggcgagatccggaagcggcctTTGatcgagacaaacggcgaaaccggggagatcgtgTGAgataagggccgggattttgccaccgtgcggaaagtgTTGagcatgccccaagtgaatatcgtgaaaaagaccgaggtgcagacaggcggcttcagcaaagagtctatcTTAcccaagaggaacagcgataagTTGatcgccagaaagaaggacTGGgaccctaagaagtacggcGGAttcgacTCTcccaccgtggcctattctgtgTTAgtggtggccaaagtggaaaagggcaagtccaagaaaTTGaagagtgtgaaagagTTGTTAgggatcaccatcatggaaagaagcagcttcgagaagaatcccatcgactttTTGgaagccaagggctacaaagaagtgaaaaaggacTTGatcatcaagTTAcctaagtactccTTAttcgagTTGgaaaacggccggaagagaatgTTAgcctctgccggcgaaTTAcagaagggaaacgaaTTAgccTTAccctccaaatatgtgaacttcTTGtacTTAgccagccactatgagaagTTGaagggctcccccgaggataatgagcagaaacagTTGtttgtggaacagcacaagcactacTTAgacgagatcatcgagcagatcagcgagttctccaagagagtgatcTTAgccgacgctaatTTAgacaaagtgTTAtccgcctacaacaagcaccgggataagcccatcagagagcaggccgagaatatcatccacTTGtttaccTTAaccaatTTAggagcccctgccgccttcaagtactttgacaccaccatcgaccggaagaggtacaccagcaccaaagaggtgTTAgacgccaccTTAatccaccagagcatcaccggcTTAtacgagacacggatcgacTTAtctcagTTAggagGTGACTctggcggctcattcaaaagatatttatcatcaacatcatcaagaagattcacatcaatcttagaagaaaaagcttttCAAgtaacaacatattcaagacctgaagatttatgtatcacaagaggtaaaaatgctaaattatatgatgatgtaaatggtaaagaatatatcgatttcacagctggtattgctgtaacagctttaggtcatgctaatcctaaagtagcCgaaatcttacatcaCCAAgctaataaattagtCcattcatcaaatttatatttcacaaaagaatgtttagatttatcagaaaaaatcgtagaaaaaacaaaaCAAttcggtggtCAAcatgatgcttcaagagtattcttatgtaattcaggtaccgaagctaatgaagctgcattaaaattcgctaaaaaacatggtatcatgaaaaatcctagtaaaCAAggtattgtagctttcgaaaattcattccatggtagaacaatgggtgctttatcagtaacaTGAaattcaaaatatagAacacctttcggtgatttagtacctcatgtatcattcttaaatttaaatgatgaaatgacaaaattaCAAtcatatatcgaaacaaaaaaagatgaaatcgctggtttaatcgtagaacctattCAAggtgaaggtggtgtatttcctgtagaagtagaaaaattaactggtttaaaaaaaatctgtCAAgataatgatgtaatcgtaattcatgatgaaattCAAtgtggtttaggtagatcaggtaaattaTGAgctcatgcttatttaccttcagaagctcatcctgatattttcacatcagctaaagcattaggtaatggtttccctattgctgctacaatcgtaaatgaaaaagtaaataatgctttaagagtaggtgatcatggtacaacatatggtggtaatcctttagcttgttcagtatcaaattatgtattagatacaattgctgatgaagcattcttaaaaCAAgtatcaaaaaaatcagatatcttaCAAaaaagattaagagaaatcCAAgctaaatatcctaatCAAatcaaaacaatcagaggtaaaggtttaatgttaggtgctgaatttgtagaacctccaacagaagtaatcaaaaaagctagagaattaggtttattaatcatcacagctggtaaatcaacagtaagattcgtacctgctttaacaatcgaagatgaattaatcgaagaaggtatggatgctttcgaaaaagctatcgaagctgtatatgctGGCAAGCCCAtccccaaccccttgttaggcttagaCAGCACCtaaCTCGAGTTAATATTTACTTATTATTAATATTTTTAATTATTAAAAATAATAATAATAATAATAATTATAATAATATTCTTAAATATAATAAAGATATAGATTTATATTCTATTCAATCACCTTATGAATTCGAATTCCGTAAGGAGTGAGGGACCCCTCCCTATACTAACGGGAGGGGGACCGAACCCCGAAGGAGTTTTATTTTTAGTATTTTATAAAATATATATTTATATGATTAATAATATTATATATATTATTTATAAAAATAATATATAATTTTAATTATTTTTAATAAAAAAAGGTGGGGTTTGGTAATATAATATTTTTATTTTATTTATAATATATAATAATAAATTATAAATAAATTTTAATTAAAAGTAGTATTAACATATTATAAATAGACAAAAGAGTCTAAAGGTTAAGATTTATTAAAATGTTAGATTTATTAAGATTACAATTAACAACATTCATTATGAATGATGTACCAACACCTTATGCATGTTATTTTCAGGATTCAGCAACACCAAATCAAGAAGGTATTTTAGAATTACATGATAATATTATGTTTTATTTATTAGTTATTTTAGGTTTAGTATCTTGAATGTTATATACAATTGTTATAACATATTCAAAAAATCCTATTGCATATAAATATATTAAACATGGACAAACTATTGAAGTTATTTGAACAATTTTTCCAGCTGTAATTTTATTAATTATTGCTTTCCCTTCATTTATTTTATTATATTTATGTGATGAAGTTATTTCACCAGCTATAACTATTAAAGCTATTGGATATCAATGATATTGAAAATATGAATATTCAGATTTTATTAATGATAGTGGTGAAACTGTTGAATTTGAATCATATGTTATTCCTGATGAATTATTAGAAGAAGGACAATTAAGATTATTAGATACTGATACTTCTATAGTTGTACCTGTAGATACACATATTAGATTCGTTGTAACAGCTGCTGATGTTATTCATGATTTTGCTATCCCAAGTTTAGGTATTAAAGTTGATGCTACTCCTGGTAGATTAAATCAAGTTTCTGCTTTAATTCAAAGAGAAGGTGTCTTCTATGGGGCATGTTCTGAGTTGTGTGGGACAGGTCATGCAAATATGCCAATTAAGATCGAAGCAGTATCATTACCTAAATTTTTGGAATGATTAAATGAACAATAATTAATATTTACTTATTATTAATATTTTTAATTATTAAAAATAATAATAATAATAATAATTATAATAATATTCTTAAATATAATAAAGATATAGATTTATATTCTATTCAATCACCTTATATTAAAAATATAAATATTATTAAAAGAGGTTATCATACTTCTTTAAATAATAAATTAATTATTGGTTCAAAAGGATAATAAAAATAATAATAAGAATAACCCAGAATTGATAATTTTTATAAATGATTAGTGGATTTACAGATGGAGATGGTAGTTTTTATATTAAATTAAATGATAAAAAATATTTAAGATTTTTTTATGGTTTTAGAATACATATTGATGATAAAGCATGTTTAGAAAAGATTAGAAATATATTAAATATACCTTCTAATTTTGAAGAACTACTTAAAACAATTATATTAGTAAATTCACAAAAGAAATGGTTATATTCTAATATTGTAACTATTTTTGATAAGTATCCTTGTTTAACAATTAAATATTATAGTTATTATAAATGAAAAATAGCTATAATTAATAATTTAAATGGTATATCTTATAATAATAAAGATTTATTAAATATTAAAAATACAATTAATAATTATGAAGTTATACCTAATTTAAAAATTCCATATGATAAAATAAATGATTATTGAATTTTAGGTTTTATTGAAGCTGAAGGTTCATTTGATCTATCTCCAAAACGTAATATTTGTGGTTTTAATGTTTCACAACATAAACGTAGTATTAATACATTAAAAGCTATTAAATCTTATGTATTAAATAATTGAAAACCAATTGATAATACACCATTATTAATTAAAAATAAATTATTAAAAGATTGAGATTCATCTATTAAATTAACTAAACCTGATAAAAATGGAGTTATTAAATTAGAATTTAATAGAATAGATTTTTTATATTATGTTATTTTACCTAAATTATATTCATTAAAATGATATAGTCGTAAAGAAATTGATTTCCAATTATGAAAAACACTTATAGAAATCTATATAAAAGGTTTACATAATACACTTAAAGGTTCTAATTTATTAAAATTAATTAATAATAATATTAATAAAAAAAGATATTATTCTAATTATAATATTTCTCCTTTCGGGGTTCCGGCTCCCGTGGCCGGGCCCCGATGATAAGCTGTCAAACATGAGAATTAATTC

COX2::COX2 5’UTR - ABE^m^- COX2 3’UTR - COX2 5’UTR – ARG8^m^- COX2 3’UTR -COX2

Blue: Upstream sequence of COX2; Green: COX2 5’UTR; Red: ABE^m^; Orange:ARG8^m^; Purple: V5; Light green: COX2 3’UTR; Light blue: COX2

AAGCTTATCGGCGGGGACCCCGAAGGAGTATAAATAAAAATTAATAATATATTATATATATATTATATTAATAATAATAATAATAATAATAAATAATAACTCCTTGCTTCATACCTTTATAAATAAGGTAATCACTAATATATTATAATAATAATAATTATATTTATAATTTATATATTTATATATATAAAAATTATATATATTATATATAATCTAAATATTATATATTTTAATAAATATTAATATATATGATATGAATATTATTAGTTTTCGGGAAGCGGGAATTCATAAATTTTAATTAAAAGTAGTATTAACATATTATAAATAGACAAAAGAGTCTAAAGGTTAAGATTTATTCAT ATGTCTGAGGTGGAGTTTTCCcacgagtactggatgagacatgccTTAaccTTAgcAaagagggcacgggatgagagggaggtgcctgtgggagccgtgTTAgtgTTGaacaatagagtgatcggcgagggctggaacagagccatcggcTTAcacgacccaacagcccatgccgaaattatggccTTGagacagggcggcTTAgtcatgcagaactacagaTTGattgacgccaccTTGtacgtgacattcgagccttgcgtgatgtgcgccggcgccatgatccactctaggatcggccgcgtggtgtttggcgtgaggaactcaaaaagaggcgccgcaggctccctgatgaacgtgTTAaactaccccggcatgaatcaccgcgtcgaaattaccgagggaatcTTGgcagatgaatgtgccgccTTATTGtgcgatttctatcggatgcctagacaggtgttcaatgctcagaagaaggcccagagctccatcaactccggaggatctagcggaggctcctctggctctgagacacctggcacaagcgagagcgcaacacctgaaagcagcGGAggcTCTagcGGTgggtcagacaagaagtacagcatcggcTTGgccatcggcaccaactctgtgggcTGAgccgtgatcaccgacgagtacaaggtgcccagcaagaaattcaaggtgTTAggcaacaccgaccggcacagcatcaagaagaacTTGatcggagccTTGTTGttcgacagcggcgaaacagccgaggccacccggTTGaagagaaccgccagaagaagatacaccagacggaagaaccggatctgctatTTAcaagagatcttcagcaacgagatggccaaggtggacgacagcttcttccacagaTTAgaagagtccttcTTAgtggaagaggataagaagcacgagcggcaccccatcttcggcaacatcgtggacgaggtggcctaccacgagaagtaccccaccatctaccacTTAagaaagaaaTTGgtggacagcaccgacaaggccgacTTAcggTTGatctatTTAgccTTAgcccacatgatcaagttccggggccacttcTTAatcgagggcgacTTGaaccccgacaacagcgacgtggacaagTTGttcatccagTTAgtgcagacctacaaccagTTGttcgaggaaaaccccatcaacgccagcggcgtggacgccaaggccatcTTAtctgccagaTTGagcaagagcagacggTTGgaaaatTTGatcgcccagTTAcccggcgagaagaagaatggcTTGttcggaaacTTAattgccTTGagcTTAggcTTGacccccaacttcaagagcaacttcgacTTAgccgaggatgccaaaTTGcagTTAagcaaggacacctacgacgacgacTTGgacaacTTGTTAgcccagatcggcgaccagtacgccgacTTGtttTTAgccgccaagaacTTAtccgacgccatcTTGTTGagcgacatcTTGagagtgaacaccgagatcaccaaggcccccTTAagcgcctctatgatcaagagatacgacgagcaccaccaggacTTAaccTTGTTGaaagctTTAgtgcggcagcagTTGcctgagaagtacaaagagattttcttcgaccagagcaagaacggctacgccggctacattgacggcggagccagccaggaagagttctacaagttcatcaagcccatcTTAgaaaagatggacggcaccgaggaaTTATTGgtgaagTTGaacagagaggacTTATTAcggaagcagcggaccttcgacaacggcagcatcccccaccagatccacTTAggagagTTAcacgccattTTAcggcggcaggaagatttttacccattcTTGaaggacaaccgggaaaagatcgagaagatcTTAaccttccgcatcccctactacgtgggccctTTAgccaggggaaacagcagattcgccTGAatgaccagaaagagcgaggaaaccatcacccccTGAaacttcgaggaagtggtggacaagggcgcttccgcccagagcttcatcgagcggatgaccaacttcgataagaacTTGcccaacgagaaggtgTTAcccaagcacagcTTATTAtacgagtacttcaccgtgtataacgagTTGaccaaagtgaaatacgtgaccgagggaatgagaaagcccgccttcTTGagcggcgagcagaaaaaggccatcgtggacTTGTTGttcaagaccaaccggaaagtgaccgtgaagcagTTGaaagaggactacttcaagaaaatcgagtgcttcgactccgtggaaatctccggcgtggaagatcggttcaacgcctccTTAggcacataccacgatTTGTTGaaaattatcaaggacaaggacttcTTGgacaatgaggaaaacgaggacattTTAgaagatatcgtgTTGaccTTGacaTTGtttgaggacagagagatgatcgaggaacggTTAaaaacctatgcccacTTGttcgacgacaaagtgatgaagcagTTGaagcggcggagatacaccggcTGAggcaggctgagccggaagTTGatcaacggcatccgggacaagcagtccggcaagacaatcTTAgatttcTTGaagtccgacggcttcgccaacagaaacttcatgcagTTAatccacgacgacagcTTAacctttaaagaggacatccagaaagcccaggtgtccggccagggcgatagcTTGcacgagcacattgccaatTTAgccggcagccccgccattaagaagggcatcTTGcagacagtgaaggtggtggacgagTTAgtgaaagtgatgggccggcacaagcccgagaacatcgtgatcgaaatggccagagagaaccagaccacccagaagggacagaagaacagccgcgagagaatgaagcggatcgaagagggcatcaaagagTTAggcagccagatcTTGaaagaacaccccgtggaaaacacccagTTAcagaacgagaagTTGtacTTGtactacTTAcagaatgggcgggatatgtacgtggaccaggaaTTAgacatcaaccggTTAtccgactacgatgtggaccatatcgtgcctcagagctttTTGaaggacgactccatcgacaacaaggtgTTGaccagaagcgacaagaaccggggcaagagcgacaacgtgccctccgaagaggtcgtgaagaagatgaagaactacTGAcggcagTTGTTAaacgccaagTTGattacccagagaaagttcgacaatTTGaccaaggccgagagaggcggcTTGagcgaaTTAgataaggccggcttcatcaagagacagTTAgtggaaacccggcagatcacaaagcacgtggcacagatcTTAgactcccggatgaacactaagtacgacgagaatgacaagTTGatccgggaagtgaaagtgatcaccTTGaagtccaagTTAgtgtccgatttccggaaggatttccagttttacaaagtgcgcgagatcaacaactaccaccacgcccacgacgcctacTTGaacgccgtcgtgggaaccgccTTAatcaaaaagtaccctaagTTAgaaagcgagttcgtgtacggcgactacaaggtgtacgacgtgcggaagatgatcgccaagagcgagcaggaaatcggcaaggctaccgccaagtacttcttctacagcaacatcatgaactttttcaagaccgagattaccTTAgccaacggcgagatccggaagcggcctTTGatcgagacaaacggcgaaaccggggagatcgtgTGAgataagggccgggattttgccaccgtgcggaaagtgTTGagcatgccccaagtgaatatcgtgaaaaagaccgaggtgcagacaggcggcttcagcaaagagtctatcTTAcccaagaggaacagcgataagTTGatcgccagaaagaaggacTGGgaccctaagaagtacggcGGAttcgacTCTcccaccgtggcctattctgtgTTAgtggtggccaaagtggaaaagggcaagtccaagaaaTTGaagagtgtgaaagagTTGTTAgggatcaccatcatggaaagaagcagcttcgagaagaatcccatcgactttTTGgaagccaagggctacaaagaagtgaaaaaggacTTGatcatcaagTTAcctaagtactccTTAttcgagTTGgaaaacggccggaagagaatgTTAgcctctgccggcgaaTTAcagaagggaaacgaaTTAgccTTAccctccaaatatgtgaacttcTTGtacTTAgccagccactatgagaagTTGaagggctcccccgaggataatgagcagaaacagTTGtttgtggaacagcacaagcactacTTAgacgagatcatcgagcagatcagcgagttctccaagagagtgatcTTAgccgacgctaatTTAgacaaagtgTTAtccgcctacaacaagcaccgggataagcccatcagagagcaggccgagaatatcatccacTTGtttaccTTAaccaatTTAggagcccctgccgccttcaagtactttgacaccaccatcgaccggaagaggtacaccagcaccaaagaggtgTTAgacgccaccTTAatccaccagagcatcaccggcTTAtacgagacacggatcgacTTAtctcagTTAggagGTGACTctggcggctcaGGCAAGCCCAtccccaaccccttgttaggcttagaCAGCACCtaaCTCGAGTTAATATTTACTTATTATTAATATTTTTAATTATTAAAAATAATAATAATAATAATAATTATAATAATATTCTTAAATATAATAAAGATATAGATTTATATTCTATTCAATCACCTTATGAATTCATAAATTTTAATTAAAAGTAGTATTAACATATTATAAATAGACAAAAGAGTCTAAAGGTTAAGATTTATTCATatgttcaaaagatatttatcatcaacatcatcaagaagattcacatcaatcttagaagaaaaagcttttCAAgtaacaacatattcaagacctgaagatttatgtatcacaagaggtaaaaatgctaaattatatgatgatgtaaatggtaaagaatatatcgatttcacagctggtattgctgtaacagctttaggtcatgctaatcctaaagtagcCgaaatcttacatcaCCAAgctaataaattagtCcattcatcaaatttatatttcacaaaagaatgtttagatttatcagaaaaaatcgtagaaaaaacaaaaCAAttcggtggtCAAcatgatgcttcaagagtattcttatgtaattcaggtaccgaagctaatgaagctgcattaaaattcgctaaaaaacatggtatcatgaaaaatcctagtaaaCAAggtattgtagctttcgaaaattcattccatggtagaacaatgggtgctttatcagtaacaTGAaattcaaaatatagAacacctttcggtgatttagtacctcatgtatcattcttaaatttaaatgatgaaatgacaaaattaCAAtcatatatcgaaacaaaaaaagatgaaatcgctggtttaatcgtagaacctattCAAggtgaaggtggtgtatttcctgtagaagtagaaaaattaactggtttaaaaaaaatctgtCAAgataatgatgtaatcgtaattcatgatgaaattCAAtgtggtttaggtagatcaggtaaattaTGAgctcatgcttatttaccttcagaagctcatcctgatattttcacatcagctaaagcattaggtaatggtttccctattgctgctacaatcgtaaatgaaaaagtaaataatgctttaagagtaggtgatcatggtacaacatatggtggtaatcctttagcttgttcagtatcaaattatgtattagatacaattgctgatgaagcattcttaaaaCAAgtatcaaaaaaatcagatatcttaCAAaaaagattaagagaaatcCAAgctaaatatcctaatCAAatcaaaacaatcagaggtaaaggtttaatgttaggtgctgaatttgtagaacctccaacagaagtaatcaaaaaagctagagaattaggtttattaatcatcacagctggtaaatcaacagtaagattcgtacctgctttaacaatcgaagatgaattaatcgaagaaggtatggatgctttcgaaaaagctatcgaagctgtatatgctGGCAAGCCCAtccccaaccccttgttaggcttagaCAGCACCtaaCTCGAGTTAATATTTACTTATTATTAATATTTTTAATTATTAAAAATAATAATAATAATAATAATTATAATAATATTCTTAAATATAATAAAGATATAGATTTATATTCTATTCAATCACCTTATGAATTCGAATTCCGTAAGGAGTGAGGGACCCCTCCCTATACTAACGGGAGGGGGACCGAACCCCGAAGGAGTTTTATTTTTAGTATTTTATAAAATATATATTTATATGATTAATAATATTATATATATTATTTATAAAAATAATATATAATTTTAATTATTTTTAATAAAAAAAGGTGGGGTTTGGTAATATAATATTTTTATTTTATTTATAATATATAATAATAAATTATAAATAAATTTTAATTAAAAGTAGTATTAACATATTATAAATAGACAAAAGAGTCTAAAGGTTAAGATTTATTAAAATGTTAGATTTATTAAGATTACAATTAACAACATTCATTATGAATGATGTACCAACACCTTATGCATGTTATTTTCAGGATTCAGCAACACCAAATCAAGAAGGTATTTTAGAATTACATGATAATATTATGTTTTATTTATTAGTTATTTTAGGTTTAGTATCTTGAATGTTATATACAATTGTTATAACATATTCAAAAAATCCTATTGCATATAAATATATTAAACATGGACAAACTATTGAAGTTATTTGAACAATTTTTCCAGCTGTAATTTTATTAATTATTGCTTTCCCTTCATTTATTTTATTATATTTATGTGATGAAGTTATTTCACCAGCTATAACTATTAAAGCTATTGGATATCAATGATATTGAAAATATGAATATTCAGATTTTATTAATGATAGTGGTGAAACTGTTGAATTTGAATCATATGTTATTCCTGATGAATTATTAGAAGAAGGACAATTAAGATTATTAGATACTGATACTTCTATAGTTGTACCTGTAGATACACATATTAGATTCGTTGTAACAGCTGCTGATGTTATTCATGATTTTGCTATCCCAAGTTTAGGTATTAAAGTTGATGCTACTCCTGGTAGATTAAATCAAGTTTCTGCTTTAATTCAAAGAGAAGGTGTCTTCTATGGGGCATGTTCTGAGTTGTGTGGGACAGGTCATGCAAATATGCCAATTAAGATCGAAGCAGTATCATTACCTAAATTTTTGGAATGATTAAATGAACAATAATTAATATTTACTTATTATTAATATTTTTAATTATTAAAAATAATAATAATAATAATAATTATAATAATATTCTTAAATATAATAAAGATATAGATTTATATTCTATTCAATCACCTTATATTAAAAATATAAATATTATTAAAAGAGGTTATCATACTTCTTTAAATAATAAATTAATTATTGGTTCAAAAGGATAATAAAAATAATAATAAGAATAACCCAGAATTGATAATTTTTATAAATGATTAGTGGATTTACAGATGGAGATGGTAGTTTTTATATTAAATTAAATGATAAAAAATATTTAAGATTTTTTTATGGTTTTAGAATACATATTGATGATAAAGCATGTTTAGAAAAGATTAGAAATATATTAAATATACCTTCTAATTTTGAAGAACTACTTAAAACAATTATATTAGTAAATTCACAAAAGAAATGGTTATATTCTAATATTGTAACTATTTTTGATAAGTATCCTTGTTTAACAATTAAATATTATAGTTATTATAAATGAAAAATAGCTATAATTAATAATTTAAATGGTATATCTTATAATAATAAAGATTTATTAAATATTAAAAATACAATTAATAATTATGAAGTTATACCTAATTTAAAAATTCCATATGATAAAATAAATGATTATTGAATTTTAGGTTTTATTGAAGCTGAAGGTTCATTTGATCTATCTCCAAAACGTAATATTTGTGGTTTTAATGTTTCACAACATAAACGTAGTATTAATACATTAAAAGCTATTAAATCTTATGTATTAAATAATTGAAAACCAATTGATAATACACCATTATTAATTAAAAATAAATTATTAAAAGATTGAGATTCATCTATTAAATTAACTAAACCTGATAAAAATGGAGTTATTAAATTAGAATTTAATAGAATAGATTTTTTATATTATGTTATTTTACCTAAATTATATTCATTAAAATGATATAGTCGTAAAGAAATTGATTTCCAATTATGAAAAACACTTATAGAAATCTATATAAAAGGTTTACATAATACACTTAAAGGTTCTAATTTATTAAAATTAATTAATAATAATATTAATAAAAAAAGATATTATTCTAATTATAATATTTCTCCTTTCGGGGTTCCGGCTCCCGTGGCCGGGCCCCGATGATAAGCTGTCAAACATGAGAATTAATTC

sgRNA expression

SNR52-sgRNA-IM83

Blue: SNR52 promoter; Teal background: sgRNA targeting ARG8^m^; yellow background: IM83

TtctttgaaaagataatgtatgattatgctttcactcatatttatacagaaacttgatgttttctttcgagtatatacaaggtgattacatgtacgtttgaagtacaactctagattttgtagtgccctcttgggctagcggtaaaggtgcgcattttttcacaccctacaatgttctgttcaaaagattttggtcaaacgctgtagaagtgaaagttggtgcgcatgtttcggcgttcgaaacttctccgcagtgaaagataaatgatcagcTTAgtgatgtaagatttgttttagagctagaaatagcaagttaaaataaggctagtccgttatcaacttgaaaAAGTGGCACCGAGTCGGTGGATCcGCCTTGTTAGCTCAGTTGGTAGAGCCCCTATGAGGCTCC

SNR52-sgRNA-msIM83

Blue: SNR52 promoter; Teal background: sgRNA targeting ARG8^m^; yellow background: IM83

TtctttgaaaagataatgtatgattatgctttcactcatatttatacagaaacttgatgttttctttcgagtatatacaaggtgattacatgtacgtttgaagtacaactctagattttgtagtgccctcttgggctagcggtaaaggtgcgcattttttcacaccctacaatgttctgttcaaaagattttggtcaaacgctgtagaagtgaaagttggtgcgcatgtttcggcgttcgaaacttctccgcagtgaaagataaatgatcagcTTAgtgatgtaagatttGTTTTAGAGCTAGAAATAGCAAGTTAAAATAAGGCTAGTCCGTTATCAACTTGAAAAAGTGGCACCGGCCTTGTTAGCTCAGTTGGTAGAGCCCCTATGAGGCTCCACGGTGAAGC

Other IM sequences

| No | Sequence |
| --- | --- |
| 1 | GGTTCGAGCCCCCTACAGGGCTCCA |
| 2 | GGTTCGAGGGCCCTACAGGCCCTCCA |
| 3 | GGTTCGAGGGCCCTACAGCCCTCCA |
| 4 | GAACCTAAAAAAAATAGGTTCGAGCCCCCTACAGGGCTCCA |
| 5 | GCCTTGTTGGCGCAATCGGTAGCGCAAATACAGGGCTCCA |
| 6 | GCCTTGTTGGCGCAATCGGTAGCGCACCCCCACCCCCTACAGGGCTCC |
| 7 | GCCTTGTGCGCAATCGGTAGCGCCCTACAGGGCTCCA |
| 8 | GCCTTGTTAGCTCAGTTGGTAGAGCCCCTATGAGGCTCCA |
| 9 | GCGCAATCGGTAGCGCTTCGAGCCCCCTACAGGGCTCCA |
| 10 | GCTCAGTTGGTAGGAGCTTCGAGCCCCCTATGAGGCTCCA |
| 11 | GCCTTGTTGGCGCAATCGGTTAGCGCGTATGACTCTTAATCATAAGGTTAGGGGTTCGAGCCCCCTACAGGGCTCCA |
| 12 | TCCTTGTTAGGCTCAGTTGGTAGAGCGTTCGGCTTTTAACCGAAATGTCAGGGGTTCGAGCCCCCTATGAGGAGCCA |
| 13 | TCCTTGTTAGGCTCAGTTGGTAGAGCGTTCGGCTCTTAACCGAAATGTCAGGGGTTCGAGCCCCCTATGAGGAGCCA |
| 14 | GCCTTGTTGGCGCAATCGGTTAGCGCGTATGACTTTTAATCATAAGGTTAGGGGTTCGAGCCCCCTACAGGGCTCCA |
| 15 | GAGAATATTGTTTAATGGTAAAACAGTTGTCTTTTAAGCAACCCATGCTTGGTTCAACTCCAGCTATTCTCACCA |
| 16 | GACTGTAAAGCTAACTTAGCATTAACCTTTTAAGTTAAAGATTAAGAGAACCAACACCTCTTTACAGTCACCA |
| 17 | GTCTACGACCATACCACGTTGAAAACACCGGTTCTCGTCCGATCACCGAAGTTAAGCAACGTAGGGCCTGCCCAGTACTTGGATGGGTGACCGCCTGGGAACAGCAGGTGTTGTAGACTT |
| 18 | GTCTCCCTGAGCTTCAGGGAG |
| 19 | TTGGCGAAACCCCATTTCGACCTTTCGGTCTCATCAGGGGTGGCACACACCACCCTATGGGGAGAGGTCGTCCTCTATCTCTCCTGGAAGGCCGGAGCAATCCAAAAGAGGTACACCCACCCATGGGTCGGGACTTTAAATTCGGAGGATTCGTCCTTTAAACGTTCCTCCAAGAGTCCCTTCCCCAAACCCTTACTTTGTAAGTGTGGTTCGGCGAATGTACCGTTTCGTCCTTTCGGACTCATCAGGGAAAGTACACACTTTCCGACGGTGGGTTCGTCGACACCTCTCCCCCTCCCAGGTACTATCCCCTTTCCAGGATTTGTTCCC |
| 20 | AGGTACACCCACCCATGGGTCGGGACTTTAAATTCGGAGGATTCGTCCTTTAAACGTTCCTCCAAGAGTCCCTTCCCCAAACCCTTACTTTGTAAGTGTGGTTCGGCGAA |
| 21 | GGGTTGCGGAGGGTGGGCCTGGGAGGGGTGGTGGCCATTTTTTGTCTAACCCTAACTGAGAAGGGCGTAGGCGCCGTGCTTTTGCTCCCCGCGCGCTGTTTTTCTCGCTGACTTTCAGCGGGCGGAAAAGCCTCGGCCTGCCGCCTTCCACCGTTCATTCTAGAGCAAACAAAAAATGTCAGCTGCTGGCCCGTTCGCCCCTCCCGGGGACCTGCGGCGGGTCGCCTGCCCAGCCCCCGAACCCCGCCTGGAGGCCGCGGTCGGCCCGGGGCTTCTCCGGAGGCACCCACTGCCACCGCGAAGAGTTGGGCTCTGTCAGCCGCGGGTCTCTCGGGGGCGAGGGCGAGGTTCAGGCCTTTCAGGCCGCAGGAAGAGGAACGGAGCGAGTCCCCGCGCGCGGCGCGATTCCCTGAGCTGTGGGACGTGCACCCAGGACTCGGCTCACACATGC |
| 22 | GGCGTAGGCGCCGTGCTTTTGCTCCCCGCGCGCTGTTTTTCTCGCTGACTTTCAGCGGGCGGAAAAGCCTCGGCCTGCCGCCTTCCACCGTTCATTCTAGAGCAAACAAAAAATGTCAGCTGCTGGCCCGTTCGCCCCTCCCGGGGACCTGCGGCGGGTCGCCTGCCCAGCCCCCGAACCCCGCCTGGAGGCCGCGGTCGGCCCGGGGCTTCTCCGGAGGCACCCACTGCCACCGCGAAGAGTTGGGCTCTGTCAGCCGCGGGTCTCTCGGGGGCGAGGGCGAGGTTCAGGCCTTTCAGGCCGCAGGAAGAGGAACGGAGCGAGTCCCCGCGCGCGGCGCGATTCCCTGAGCTGTGGGACGTGCACCCAGGACTCGGCTCACACATGC |
| 23 | GGGACAGTAGCTCAATTGGTAGAGCAT |
| 24 | GGGACAGTAGCTGAATTGGTACAGCAT |
| 25 | AGAAGCGTATCCCGCTGAGC |
| 26 | TCTCCCTGAGCTTCAGGGAG |
| 27 | GGCCTGGTTAGTACTTGGATGGGAGACCGCCAAGGAATACCGGGTG |
| 28 | GCGCAATCGGTAGCGC |
| 29 | GAGCCCCCTACAGGGCTC |
| 30 | GCGCAATCGGTAGCGCGAGCCCCCTACAGGGCTC |
